# Rational design of intrinsic transcription terminators with programmable strength

**DOI:** 10.64898/2026.01.22.701115

**Authors:** Guilherme E. Kundlatsch, Almiro P. S. Neto, Gabriela B. de Paiva, Elibio Rech, Leonardo T. Duarte, Danielle B. Pedrolli

## Abstract

Intrinsic terminators are widely used in synthetic biology and metabolic engineering, yet their strength must be selected empirically from characterized libraries. We therefore developed a framework for designing intrinsic terminators with programmable strength. TK, designed to maximize termination strength, exceeded both B0010, a widely used standard, and the strongest terminator in a 582-member reference library when assayed in parallel. To establish that these gains reflect intrinsic termination rather than cellular context, we developed a cloning-free *in vitro* transcription assay using fluorescent light-up aptamers; TK again ranked highest. The framework improved the canonical T7 terminator and recovered strength rankings across three other bacterial species. Designs were generated using an interpretable sequence-to-function model trained from primary-sequence descriptors alone. The framework is available as open web tools, TerSP for strength prediction and TerFac for target-driven design. Model interpretation revealed previously unreported determinants, including a preference for compositionally diverse hairpin stems.

## INTRODUCTION

Intrinsic terminators are the default parts for transcriptional termination in synthetic biology and metabolic engineering, valued for their simplicity, genetic compactness, and portability across host contexts [1]. Nevertheless, the property that matters most to a designer, how strongly a given terminator will terminate, still cannot be specified in advance. Terminators act by destabilizing the elongation complex (EC), a highly stable intermediate in the transcription cycle, causing RNA polymerase to dissociate from the DNA template and the nascent RNA [2]. They fall into two mechanistic classes: factor-dependent terminators, which recruit trans-acting proteins such as Rho and Mfd to induce EC collapse [3,4], and intrinsic terminators, which encode termination entirely within their own nucleotide sequence and secondary structure [5]. This autonomy (operating without dedicated protein factors) is precisely what makes intrinsic terminators portable between hosts, while their unpredictability is a recurring obstacle to reliable genetic design.

This limitation has direct consequences for gene-circuit performance, as accurate and efficient transcriptional termination is critical for modularity and predictability. Incomplete termination can result in transcriptional read-through into downstream elements, leading to aberrant gene expression, circuit malfunction, and increased metabolic burden. For instance, substituting a leaky T7 terminator with a tandem array of three intrinsic terminators increased volumetric protein yield in *E. coli* by 1.6-fold, an improvement attributed to enhanced termination efficiency and reduced resource competition [6]. In *Bacillus subtilis*, adding a terminator increased gene expression by 20-fold compared to a terminator-less construct [7].

As the scale and complexity of synthetic constructs increase, the demand for high-efficiency terminators becomes increasingly stringent. This is particularly evident in systems composed of densely packed transcriptional units, strong promoters, and multigene operons. In such contexts, high RNA polymerase flux across short intergenic regions necessitates precise transcriptional termination to prevent transcriptional interference. Additionally, engineering multigene constructs requires sequence-diverse terminators with predefined strengths to facilitate DNA synthesis and reduce the risk of homologous recombination associated with repeated use of identical terminator elements [8].

Despite the growing engineering demand and their relative simplicity, intrinsic terminators remain difficult to design predictably. To date, we have been unable to rationally engineer them to display a desired level of transcription termination or even precisely predict the termination efficiency of a given terminator. Early computational approaches used features such as hairpin stability and the downstream U-tract to identify intrinsic terminators, but primarily addressed whether a terminator was present rather than how strongly it terminated [9–15]. Later sequence-to-function and biophysical models enabled quantitative prediction, but their performance remained limited [16]. Chen *et al.* [8], for example, characterized 582 natural and synthetic *E. coli* terminators and derived a biophysical model that explained 40% of the variance in strength after excluding a subset of weak terminators. More revealing than this predictive ceiling, however, was their failure in design: combining individually favorable features did not reliably produce stronger terminators across scaffolds, showing that features associated with strength do not necessarily provide rules for engineering it. More recent machine-learning approaches have distinguished strong from weak terminators, but categorical outputs neither resolve quantitative differences in strength nor provide the continuous objective required for target-driven design [17]. Thus, despite progress from terminator identification to quantitative modelling, the gap between characterizing terminators and engineering them to a specified strength remains. A detailed comparison of previous computational approaches to terminator modelling is provided in Table S1.

Here we set out to make terminator strength a design specification rather than a property discovered after construction. We report synthetic terminators generated by inverse design and validate them both in living cells and in a purified transcription system, where the strongest designs exceed established reference terminators. To obtain a measure of termination uncoupled from translation, we developed an *in vitro* fluorescence assay based on fluorescent light-up aptamers (FLAPs), which removes the sequence-dependent translational effects that confound protein-reporter measurements [18]. The designs were produced by inverting an interpretable sequence-to-function model trained on the 582-terminator dataset of Chen *et al.* [8]. We release the model, together with its inverse-design routine, as two open tools, the Terminator Strength Predictor (TerSP) and the Terminator Factory (TerFac), which respectively predict and generate terminators of specified strength.

## RESULTS

### Global sequence metrics provide informative, low-dimensional features that limit overfitting risk

Using the 582 intrinsic terminators characterized by Chen *et al.*, we evaluated more than 130 sequence features across the A-tract, hairpin, loop, and U-tract (Figure 1a and Table S2). Feature selection retained 16 descriptors for model training (Figure S1). Within the A-tract, local composition immediately upstream of the hairpin was informative, with adenine and cytosine content in the final six nucleotides retained over equivalent composition descriptors calculated across alternative windows, consistent with the region emphasized by Chen *et al.* Meanwhile, the U-tract was represented by informative features spanning different windows. Uracil content across the first six and ten nucleotides was retained, as was cytosine content across the full U-tract. However, since cytosine occurrence was unevenly represented in the dataset in both tracts, these apparent preferences may reflect dataset composition rather than distinct functional effects (Table S3). For the hairpin, the length of the initial contiguous GC stretch was more informative than GC content within the first three stem positions or across the hairpin as a whole. This supports the importance of the GC-rich hairpin base reported by Chen *et al.,* while indicating that the informative region can extend beyond the first three positions. A more detailed comparison of the selected features with previous findings in the literature is provided in Table S4.

**Figure 1.**
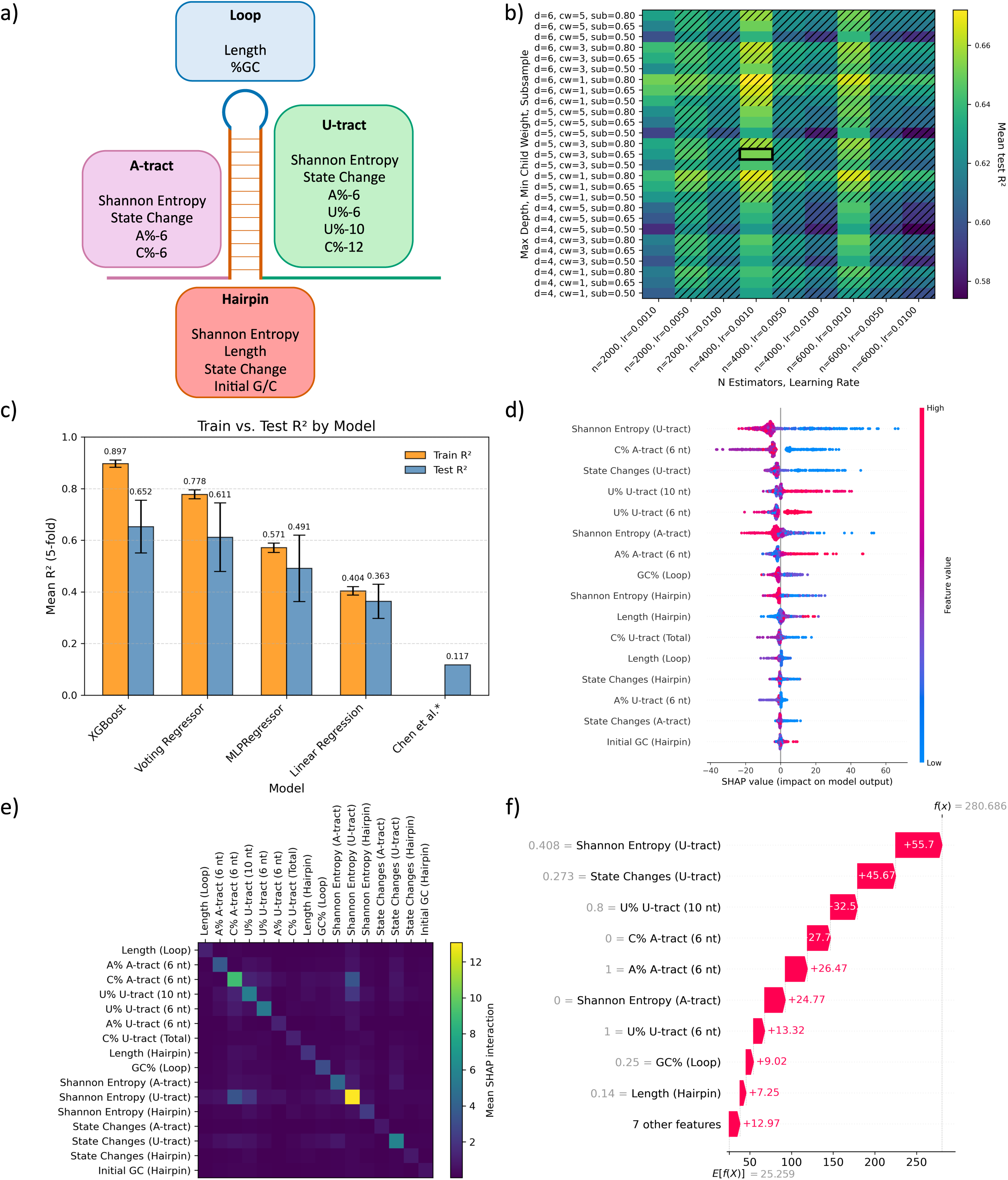
Feature engineering, model selection, and SHAP interpretation of the terminator-strength predictor. (a) Schematic representation of the terminator regions used for feature extraction and the final selected features assigned to each region. (b) XGBoost hyperparameter optimization. Heatmap values indicate mean held-out R² across cross-validation folds. Configurations with a training–test R² gap greater than 0.3 or training R² greater than 0.9 were excluded from model selection and are shown as hatched cells. The optimal configuration is indicated by a black rectangle (n_estimators = 4000, learning_rate = 0.001, max_depth = 5, min_child_weight = 3, and subsample = 0.65). (c) Mean training and held-out R² values for the evaluated models, compared with the kinetic model reported by Chen *et al*. [8]. Data are shown as mean ± SD across five cross-validation folds. (d) SHAP global summary plot showing the contribution of each feature to model predictions. Each point represents one terminator, with color indicating the corresponding feature value. (e) SHAP interaction heatmap showing mean pairwise interaction values among selected features. (f) SHAP waterfall plot for a representative high-strength terminator, showing how individual feature values shift the prediction from the model baseline to the final predicted strength. See also Tables S2-S4 and Figures S1-S6.

Rather than relying on predicted ΔG or other structural properties, we prioritized sequence-derived descriptors and found that two global metrics, Shannon entropy and state-change count, were themselves informative predictors of terminator strength. Shannon entropy captures compositional heterogeneity, whereas state-change count captures the local arrangement of nucleotides. Their retention among the selected features shows that broad sequence organization can be represented by a small number of interpretable descriptors, rather than through many highly localized variables. This compact representation limits feature-space dimensionality while preserving biologically meaningful information, as further reflected in their contributions to model predictions below. A similar sequence-first strategy has also outperformed predefined structural inputs in previous machine-learning models of RNA switches [19].

### The model outperforms the previous kinetic model and effectively captures distinct biological signals

Following grid-search optimization, the best-performing XGBoost configuration achieved an average training R² of 0.897 and test R² of 0.652 under five-fold cross-validation (Figure 1b and Figure S2), substantially outperforming the kinetic model reported by Chen et al. (R² = 0.117; Figure S3). A linear regression baseline performed considerably worse (R² = 0.363), indicating that the relationship between the selected sequence features and terminator strength was not adequately captured by a linear model (Figure 1c and Figure S4). We also evaluated neural-network and ensemble approaches, but neither improved on standalone XGBoost performance (Figure 1c and Figures S5 and S6). The optimized XGBoost model was therefore selected for downstream analyses.

SHAP analysis showed that A-tract and U-tract descriptors contributed more strongly to model predictions than hairpin-related features (Figure 1d). Within the U-tract, low Shannon entropy and low state-change count were associated with higher predicted strength, consistent with stronger termination from homogeneous poly-U regions with few interruptions. Uracil content extending to position 10 also contributed substantially, supporting the broader U-rich signal identified during feature selection (Table S4). In a representative high-strength terminator, most of the increase above the model mean was attributed to U-tract features, with smaller contributions from A-tract descriptors (Figure 1e). Finally, SHAP interaction values were predominantly diagonal, with limited off-diagonal intensity, indicating that most selected features contributed largely independently to model output (Figure 1f). This limited interaction structure supports the use of the selected descriptors as distinct, interpretable sequence signals.

### Forward-engineered terminators efficiently terminate transcription in *E. coli*

To assess the model’s generative capacity, we designed terminators while constraining the A– and U-tracts to six adenines and eight uracils, respectively, allowing the model to explore new hairpins. This produced TK, a 72-nt terminator with a predicted strength of 275. We then restricted hairpin size to generate miniTK, a compact 44-nt variant predicted at 255. For comparison, the highest model-predicted strength among the original library was 238.

To experimentally evaluate the model-designed terminators, we characterized their activity in *E. coli* using the same dual-reporter assay used by Chen *et al.* (Figure 2a) alongside B0010, a widely used strong terminator, and Tmax, the strongest terminator in the original dataset (Figure 2b and Table S5). TK showed the highest activity among all terminators tested *in vivo*, reaching a strength of 475.1 ± 6.8 and exceeding both B0010 (357.4 ± 17.3) and Tmax (342.2 ± 11.2). MiniTK also functioned as a strong terminator, with a measured strength of 201.2 ± 7.2, placing it within the top 4.3% of the Chen library. Thus, TK outperformed both a widely used reference and the strongest terminator from the training dataset under the same assay framework, supporting the model’s ability to guide the design of high-efficiency terminators.

**Figure 2.**
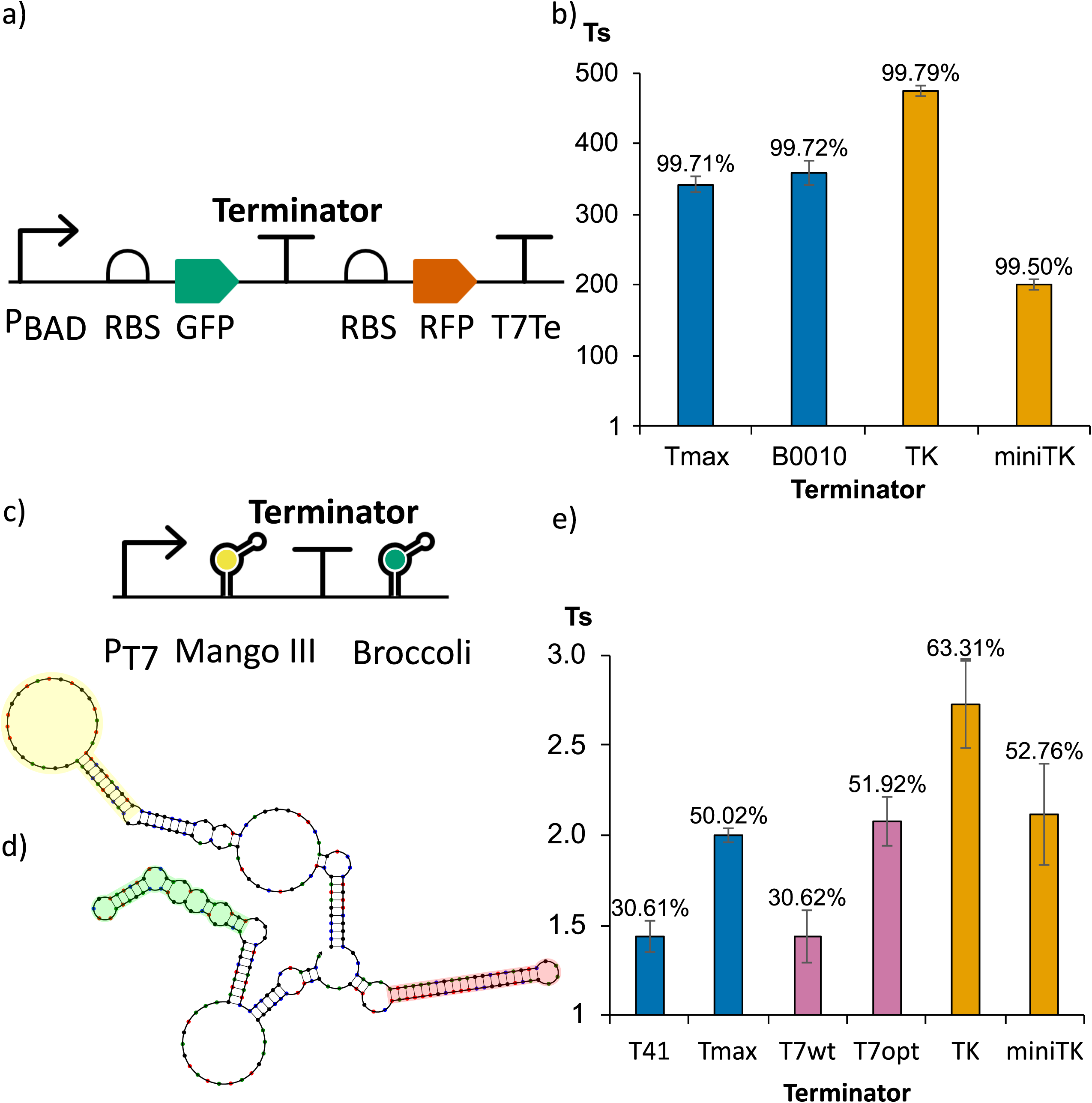
Measurement of terminator strength *in vivo* and *in vitro*. (a) Schematic representation of the dual-reporter construct used for *in vivo* terminator characterization in *E. coli*. Terminators were inserted between *GFP* and *RFP* coding sequences. RBS, ribosome-binding site; T7Te, T7 terminator. (b) *In vivo* terminator strength (Ts) of reference and model-designed terminators, calculated from GFP and RFP fluorescence using the normalized upstream/downstream reporter ratio described by Chen *et al.* Bars show mean Ts, and error bars indicate SD. (c) Schematic representation of the DNA template used for the dual-FLAP *in vitro* transcription assay. The terminator was positioned between Mango III, used as the upstream transcription reference, and Broccoli, used as the downstream readthrough reporter. (d) Predicted RNA secondary structure of the dual-FLAP transcript containing Tmax, showing insulated Mango III and Broccoli aptamers flanking the terminator region. (e) *In vitro* terminator strength for reference terminators, T7wt, T7opt, and model-designed terminators measured using the dual-FLAP assay. Bars show mean Ts and error bars indicate SD. The equation shown defines the calculation of Ts, where U and D denote upstream and downstream reporter signals, respectively, and 0 denotes the no-terminator control. For the *in vivo* assay, U = GFP and D = RFP; for the *in vitro* assay, U = Mango III and D = Broccoli. See also Tables S6 and S7 and Figures S7-S11.

Although the *in vivo* assay demonstrated that the model-designed terminators function efficiently in *E. coli*, these measurements reflect termination within a cellular context in which transcription and translation are coupled and can be influenced by variations in DNA copy number, mRNA stability, and accessory proteins such as NusA, NusG, and Rho. Since the training dataset was generated entirely *in vivo*, we asked whether the learned sequence features reflected intrinsic termination properties independent of these cellular effects. We therefore developed an *in vitro* termination assay based on FLAPs [20]

### Newly designed *in vitro* assay isolates transcription and directly assesses termination function

Adapting the dual-reporter strategy to an *in vitro* transcription format required RNA reporters compatible with real-time monitoring with minimal spectral interference. We tested Corn, Broccoli, Baby Spinach, Mango II, and Mango III, individually and in combination, and selected Broccoli-BI and Mango III-TO1-biotin for dual-channel monitoring. Reporter-only controls showed no Broccoli interference in the Mango III channel, while limited Mango III spillover into the Broccoli channel was quantified and corrected (Figure S7). We also compared hybrid and double-stranded DNA templates; the hybrid format showed delayed signal accumulation and approximately fourfold lower Broccoli fluorescence, so double-stranded templates were used thereafter (Figure S8).

We next validated whether the Mango III–terminator–Broccoli architecture could simultaneously report upstream transcription and downstream readthrough. Using T1, a terminator expected to permit extensive readthrough, fluorescence accumulated in both channels, confirming detection across the terminator region (Figure S9). Since the downstream reporter must remain sufficiently sensitive to resolve strong terminators, we also tested the inverse Broccoli–terminator–Mango III orientation using T41, the weakest strong-classified terminator in the Chen dataset. Although upstream Broccoli fluorescence was robust, downstream Mango III fluorescence remained close to background and highly variable (Figure S10). Because even T41 reduced the readthrough signal to near-background levels, this configuration was unsuitable for resolving strong terminators. We therefore selected Mango III–terminator–Broccoli as the final reporter configuration on a double-stranded DNA template (Figure 2c).

Finally, we optimized rNTP concentration. At the 5 mM rNTP concentration typical of commercial high-yield *in vitro* transcription systems, we observed substantial downstream Broccoli fluorescence even for Tmax, indicating high readthrough-associated signal. Reducing the concentration to 0.3 mM markedly decreased this signal, enabling clear discrimination among T1, T41, and Tmax (Figure S11). This effect has been described previously [21] and is consistent with the kinetic competition between terminator-mediated transcript release and continued transcription elongation.

During assay development, a central concern was that perturbation of FLAP secondary structure, particularly in the downstream reporter, could artificially increase the calculated terminator strength. If Broccoli folding was impaired, fluorescence would decrease independently of termination and could be misinterpreted as reduced readthrough. Accordingly, we flanked both FLAPs with RNA insulators [22] and used NUPACK [23] to simulate the secondary structure of transcripts containing each tested terminator. All constructs were predicted to preserve the Broccoli folding without structural interference between the terminator and reporter modules (Figure 2d). The expected ranking of weak, threshold-strength, and strong terminators *in vitro* (Figure S11) supports the functionality of this approach.

### Forward-engineered terminators outperform the strongest dataset terminator *in vitro*

Using the optimized dual-FLAP assay, we compared TK and miniTK with T41 and Tmax, benchmarks near the weak/strong threshold and at the top of the Chen dataset, respectively. Tmax showed higher termination than T41 (2.00 ± 0.03 versus 1.44 ± 0.09), confirming that the assay distinguished terminators of different strengths. As observed *in vivo*, TK produced the highest score, reaching 2.72 ± 0.24. Surprisingly, miniTK also performed strongly, reaching 2.12 ± 0.28, comparable to Tmax despite its lower relative activity *in vivo* (Figure 2e and Table S6).

We also evaluated the canonical T7 terminator, which has known incomplete termination [6,24,25], comparing the wild-type sequence (T7wt) with an optimized variant (T7opt). T7wt showed intermediate termination (1.44 ± 0.14), similar to T41, consistent with its previously reported suboptimal termination capacity. T7opt retained the native hairpin and loop but replaced the A– and U-tracts with model-selected sequences, increasing the score to 2.08 ± 0.14, comparable to Tmax and miniTK. This agreed with model predictions, which placed T7wt slightly below T41 and T7opt substantially above it (Table S7). Notably, this agreement extended to a viral terminator despite the absence of viral sequences from the training dataset.

### Terminator Strength Predictor, a tool to predict strength from sequence in *E. coli* and beyond

The optimized XGBoost model was implemented as the Terminator Strength Predictor (TerSP), which validates user-provided sequences, computes the 16 model features, and returns predicted strength, efficiency, and weak/strong classification using the ≥40 threshold defined by Chen *et al..* Since the surrogate was trained exclusively on terminators characterized in *E. coli*, its primary intended use was to estimate how synthetic or heterologous sequences might perform in an *E. coli*-based context. Nevertheless, we asked whether the learned feature-to-strength mapping could recover experimental trends across species. As an exploratory test of cross-species transferability, we excluded training-set terminators and applied TerSP to datasets from three non-model bacteria.

TerSP transferred remarkably well to six natural *Corynebacterium glutamicum* [26] terminators, showing strong ranking agreement (Kendall’s τb = 0.87), correctly ordering the four strongest variants and correctly classifying all six as weak (Table 1 and Figure 3). Substantial ranking agreement was also retained for *Vibrio natriegens* [27] (τb = 0.50) and *Halomonas bluephagenesis* [28] (τb = 0.41). TerSP correctly identified B1006 as the only strong *V. natriegens* terminator and classified all eight sequences correctly. For *H. bluephagenesis*, 11 of 14 terminators were correctly classified, with A02 and A01 accounting for the largest ranking discrepancies. Full predicted and experimental values are provided in Table S8.

**Figure 3.**
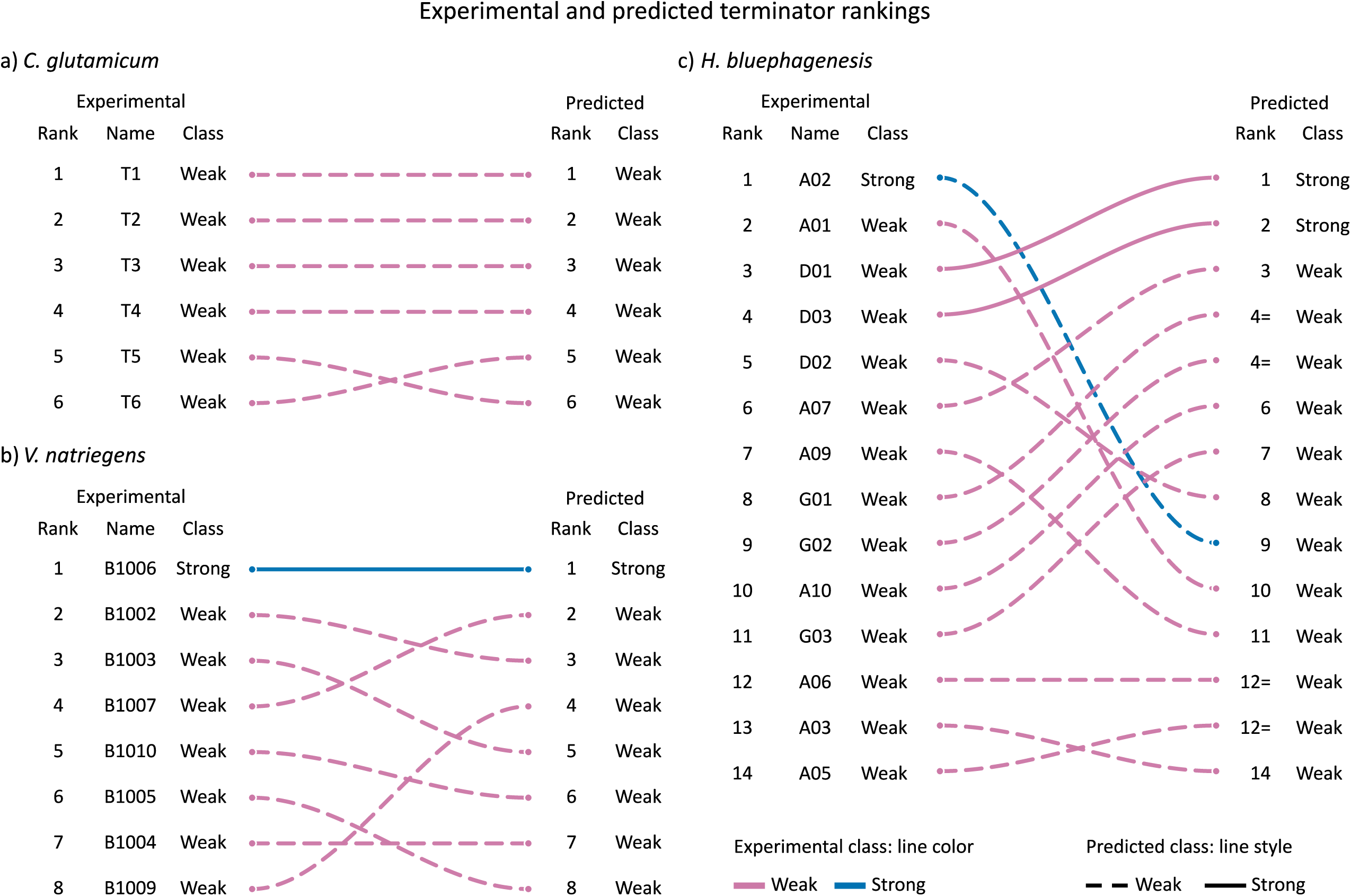
TerSP strength prediction beyond *E. coli*. Experimental and TerSP-predicted terminator rankings for (a) *C. glutamicum*, (b) *V. natriegens*, and (c) *H. bluephagenesis*. For each terminator, a connector links its experimentally reported rank and class on the left to its TerSP-predicted rank and class on the right. Rank numbers increase from top to bottom, with the strongest experimental terminator ranked as 1. Connector color indicates the experimental classification, with blue representing strong terminators and magenta representing weak terminators. Line style indicates the predicted classification, with solid lines representing strong predictions and dashed lines representing weak predictions. Predicted rank ties are indicated by an equal sign; tied terminators were vertically ordered according to their experimental ranking for visualization. Terminators reported in the original studies that were included in the TerSP training dataset were excluded from the analysis. See also Table S8.

**Table 1.** Evaluation of TerSP performance across external terminator datasets. Ranking agreement was assessed using Kendall’s τ_b_ and the proportion of concordant within-dataset pairs. Classification accuracy represents the percentage of terminators correctly assigned as weak or strong.

| Dataset | Kendall's $\tau_b$ | Concordant pairs | Concordance (%) | Classification accuracy (%) |
| --- | --- | --- | --- | --- |
| *C. glutamicum* | 0.87 | 14/15 | 93.3 | 100 |
| *V. natriegens* | 0.50 | 21/28 | 75.0 | 100 |
| *H. bluephagenesis* | 0.41 | 63/91 | 69.2 | 78.6 |
| Pooled total | 0.48 | 98/134 | 73.1 | 89.3 |

Across the three external datasets, TerSP demonstrated moderate overall agreement with experimental rankings while maintaining high classification performance despite differences in reporters and plasmid architectures among studies (Table 1), demonstrating useful predictive power beyond the *E. coli* training dataset. To our knowledge, TerSP is the first non-kinetic tool to provide sequence-based quantitative strength prediction for natural and synthetic intrinsic terminators. By moving beyond terminator identification toward quantitative strength prediction, TerSP emerges as a promising tool for exploring transcriptional regulation in non-model organisms, particularly where organism-specific datasets are not yet available to train context-specific models.

### Terminator Factory, a tool to design user-specified strength terminators with sequence diversity

Next, we developed Terminator Factory (TerFac), an inverse-design tool built on the same feature-based surrogate model to generate terminators for a user-defined target strength. Considering that arbitrary combinations of features may not correspond to biologically realizable sequences, we constructed a discrete feasibility map of valid combinations for each terminator region. The complete A-tract, U-tract, and loop spaces were exhaustively enumerated, whereas the much larger hairpin space was mapped using composition-constrained enumeration and adaptive Monte Carlo sampling. Hairpin coverage saturated after ∼2.33 × 10^8^ iterations and matched exhaustive enumeration for L = 6–24, supporting completeness. TerFac searches this discrete, non-convex space by optimizing candidate feature vectors and snapping continuous candidates to the nearest valid feature combinations. It generates sequences that satisfy the requested structural constraints and returns their predicted strengths and corresponding features. We also used the software to generate a list of maximum-strength terminators for each hairpin size (Table S9). Interestingly, the model frequently favored increased state changes within the hairpin to achieve higher strength values, a trend not previously reported in the literature. Together, these results establish a unified framework from feature discovery and model training to experimental validation, strength prediction, and inverse design, summarized in Figure S12.

## Discussion

Terminator strength can now be specified in advance rather than discovered by testing. The designed terminator TK exceeded both B0010, a widely used standard, and Tmax, the strongest terminator in the training library, and this advantage persisted in a purified *in vitro* assay, indicating that it reflects intrinsic termination rather than cellular context. These results demonstrate that the model can be used prospectively to guide the design of terminators. Its interpretability provides an additional advantage, allowing the predictive features to be examined as sequence determinants of termination.

The selected features both recapitulated established terminator biology and refined previously defined sequence windows, while also revealing previously undescribed descriptors, particularly within the hairpin (Table S4). The final model achieved a mean held-out R² of 0.652 under five-fold cross-validation, which can be considered modest for many applications. However, this performance is informative in the context of the dataset and the intended use of the model for several reasons. In addition to being relatively small, the dataset is strongly imbalanced, with high-strength terminators sparsely represented (Table S3). Moreover, the numerical strength scale amplifies differences among highly efficient terminators: strengths of 100 and 300 correspond to termination efficiencies of 99.0% and 99.67%, respectively. Thus, numerically large errors in strength can correspond to much smaller functional differences. Classification analysis further showed that the model retained substantial ability to identify strong terminators despite the marked class imbalance (Table S3).

During the preparation of this manuscript, Li *et al.* [29] published a model derived from the same dataset. Their computational pipeline differed markedly from ours: k-mer, CKSNAP [30], and PseKNC [31] descriptors increased feature-space dimensionality by two to three orders of magnitude, and thermodynamic and secondary-structure metrics that we intentionally omitted were incorporated. Although their reported test R² (0.7204) surpassed ours, their generative performance did not: the 18 synthetic terminators produced did not exhibit termination efficiencies significantly higher than their reference B0010. Moreover, none of the hairpin-related insights identified here emerged in their analysis, directly reflecting differences in training design. These differences illustrate that predictive accuracy alone does not fully capture the usefulness of a model for biological design. For small and constrained datasets, evaluation should therefore consider not only held-out predictive performance but also interpretability, designability, and experimental validation of generated sequences.

Here, experimental validation was extended beyond the established *in vivo* method to a purified transcription system using a new dual-FLAP assay designed to minimize the influence of cellular context. The assay confirmed TK as the strongest intrinsic terminator. In addition to validating individual designs, the assay provides a scalable methodology for investigating transcriptional regulation and generating datasets tailored to *in vitro* systems. This is particularly relevant as increasingly complex *in vitro* circuits require biological parts characterized for those conditions. Although optimizing the reaction conditions required several iterative refinements, the finalized protocol enables high-throughput evaluation of a large number of sequences with minimal equipment and time. Compared to the workflow established by Chen *et al.* [8], our method eliminates cloning, reducing experimental time while facilitating automation and improving reproducibility.

Finally, we developed two free, open-source tools, TerSP and TerFac, both using the model as a surrogate for biological experimentation. TerSP implements the forward use of this surrogate, enabling prediction of terminator strength from user-provided sequences, and was developed primarily for an *E. coli*-based context. However, its meaningful, though still imperfect, success in ranking terminators from other bacteria suggests that the model captured sequence determinants that extend beyond the training organism. This represents a quantitative extension of a principle already established by terminator-identification tools: intrinsic terminators can often be recognized across bacterial species because some of their core structural features are broadly conserved. Classical tools such as TransTermHP [15] and RNIE [32], as well as newer machine-learning approaches such as BacTermFinder [33] and TerminatorNet [34], have demonstrated that terminator occurrence can be predicted across diverse bacterial genomes, including organisms not used during model development. TerSP builds on this foundation by moving from qualitative identification to relative strength prediction. Although organism-specific datasets could produce more accurate context-aware models, such datasets are unavailable for most bacteria, making TerSP a practical first-pass tool for estimating the relative strength of uncharacterized natural or synthetic terminators in non-model organisms and for prioritizing candidates for experimental validation.

Whereas TerSP supports the analysis of existing terminators, TerFac extends the surrogate toward terminator design. A user-defined target strength guides the optimization of feasible feature profiles, which are then reconstructed into candidate sequences. Experimental validation of TK and miniTK provides an initial proof-of-principle that this surrogate-guided strategy can generate functional synthetic terminators with high activity. Beyond the design of strong terminators, TerFac could also be used to generate intermediate-strength sequences, potentially enabling controlled readthrough to tune downstream gene expression [35]. In addition, TerFac can generate distinct sequences with similar predicted strengths, creating diversity that minimizes sequence repetition and lowers recombination risk in large or multipart constructs.

To further interrogate the design rules captured by the model, we used TerFac to generate maximum-strength terminators across different hairpin lengths. These designs exhibited features that had not emerged as dominant in the global analysis; we interpret them as modifiers that distinguish between otherwise strong terminators. A stronger interaction at the hairpin base seems to become progressively more necessary as the hairpin stem lengthens, with the initial G-C count reaching nine pairs in the longest design. Moreover, the six uracils immediately downstream of the hairpin were invariably conserved. Although U content through position +10 ranked highly in the feature analysis, maximum-strength designs converged on eight uracils in this window and a U-tract state-change count of three, yielding uninterrupted proximal runs of six to eight uracils. This agrees with previous reports that the first eight positions are most critical, with distal uracils contributing more modestly to strength. Another pattern concerned the sequence organization of the hairpin. Maximum-strength terminators consistently displayed high Shannon entropy and state-change counts, indicating that the model favored compositionally diverse stems with frequent changes in nucleotide identity. We propose two complementary hypotheses for this preference. First, greater sequence variation and fewer consecutive identical nucleotides may favor the intended stem by reducing alternative base-pairing and competing structures. Repetitive RNA sequences are known to accommodate slipped duplex conformations [36,37], and previous work has established that sequence choice can strongly influence ensemble defect [38]. Although the model itself does not directly measure ensemble defect, its preference for high state-change may therefore reflect sequence properties associated with greater pairing specificity. Second, nucleotide order can alter the physical properties of the stem even when base-pair composition is unchanged. For example, GG/CC, GC/CG, and CG/GC RNA nearest-neighbor steps have ΔG°_37_ values of approximately −3.28, −3.46, and −2.33 kcal mol^−1^, respectively [39], demonstrating that rearranging the same G-C pairs generates distinct stacking environments. Consistently, structural analysis of alternating CG RNA duplexes revealed sequence-dependent differences in stacking geometry and helical parameters [40,41]. Such differences could influence intrinsic termination because hairpin formation occurs cotranscriptionally under structural constraints imposed by RNA polymerase [42]. Although these mechanisms are consistent with established principles of RNA folding and structure, their contribution to terminator strength remains to be directly tested and will be explored in future applications of TerFac. Notably, these properties emerged despite the deliberate exclusion of predicted free-energy and secondary-structure descriptors from the model, suggesting that sequence-level features may have captured information related to both structural specificity and folding behavior directly from the primary sequence.

### Concluding remarks

In this work, we present an *in vitro* dual-FLAP assay for terminator characterization and two computational tools that extend terminator engineering from sequence-based strength prediction to target-driven design. Together, these resources expand the synthetic-biology toolkit for measuring, predicting, and designing transcription terminators, supporting the engineering of increasingly complex genetic systems. These resources address two questions recently raised [43]. The dual-FLAP assay shows that fluorescent RNA aptamers can report transcription flux directly, while TerSP and TerFac show that AI can predict terminator efficiency and design new ones with specified strength. The approaches are complementary: that study identifies natural terminators genome-wide, whereas ours generates new-to-nature sequences. Moving from designed parts to designed function raises a further set of questions, from the mechanistic basis of the stem compositions the model favors to the amount of host-specific data required for organism-specific design (see Outstanding Questions).

### Outstanding Questions

What exactly needs to be relearned when transferring a quantitative terminator model between hosts? Can a small, model-guided calibration set enable host-specific recalibration, and how does the amount of required data change with phylogenetic distance?

When designing an entire transcriptional unit, can quantitative models for promoters, RBSs, and terminators be combined modularly, or must interactions among these regulatory elements be learned jointly? Could whole-unit co-design outperform independent part optimization?

When quantitative datasets are limited, can compact interpretable models provide more reliable design rules than large pretrained models? Can low-dimensional models abstract general principles that then guide high-capacity models in exploring and optimizing promising regions of design space?

### Technology readiness

TerSP and TerFac establish a framework that moves terminator engineering from characterization toward specification: TerSP predicts strength from sequence, whereas TerFac generates sequences meeting a user-defined strength target. Both are freely accessible through a browser, and the model and code are openly available. Designed terminators were experimentally validated in *E. coli* and in a newly developed *in vitro* assay, including a sequence that outperformed established benchmarks, while TerSP retained useful ranking power across datasets from three additional bacteria. Together, these results place the core prediction and design technologies at Technology Readiness Level (TRL) 4: validated under laboratory conditions. Their most consequential applications, however, remain to be demonstrated. Advancing toward TRL 5 will require showing that designed terminators improve system-level performance in relevant settings. Two particularly promising applications are the use of intermediate-strength terminators to tune transcriptional or metabolic flux through multigene pathways, and the generation of multiple sequence-diverse strong terminators that retain matched function while reducing recombination in large constructs. Broader prospective validation is also needed to establish TerSP portability across hosts, genetic contexts, and transcription systems and to determine when host-specific recalibration is required. Ultimately, integration with promoter and RBS design could further extend this approach toward joint specification of complete transcriptional units.

## DATA AVAILABILITY

All datasets, model artifacts, and custom code underlying this study are publicly available at https://github.com/gkundlatsch/TerSP-TerFac. This repository includes the datasets used for model development and evaluation, derived feature tables, feature-mapping data, benchmarking analyses, hyperparameter-optimization outputs, trained models, and the scripts used to reproduce the computational analyses.

The TerSP web application and associated prediction model are available at https://github.com/gkundlatsch/TerSP-Web. TerFac is available as both an online version (https://github.com/gkundlatsch/TerFac) and a standalone offline version (https://github.com/gkundlatsch/TerFac-offline). TerSP and TerFac can also be accessed directly through a web browser at https://taaqui.unesp.br/a3lf1.

## SUPPLEMENTARY DATA STATEMENT

Supplementary data are provided as a supplementary Word file containing supplementary tables, supplementary figures, and additional methods.

## AUTHOR CONTRIBUTION

G.E.K., A.P.S.N., and G.B.P. designed and performed the experiments. G.E.K. constructed the models and analyzed the data. E.R., L.T.D., and D.B.P. conceived and supervised the project. D.B.P. acquired funding. G.E.K. and D.B.P. wrote the manuscript. All authors revised the final manuscript.

## METHODS

### EXPERIMENTAL MODEL AND STUDY PARTICIPANT DETAILS

#### Escherichia coli TOP10

*E. coli* TOP10 was used for plasmid cloning and in vivo terminator-strength assays. Cells were maintained in LB medium at 37°C. Strains carrying the pGR dual-reporter plasmid or its derivatives were cultured in the presence of 100 μg mL⁻¹ ampicillin. Chemically competent TOP10 cells used for plasmid transformation were prepared in-house.

## METHOD DETAILS

### Dataset and feature engineering

The model was developed using the dataset of 582 characterized intrinsic terminators reported by Chen *et al.* [8]. The reported Average Strength values were used directly as the regression target. All 582 terminators were retained, without filtering, outlier removal, transformation, or target normalization. Synthetic and natural terminators were combined into a single dataset, and 130 candidate sequence-derived descriptors were generated across the four regions defined by Chen *et al.*: A-tract, hairpin, loop, and U-tract. The complete list and definitions of candidate descriptors are provided in Table S2.

For the 8-nt A-tract, nucleotide composition was evaluated as the fractions of A, C, G, U, and GC across the complete region and across windows of one to seven nucleotides extending upstream from the hairpin. Sequence diversity was additionally represented by Shannon entropy, calculated with base 2 from the nucleotide-frequency distribution of the region [44]. Shannon entropy was calculated from nucleotide frequencies as 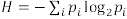, where *p_i_* is the frequency of nucleotide *i* ɛ {*A*, *C*, *G*, *U*} within the analyzed sequence region. State-change count was defined in this study as the number of adjacent nucleotide pairs with different nucleotide identities, 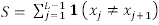, where *L* is sequence length, and **1** is the indicator function. Adjacent-nucleotide transition/transversion descriptors and positional nucleotide-scoring descriptors were also explored. For the latter, alternative weighting schemes emphasizing positions closer to the hairpin were manually evaluated as part of the broader candidate-feature exploration.

Hairpin descriptors were calculated both from the complete hairpin sequence and from the loop-excluded sequence corresponding to the paired region. Candidate features included the number of nucleotides in the loop-excluded hairpin, nucleotide and GC composition of the paired region, Shannon entropy, and state-change count. A composite stability descriptor was calculated as the loop-excluded hairpin length multiplied by its GC fraction. GC content was additionally evaluated across the first one, two, and three nucleotides of the hairpin. The initial hairpin GC count was defined as the number of consecutive G or C nucleotides from the beginning of the hairpin sequence before the first A or U was encountered. Adjacent-nucleotide transition/transversion descriptors were also evaluated for the hairpin. Loop descriptors were treated separately from those of the paired hairpin region and included loop length, A, C, G, U, and GC fractions, and Shannon entropy.

For the 12-nt U-tract, A, C, G, U, and GC fractions were calculated across the complete region and across windows of one to eleven nucleotides extending downstream from the hairpin. Shannon entropy and state-change count were calculated as described above, and adjacent-nucleotide transition/transversion and positional scoring descriptors were also evaluated.

For adjacent-nucleotide transition/transversion descriptors, AG/GA and CU/UC pairs were classified as transitions, whereas AC/CA, AU/UA, GC/CG, and GU/UG were classified as transversions. The descriptor was calculated as the number of transitions divided by the number of transversions; when no transversions were present, the transition count itself was returned.

Feature design intentionally emphasized descriptors derived directly from the primary nucleotide sequence rather than predicted thermodynamic or structural attributes such as free energy (Δ*G*). We considered these predicted properties to be derived from the underlying sequence and therefore did not include them as candidate predictors. Instead, descriptors such as Shannon entropy and state-change count were used to capture broader sequence properties within each terminator region while limiting feature-space dimensionality. This strategy was intended to control model complexity and reduce the risk of overfitting during supervised learning [45]. A similar sequence-centered approach has been reported for machine-learning models of RNA switches, in which models trained directly on sequence information outperformed models incorporating predefined structural inputs [19].

Feature engineering was performed in Python 3.11.11 using pandas v2.2.2, NumPy v1.26.4, SciPy v1.13.1, and scikit-learn v1.3.2. Derived descriptors requiring normalization were scaled to the [0,1] range using MinMaxScaler. Scaling parameters were determined using the complete set of 582 terminators before division into training and evaluation subsets. Raw sequence fields and intermediate variables used only for descriptor calculation were excluded from the modelling matrix.

### Feature selection

Feature selection was performed manually using a forward-selection strategy. The complete dataset was divided into fixed training (80%) and evaluation (20%) subsets using random_state=1996, and the same partition was maintained throughout feature evaluation. Candidate descriptors were iteratively incorporated into an XGBoost regression model, and changes in training and evaluation R² were inspected after each addition. Descriptors associated with gains in predictive performance were retained, resulting in a final set of 16 features for subsequent model optimization. The sequential incorporation of the retained descriptors and the corresponding changes in training and evaluation R² are shown in Figure S1.

### Hyperparameter optimization

Following feature selection, XGBoost hyperparameters were optimized using the fixed 16-feature set. The normalized dataset was divided into training (80%) and held-out test (20%) subsets using train_test_split with test_size=0.2 and random_state=1996. Hyperparameter optimization was performed on the training subset using GridSearchCV with R² as the scoring metric and five-fold cross-validation. The optimization evaluated n_estimators (2,000, 4,000, 6,000, 8,000, and 10,000), learning_rate (0.001, 0.0025, 0.005, 0.0075, and 0.01), max_depth (2–6), min_child_weight (1, 3, and 5), and subsample (0.50, 0.65, and 0.80), using XGBRegressor(objective=’reg:squarederror’, random_state=1996). Mean training and cross-validation R² values were retained for every parameter combination.

The complete hyperparameter landscape was visualized as a heatmap of mean cross-validation R², organized by max_depth, min_child_weight, and subsample versus n_estimators and learning_rate (Figure S2). To reduce selection of strongly overfit configurations, combinations were flagged when the mean training-validation R² gap exceeded 0.3 or when mean training R² exceeded 0.9. These thresholds were used as conservative empirical safeguards during model selection rather than as general performance criteria. Parameter combinations satisfying either criterion were excluded from selection, and the remaining configurations were ranked according to mean cross-validation R². The selected configuration comprised 4,000 estimators, a learning rate of 0.001, maximum depth of 5, minimum child weight of 3, and a subsampling fraction of 0.65. The separate ranking procedure calculated the training-validation R² difference, removed configurations with a gap of 0.3 or greater, and ranked the remaining solutions by validation R².

### Alternative model evaluation

For comparison with a previously established quantitative model, predictions were also obtained using the biophysical model reported by Chen *et al.* The model combines energetic contributions from the U-tract, hairpin loop, stem base, and A-tract according to the equation and fitted parameters defined in the original study. They reported an R² of 0.40 from the measured-versus-predicted comparison shown on logarithmic axes in their Fig. 4a. For consistency with the regression metrics reported here, we additionally calculated R² between the predicted and experimental values on the untransformed Average Strength scale, obtaining R² = 0.117 (Figure S3).

We additionally investigated three alternative modeling approaches: linear regression, multilayer perceptron neural networks, and XGBoost–MLP VotingRegressor ensembles. Linear regression was assessed using five-fold cross-validation. In each fold, the model was fitted on four folds, and predictions were generated for both the training data and the held-out validation fold. R² was calculated for each fold, and mean training and validation R² values were calculated across the five folds. Experimental and predicted terminator strengths for each validation fold are shown in Figure S4. Linear regression achieved mean training and validation R² values of 0.404 and 0.363, respectively.

MLP performance was evaluated across nine candidate hidden-layer architectures using five-fold cross-validation (Figure S5). The tested architectures were 10, 50, 10:10, 10:20:10, 20:50:20, 50:100:50, 100:100, 100:200:100, and 50:100:100:50, where each number indicates the number of neurons in the corresponding hidden layer. For each architecture and fold, an MLPRegressor was trained with ReLU activation, the Adam optimizer, random_state=1996, max_iter=20000, and tol=1 × 10−4. Predictions were generated for both the training and validation folds, and R² and mean squared error (MSE) were calculated. Mean and standard deviation of each metric were then calculated across the five folds for each architecture. Although the four-layer 50:100:100:50 architecture produced the highest mean validation R², the 10:20:10 architecture was selected for subsequent ensemble evaluation because it provided similar cross-validated performance with lower model complexity and a smaller training–validation R² discrepancy (Figure S5).

The selected MLP architecture was subsequently evaluated in weighted ensembles with the optimized XGBoost model using VotingRegressor. Relative XGBoost:MLP weights of 0:1, 1:0, 1:1, 2:1, 1:2, 5:1, and 10:1 were tested. Each weighting scheme was evaluated by five-fold cross-validation, with R² and MSE calculated independently for the training and validation folds and summarized as the mean and standard deviation across folds. The resulting models were compared according to mean validation R² (Figure S6). XGBoost alone achieved the highest mean validation R² (0.652), whereas incorporating the MLP predictions did not improve cross-validated performance. XGBoost was therefore retained as the final regression model.

### Final model training

Following model comparison, the optimized XGBoost regressor was trained on the complete 582-terminator dataset using the 16 selected sequence-derived features and the hyperparameters identified above. The trained model was serialized using joblib.dump for subsequent deployment in the prediction and design tools.

### Model interpretability using SHAP

Model interpretability was assessed using SHAP (SHapley Additive exPlanations) [46] v0.47.2 within a five-fold cross-validation framework. In each fold, the XGBoost model was fitted on the training folds, and a shap.TreeExplainer was instantiated from the fitted model. SHAP values (shap_values) and pairwise SHAP interaction values (shap_interaction_values) were then calculated exclusively for the held-out validation fold. The SHAP values, interaction matrices, and corresponding feature values obtained from each fold were concatenated, producing out-of-fold SHAP estimates in which each of the 582 terminators was represented once.

The aggregated out-of-fold SHAP values were used to characterize feature contributions across the complete dataset. A SHAP summary (beeswarm) plot was generated to visualize the magnitude and direction of each feature’s contribution across terminators. Pairwise feature interactions were summarized by calculating, for each feature pair, the mean absolute SHAP interaction value across all held-out observations; these values were displayed as an interaction heatmap, with larger values indicating greater pairwise contributions to model predictions. Finally, a representative terminator was selected as the held-out observation with the largest sum of absolute SHAP values across all features. A waterfall plot was generated for this observation to decompose its prediction into the individual feature contributions relative to the model’s expected value.

### Terminator Strength Predictor implementation

The final trained XGBoost model was implemented as the Terminator Strength Predictor (TerSP) in both offline and web-based formats (Figure S12). TerSP represents the forward use of the trained model as a surrogate for experimental terminator characterization: a user-provided sequence is mapped to its predicted strength.

The offline implementation was developed as a standalone Python script that loads the serialized model using joblib.load. Input terminators are provided as an A-tract, two hairpin half-stems, a loop, and a U-tract. Sequences are converted to uppercase RNA notation (T to U) and validated according to the sequence-region constraints used by TerSP: A-tract, 8 nt; U-tract, 12 nt; each half-stem, 3–24 nt; and loop, 3–16 nt. The two half-stems are concatenated to reconstruct the loop-excluded hairpin sequence, and the same 16 sequence-derived descriptors used for model training are calculated and normalized to [0,1] using the normalization scheme applied during model development. The resulting feature vector is passed to the trained XGBoost model to obtain the predicted Average Strength. For batch prediction, input sequences are read from an Excel file and the raw feature values, normalized feature values, and predictions are exported to an output Excel file.

The web-based implementation was developed in Python using Flask v2.2.5 and applies the same model-loading, sequence-validation, feature-calculation, normalization, and prediction pipeline as the offline implementation. Predicted terminator strength (*T_S_*) is converted to termination efficiency (*TE*) as 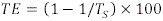. Predictions are classified as strong when *T_S_* ≥ 40 and weak otherwise. The web-based tool is available at: https://taaqui.unesp.br/a3lf1

### TerSP cross-species evaluation

Cross-species transferability of TerSP was evaluated using terminators characterized independently in *Corynebacterium glutamicum*, *Vibrio natriegens*, and *Halomonas bluephagenesis* by Kang *et al.* [26], Stukenberg *et al.* [27], and Xu *et al.* [28], respectively. No species-specific retraining or recalibration of TerSP was performed before prediction. Terminators present in the 582-sequence model-training dataset were excluded from the external evaluation.

Six *C. glutamicum* terminators were retained for analysis. Terminator strength was measured using an eGFP-terminator-mRFP1 dual-reporter construct implemented in the RoboMoClo vector system, with calculated according to the approach of Chen *et al.* The six terminators were named in decreasing order of experimentally measured strength, with T1 the strongest and T6 the weakest. All had *T_S_* < 10 and were described by the authors as relatively intermediate-strength terminators. For *V. natriegens*, B0010 was excluded because it was present in the TerSP training dataset, and B0015 was excluded because it is a double terminator, leaving eight terminators for analysis. Since exact numerical values were not reported, their experimental rank order was inferred from Figure 6D of the original study. Stukenberg *et al.* measured terminator activity using the ratio between downstream luminescence and upstream mScarlet-I fluorescence, with lower ratios indicating stronger termination. For *H. bluephagenesis*, Xu *et al.* quantified termination efficiency using an sfGFP-terminator-mRFP dual-reporter system. Four terminators with negative experimentally reported termination efficiencies were excluded. The synthetic terminator G04 was additionally excluded because its U-tract contained eight nucleotides, below the 12-nucleotide U-tract length required by TerSP, leaving 14 terminators for analysis. Experimental termination-efficiency values were used for the retained terminators (Table S8).

Taking in account that the three source studies used different experimental assays, the cross-species comparison was based primarily on within-dataset rank order and experimentally defined weak/strong classes rather than direct comparison of absolute activities. Agreement between predicted and experimental rankings was quantified separately for each dataset using Kendall’s τ_b_ [47,48]. For each dataset, all possible terminator pairs were classified as concordant, discordant, or tied. A pair was considered concordant when the predicted and experimental rankings ordered the two terminators in the same direction, and Kendall’s τ_b_ incorporated correction for ties in either ranking. Pairwise concordance was additionally calculated as the number of concordant pairs divided by the total number of possible within-dataset pairs, 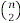; tied pairs therefore remained part of the denominator.

For the pooled summary, concordant, discordant, and tied pair counts were summed across the three datasets before calculation of Kendall’s τ_b_ and pairwise concordance. Cross-dataset pairs were not considered because experimental rankings restart independently within each dataset and the underlying assays and measurement scales are not directly comparable. The resulting dataset-specific and pooled rank statistics are reported in Table 1, and the complete terminator-level predictions and experimental information are provided in Table S8.

Terminators were also evaluated by weak/strong classification. TerSP predictions were classified as strong when *T_S_* ≥ 40 and weak otherwise. For the external datasets, experimental weak/strong classes were assigned by the original authors. Classification accuracy was calculated as the proportion of terminators for which the TerSP-predicted class agreed with the experimentally assigned class.

### Terminator Factory implementation

In contrast to the forward-prediction workflow implemented in TerSP, Terminator Factory (TerFac) uses the trained XGBoost model as an inverse-design surrogate. Starting from a user-defined target terminator strength, TerFac searches the model feature space for feasible combinations predicted to produce the requested strength and subsequently reconstructs those feature combinations as nucleotide sequences. Since arbitrary combinations of the 16 model features do not necessarily correspond to physically realizable sequences, discrete feasibility maps were first constructed independently for the A-tract, U-tract, loop, and hairpin stem regions.

For the loop, all combinations of loop length *L* = 3-16 nt and GC count from 0 to *L* were represented, normalized according to the model feature ranges, and deduplicated into unique feature pairs. For the fixed-length A-tract and U-tract regions, the complete sequence spaces were exhaustively enumerated. All 4^8^ = 65,536 A-tract sequences and 4^12^ = 16,777,216 U-tract sequences were generated, their corresponding model features calculated and normalized, and duplicate feature combinations removed.

Direct exhaustive enumeration became prohibitive for longer hairpin stems. In TerFac, designed stems were restricted to perfect reverse-complement pairing between the two stem halves and therefore to even total loop-excluded lengths *L* = 6, 8, …, 48 nt. For each *L*, the first-half length was ℓ= *L* / 2. Instead of enumerating every possible first-half nucleotide sequence, we first grouped sequences by nucleotide composition, defined only by the numbers of A, C, G, and U residues present. Thus, sequences with different nucleotide orders but the same counts, for example ACGU and UGCA, belong to the same composition. All compositions satisfying *a* + *c* + *g* + *u* = ℓ were enumerated. The number of distinct compositions is 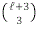, whereas the number of possible ordered first-half sequences is 4^ℓ^. At the maximum stem length, *L* = 48nt (ℓ = 24), this corresponds to only 2,925 nucleotide compositions compared with approximately 2.81 × 10^14^ possible ordered first-half sequences.

Once the stem length and nucleotide composition of the first half were defined, two of the four hairpin descriptors used by the model could be determined directly: loop-excluded hairpin length was fixed by *L*, and Shannon entropy was determined by the nucleotide composition of the resulting stem. In contrast, state-change count and initial GC count depend on the order in which those nucleotides occur and therefore cannot be inferred from composition alone. To map these ordering-dependent dimensions of the feasible feature space, alternative arrangements were generated for each nucleotide composition. Three deterministic arrangements were first evaluated: a sorted arrangement, its reverse-sorted counterpart, and a high-transition jagged arrangement. Each first-half sequence was joined to its reverse complement to construct the complete loop-excluded stem, after which the four hairpin descriptors were calculated and normalized using the feature bounds applied by the model.

Intermediate feature combinations were populated by adaptive Monte Carlo sampling. For each nucleotide composition, random permutations of the corresponding nucleotide multiset were generated in batches of 5,000. Newly observed normalized feature tuples were added to the mapping table, and sampling of a composition was terminated after three consecutive batches produced no new feature combinations. The cumulative number of sampled sequences and unique feature combinations was recorded at intervals of 10,000 draws. Approximately 2.33 ×10^8^ permutations were evaluated during construction of the complete hairpin feature map.

To assess coverage of the sampling procedure, the resulting feature spaces were compared with complete sequence enumeration for stem lengths *L* = 6–24. For each of these lengths, all possible first-half sequences under the reverse-complement constraint were enumerated and converted to normalized feature tuples. The adaptive mapping procedure recovered the same set of unique feature combinations as exhaustive enumeration over this range, supporting saturation of the sampled feature space.

### TerFac target-strength optimization

TerFac searches the resulting discrete feasibility maps using differential evolution (DE) implemented with scipy.optimize.differential_evolution in SciPy v1.13.1. Candidate solutions are represented initially as continuous 16-dimensional vectors bounded between 0 and 1 and are converted to feasible feature combinations through a hierarchical snapping procedure before model evaluation. For the loop, the nearest valid normalized loop length is selected first, followed by the nearest permissible GC value. For the A-tract and U-tract, the nearest entropy value is selected, after which the remaining region-specific features are matched to the closest valid tuple by Euclidean distance. For the hairpin, stem length is selected first, followed by the valid combination of initial GC count, entropy, and state-change count minimizing Euclidean distance.

For target-strength design, the objective function was defined as the absolute difference between model-predicted and user-requested strength,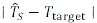. The user can additionally specify a maximum hairpin stem length, an acceptable deviation from the target strength, a hairpin entropy tolerance, and a random seed. DE was run using the best1bin strategy with maxiter=1000, popsize=15, and tol=1\times10^{–6}. Optimization terminates early when a candidate reaches the user-defined acceptable deviation.

The selected feature vector is subsequently reconstructed into nucleotide sequences. A-tract sequences are identified by exhaustive enumeration of the 8-nt sequence space. U-tract reconstruction first filter nucleotide compositions according to the required entropy and achievable state-change range and then enumerate unique multiset permutations until all selected positional-composition constraints are satisfied, using an entropy tolerance of 0.01 in the deployed implementation. Hairpin reconstruction similarly filters first-half nucleotide compositions according to entropy, achievable state-change count, and sufficient G/C content for the required initial GC run; candidate arrangements are then evaluated using the user-specified hairpin entropy tolerance, with a default of 0.1 in the web implementation. The second stem half is generated as the reverse complement of the selected first half. A loop sequence satisfying the selected length and GC-content constraints is then generated.

The four regions are assembled in the order A-tract, first half-stem, loop, second half-stem, U-tract. The complete set of model features is recalculated directly from the assembled sequence, and its final predicted strength is obtained from this reconstructed sequence rather than from the intermediate optimized feature vector.

### *De novo* terminator design

For the experimentally tested designs, the A– and U-tracts were constrained to contain six adenines and eight uracils, respectively, allowing optimization to focus on the hairpin. This choice was motivated by the feature analysis, in which the model-selected A– and U-tract descriptors largely agreed with previously described sequence preferences, whereas the selected hairpin descriptors showed the greatest divergence from previous descriptions of intrinsic terminators (Table S4). Under these constraints, TerFac was used to generate TK. To obtain a more compact design and reduce potential difficulties associated with synthesis and cloning of long terminator sequences, the allowable hairpin length was subsequently restricted, producing miniTK.

### Maximum-strength analysis

Separately from the target-strength TerFac workflow, the same feasibility maps and surrogate model were used to identify maximum-strength designs for each even hairpin stem length from 6 to 48 nt. For each fixed stem length, DE maximized predicted strength by minimizing its negative value. Five independent runs were performed using random seeds 0, 13, 96, 555, and 1000, and the highest-scoring solution was retained. These searches used the best1bin strategy, maxiter=3000, popsize=100, tol=1\times10^{–5}, Latin-hypercube initialization, mutation factors from 0.7 to 1.1, and a recombination rate of 0.6. The resulting maximum-strength designs are reported in Table S9.

### TerFac web implementation

The online TerFac implementation was developed as a Flask-based web application. Users provide a target strength and may additionally specify maximum hairpin length, acceptable target deviation, hairpin entropy tolerance, and random seed. The deployed interface uses default values of 24 nt for maximum hairpin stem length, 1 strength unit for acceptable deviation, 0.1 for hairpin entropy tolerance, and 1 for the random seed when these parameters are not supplied. Optimization jobs are executed asynchronously using Redis and RQ, with a one-hour execution limit. Upon completion, the application returns the assembled A-tract, hairpin halves, loop, U-tract, and complete terminator sequence together with the reconstructed feature values and final model-predicted strength. The complete computational workflow linking model development, feasible-feature mapping, TerSP, and TerFac is summarized in Figure S12.

All computational analyses were performed on a workstation equipped with an Intel Core i7-10700KF CPU (3.80 GHz), 64 GB of RAM, and an NVIDIA GeForce GT 730 GPU with 4 GB of memory. On this hardware, model training required only a few minutes, and execution of TerSP was typically completed within a few seconds per sequence. TerFac was more computationally demanding, with execution time increasing with hairpin length and target strength, ranging from a few minutes for shorter designs to several hours for more demanding searches. Maximum-strength searches were the most computationally intensive and required several hours per hairpin length in some cases.

### Plasmid construction

The terminators were cloned into the pGR plasmid [8] between *the eGFP* and *mRFP1* reporter genes using the *Eco*RI and *Spe*I restriction sites. Terminator inserts were ordered as oligos for both forward and reverse strands, with compatible sticky ends built into the oligos that matched the restriction sites after digestion. The forward and reverse oligos were resuspended in TE buffer and mixed to form an equimolar 100 nM solution. The mixture was heated at 98°C for 5 min and slowly cooled down to 10°C for annealing. Terminator inserts were ligated to the digested plasmid backbone using T4 DNA ligase. The ligation products were used to transform in-house prepared chemically competent *E. coli* TOP10 cells. Transformant colonies were selected from LB-agar plates containing 100 μg mL⁻¹ ampicillin and inoculated in LB medium. Overnight cultures were used to extract plasmids by miniprep. The resulting plasmids were digested with *Eco*RI and *Spe*I to select potential positive clones. The positive clones were confirmed by Sanger sequencing.

### Cell growth and induction

All cloning and assays for terminator strength were performed in *E. coli* TOP10. The transformant strains were inoculated from glycerol stocks in 4 mL Luria-Bertani medium supplemented with 100 μg mL^-1^ ampicillin in a 24-deep-well plate covered with a breathable membrane, and incubated at 37°C and 1,000 rpm in a plate incubator. After 16h, 200 µL of the pre-culture was used to inoculate fresh 4 mL LB medium supplemented with 100 μg mL^-1^ ampicillin and 10 mg mL^-1^ L-arabinose in another 24-deep-well plate, again covered with a breathable membrane. After 4h of induction at 37°C and 1,000 rpm, 15 μL of each culture was added to 185 μL Phosphate-buffered saline (PBS) for flow cytometry measurement.

### Flow cytometry

The fluorescence intensity was measured using the CytoFLEX B5-R3-V5 flow cytometer (Beckman-Coulter, California, USA). The voltage gain for each detector was set so that the full dynamic range was used for a control specimen expressing *GFP* and *RFP* without any terminators in between: FSC, 700 V; SSC, 241 V; FITC, 407 V; ECD, 650 V, at a flow rate of 30 µL/min. Compensation was measured using *E. coli* TOP10 expressing only *eGFP* or *mRFP1*. Sixty thousand events were recorded, and approximately 92% of events were retained within the cell population gate based on FSC-A and SSC-A. The fluorescence of eGFP and mRFP1 was exported as the geometric means of FITC and ECD channels, respectively. CytExpert software (v2.5.0.77) was used to analyze the data.

### Template design and preparation for *in vitro* transcription

Five FLAPs were tested as RNA reporters for in vitro transcription: Corn, Broccoli, Baby Spinach, Mango II, and Mango III. Each FLAP was positioned downstream of a T7 promoter, and the corresponding sense and antisense DNA oligonucleotides were synthesized (Table S10). Complementary oligos were annealed by mixing 10 µL of each strand with 80 µL of nuclease-free TE buffer, followed by denaturation at 98 °C and gradual cooling to room temperature on a thermal block.

To test the FLAPs in the same template separated by a terminator, both sequences were placed downstream of a T7 promoter, separated by various candidate terminators. To ensure proper folding of the aptamers and enable fluorescence emission, structural adapters were incorporated [22]. DNA templates encoding FLAP sequences, spacers, and transcription terminators were synthesized as Gene Fragments (Twist Bioscience, San Francisco, USA) (Table S11) and subsequently amplified by PCR. PCR products were purified by ethanol precipitation, and DNA concentration was assayed using the QuantiFluor® dsDNA System (Promega, Madison, USA), following the manufacturer’s instructions.

### *In vitro* transcription assay

Multiple protocols for *in vitro* transcription were evaluated. Initial assays employed the Corn aptamer in 50 µL reactions composed of 10 µL of transcription buffer (200 mM Tris-HCl, 30 mM MgCl₂, 10 mM spermidine, 50 mM NaCl), 15 µL of rNTP mix (25 mM each), 3.5 µL of T7 RNA polymerase (Promega), 50 ng of DNA template, and the appropriate fluorophore. Miniaturized 10 µL reactions were subsequently tested using 2 µL of buffer, 3 µL of rNTPs, and 0.7 µL of T7 polymerase.

For fluorescent readout, DFHO (Jena Bioscience, Jena, Germany) was added to Corn aptamer reactions; BI fluorophore (Tocris Bioscience, Bristol, UK) was used with Broccoli and Baby Spinach aptamers; and TO1-biotin (Applied Biological Materials, Richmond, Canada) was used with Mango II and Mango III. All fluorophores were solubilized in DMSO to stock concentrations of 2.5 mM (DFHO and BI) or 400 µM (TO1), and diluted in the reaction mixture to final concentrations of 60 µM (DFHO and BI) or 10 µM (TO1), respectively. Fluorescence was monitored in real time using an Infinite 200 Pro microplate reader (Tecan) in 384-well black flat-bottom microplates. Reactions were maintained at 32°C and monitored for 4 h at 3-min intervals, with orbital shaking for 150 s at a 6-mm amplitude between measurements. Mango III-TO1-biotin fluorescence was measured using excitation and emission wavelengths of 510 and 535 nm, respectively, whereas Broccoli-BI fluorescence was measured at 470 and 505 nm. Fluorescence was acquired by top reading using 25 flashes and an integration time of 20 μs.

Fluorescence output was significantly influenced by the presence of potassium and magnesium ions, necessitating the addition of extra KCl and MgCl₂ to the transcription buffer. Potassium chloride was particularly critical for proper aptamer folding, and optimal salt concentrations varied by fluorophore. RNase-free KCl and MgCl₂ stock solutions were prepared using diethyl pyrocarbonate (DEPC) treatment: 100 µL of DEPC was added per 100 mL of solution, followed by vigorous agitation, overnight incubation at 37 °C, and autoclaving for 15 minutes to inactivate residual DEPC.

Initially, we evaluated the use of fully double-stranded DNA templates versus hybrid templates in which only the T7 promoter region was double-stranded, following the protocol described by Kartje *et al.* [49]. Based on comparative fluorescence output, all subsequent transcription assays were conducted using fully double-stranded DNA templates.

To prevent accumulation of inorganic pyrophosphate, inorganic pyrophosphatase (Thermo Fisher Scientific, Waltham, USA) was added at 0.002 U/µL. After successive protocol refinements, the final optimized reaction (10 µL total volume) combining Broccoli and Mango III aptamers was assembled using 2 µL transcription buffer (200 mM Tris-HCl, 30 mM MgCl₂, 10 mM spermidine, 50 mM NaCl), 0.38 µL KCl (4 M), 0.3 µL MgCl₂ (1 M), 1 µL DTT (100 mM), 1.5 µL rNTP mix (2 mM each), 50 ng DNA template, 0.25 µL BI (2.5 mM), 0.25 µL TO1 (400 µM), 0.2 µL inorganic pyrophosphatase (0.1 U/µL), and 0.5 µL T7 RNA polymerase (80 U/µL). A commented version is available in the Supplementary Materials to increase reproducibility.

### Terminator strength

Terminator strength was calculated from the *in vivo* assay exactly as described by Chen *et al.* [8], using equation 1.

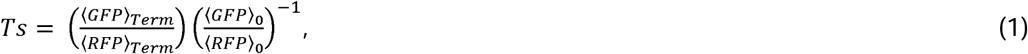

where the averages are of populations measured by flow cytometry, and the subscript *Term* refers to the measurements when the terminator is present, and 0 refers to the measurements of the control. The control is defined as a reporter construct lacking a terminator sequence.

The calculation was adjusted for the *in vitro* assay, using equation 2.

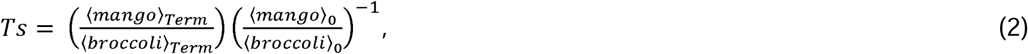

The Terminator strength (Ts) relates to the Termination efficiency (TE) through equation 3.

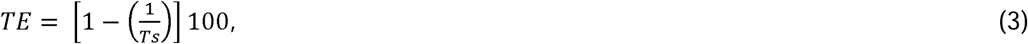

where TE is given in %.

## QUANTIFICATION AND STATISTICAL ANALYSIS

Model performance was primarily quantified using the coefficient of determination R^2^. Five-fold cross-validation was used for hyperparameter optimization and model comparison. In each iteration, four folds were used for model fitting and the remaining fold was held out for validation, such that each observation was evaluated once outside the corresponding training set. For XGBoost hyperparameter optimization, folds were generated using KFold (n_splits=5, shuffle=True, random_state=18), and mean training and validation R^2^ values were calculated across folds. The training-validation R^2^ difference was used to assess overfitting; a gap greater than 0.3 or training R^2^ greater than 0.9 was treated as a conservative empirical indicator of potential overfitting rather than as a general performance threshold. The preceding feature-selection procedure used a fixed 80:20 training/evaluation split (random_state=1996). For MLP and voting-regressor comparisons, R^2^ was calculated for each fold and summarized as mean ± SD. Linear regression was evaluated using R^2^ under the same five-fold framework. The final XGBoost model was subsequently trained on the complete dataset of 582 terminators using the selected 16 features and optimized hyperparameters.

SHAP analyses were also performed using five-fold cross-validation. For each fold, the model was trained on four folds, and SHAP values and SHAP interaction values were calculated only for the held-out fold. Results were concatenated across folds so that each terminator contributed once to the out-of-fold interpretability analysis. Pairwise interactions were summarized using the mean absolute SHAP interaction value, and the waterfall example was selected as the held-out observation with the greatest sum of absolute SHAP contributions.

Cross-species agreement between predicted and experimentally reported terminator rankings was quantified using Kendall’s τ_b_, accounting for tied ranks. Analyses were performed separately for the *C. glutamicum*, *V. natriegens*, and *H. bluephagenesis* datasets and in a pooled analysis containing only pairwise comparisons within each original dataset. Concordance was additionally reported as the proportion of experimentally ordered terminator pairs reproduced by the model. No species-specific retraining or recalibration was performed.

For the *in vivo* dual-reporter assay, eGFP and mRFP1 fluorescence were represented by the geometric mean fluorescence of the corresponding gated flow-cytometry populations. Terminator strength (*T_s_*) was calculated from the GFP:RFP fluorescence ratios relative to the terminator-free control according to the normalization procedure described above, and termination efficiency was calculated as 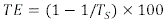. Three measurements were obtained for each tested construct, and results were summarized as mean ± SD.

For the *in vitro* dual-FLAP assay, the maximum Mango III and Broccoli fluorescence values reached during each kinetic transcription reaction were used for quantification. Their ratio was normalized to the corresponding terminator-free reference to calculate the *in vitro* termination score as described above. Four measurements were obtained for each tested terminator, and results were summarized as mean ± SD.

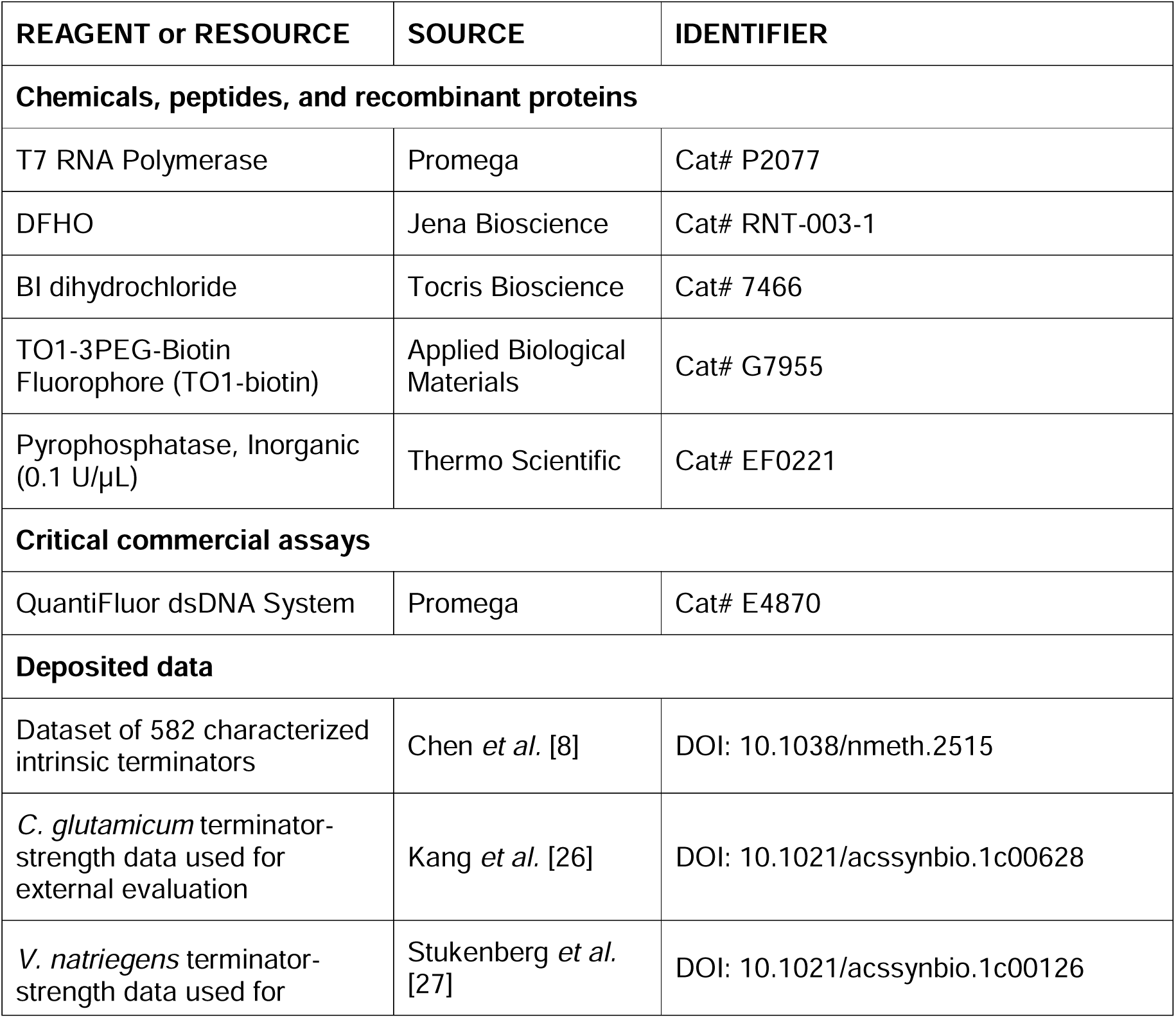
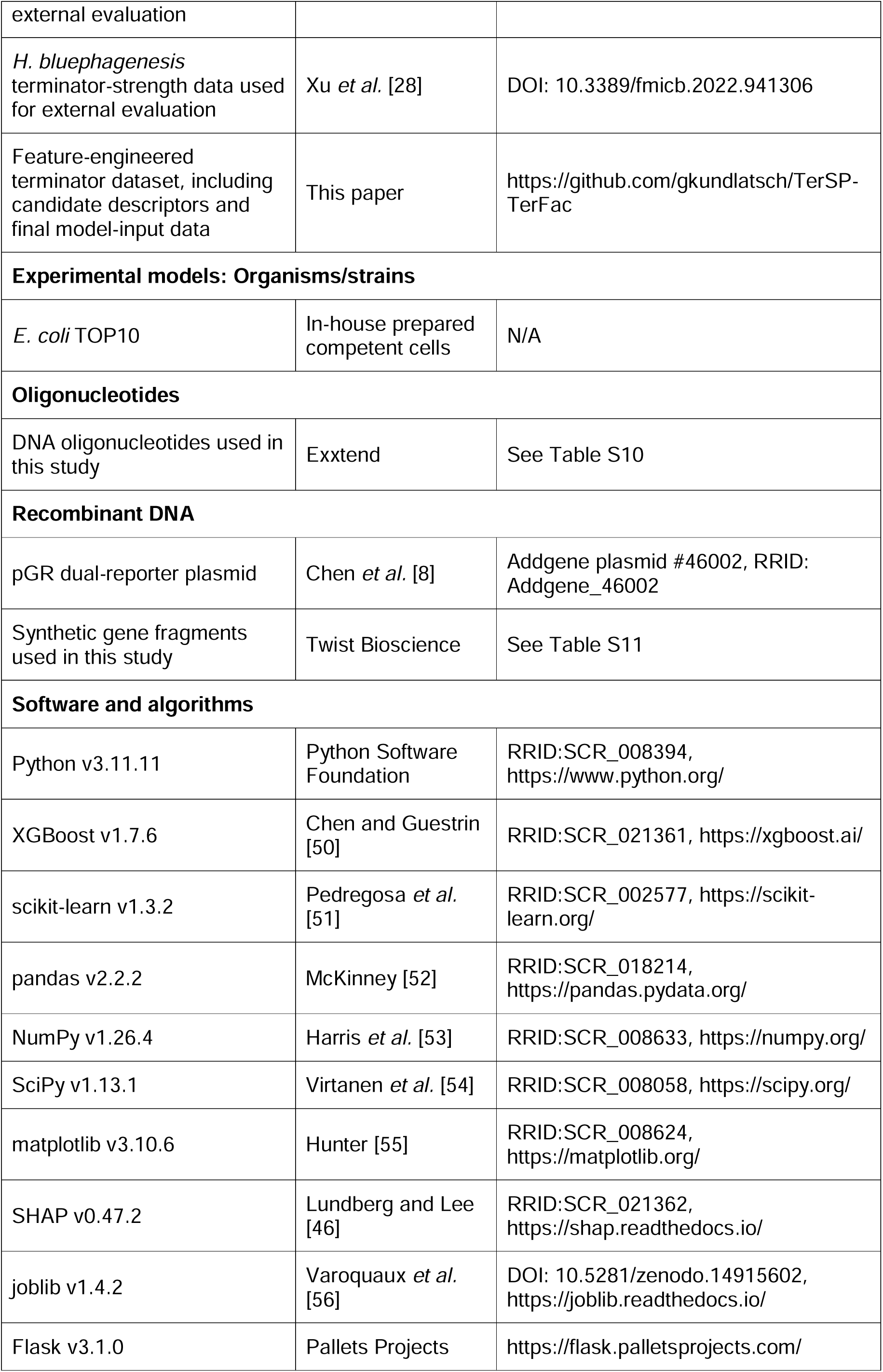
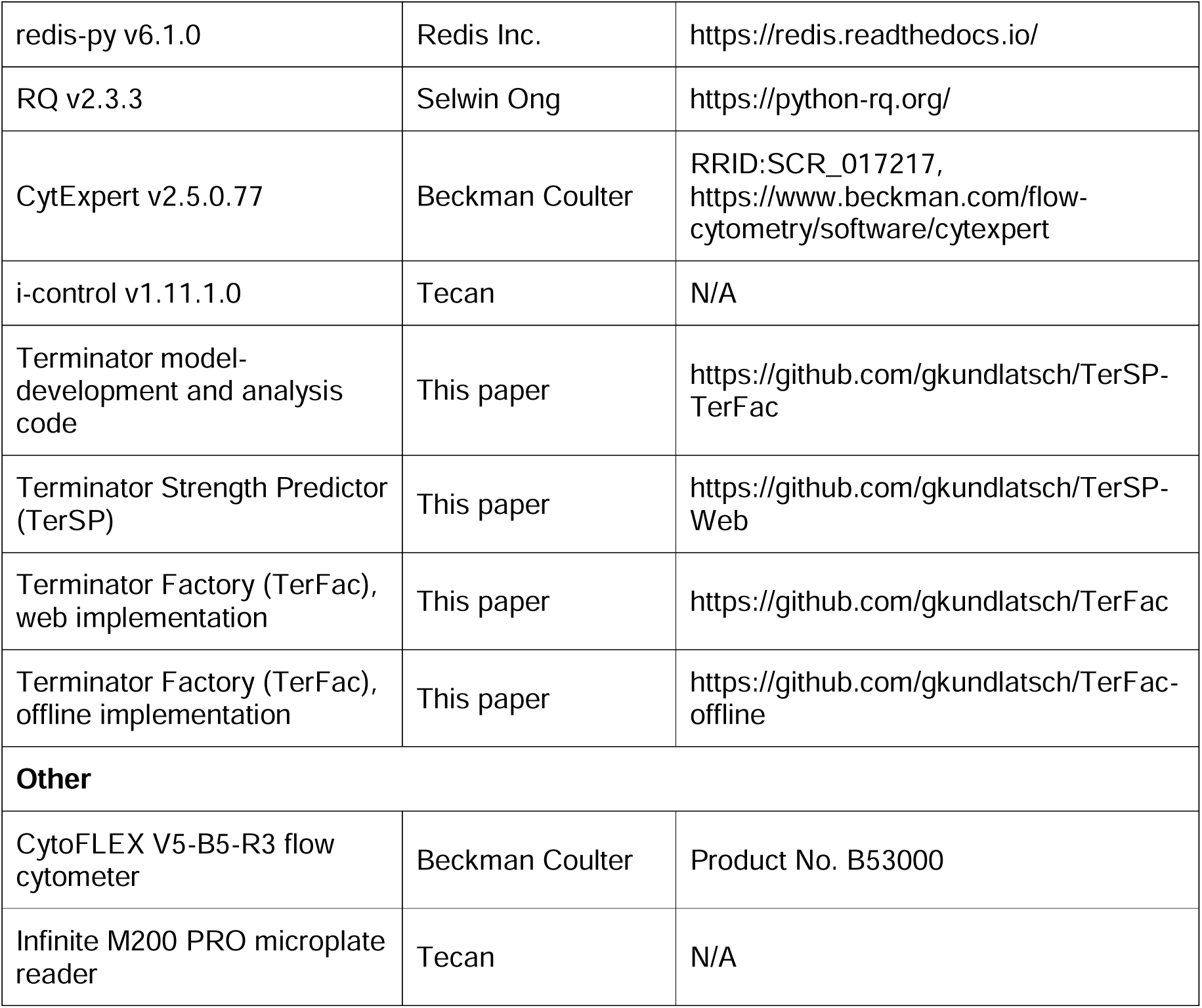
KEY RESOURCES TABLE.

## REFERENCES

1. Cui, W. et al. (2021) Data-driven and in silico-assisted design of broad host-range minimal intrinsic terminators adapted for bacteria. ACS Synth. Biol. 10, 1438–1450

2. Ray-Soni, A. et al. (2016) Mechanisms of bacterial transcription termination: all good things must end. Annu. Rev. Biochem. 85, 319–347

3. Roberts, J. and Park, J.-S. (2004) Mfd, the bacterial transcription repair coupling factor: translocation, repair and termination. Curr. Opin. Microbiol. 7, 120–125

4. Hao, Z., et al. (2021) Rho-dependent transcription termination: a revisionist view. Transcription 12, 171–181

5. You, L. et al. (2023) Structural basis for intrinsic transcription termination. Nature 613, 783– 789

6. Mairhofer, J. et al. (2015) Preventing T7 RNA Polymerase Read-through Transcription – A Synthetic Termination Signal Capable of Improving Bioprocess Stability. ACS Synth. Biol. 4, 265–273

7. Castillo-Hair, S.M. et al. (2019) Optogenetic control of Bacillus subtilis gene expression. Nat. Commun. 10, 3099

8. Chen, Y.-J. et al. (2013) Characterization of 582 natural and synthetic terminators and quantification of their design constraints. Nat. Methods 10, 659–664

9. Brendel, V. and Trifonov, E.N. (1984) A computer algorithm for testing potential prokaryotic terminators. Nucleic Acids Res. 12, 4411–4427

10. Brendel, V. et al. (1986) Terminators of transcription with RNA polymerase from Escherichia coli: what they look like and how to find them. J. Biomol. Struct. Dyn. 3, 705–723

11. Carafa, Y., d’Aubenton et al. (1990) Prediction of rho-independent Escherichia coli transcription terminators: a statistical analysis of their RNA stem-loop structures. J. Mol. Biol. 216, 835–858

12. Ermolaeva, M.D. et al. (2000) Prediction of transcription terminators in bacterial genomes. J. Mol. Biol. 301, 27–33

13. Lesnik, E.A. et al. (2001) Prediction of rho-independent transcriptional terminators in Escherichia coli. Nucleic Acids Res. 29, 3583–3594

14. de Hoon, M.J.L., et al. (2005) Prediction of transcriptional terminators in Bacillus subtilis and related species. PLoS Comput. Biol. 1, e25

15. Kingsford, C.L. et al. (2007) Rapid, accurate, computational discovery of Rho-independent transcription terminators illuminates their relationship to DNA uptake. Genome Biol. 8, R22

16. Cambray, G., et al. (2013) Measurement and modeling of intrinsic transcription terminators. Nucleic Acids Res. 41, 5139–5148

17. Zhai, W. et al. (2022) Sequence and thermodynamic characteristics of terminators revealed by FlowSeq and the discrimination of terminators strength. Synth. Syst. Biotechnol. 7, 1046– 1055

18. He, Z. et al. (2020) Evaluating terminator strength based on differentiating effects on transcription and translation. ChemBioChem 21, 2067–2072

19. Angenent-Mari, N.M. et al. (2020) A deep learning approach to programmable RNA switches. Nat. Commun. 11, 5057

20. Zhou, H. and Zhang, S. (2022) Recent development of fluorescent light-up RNA aptamers. Crit. Rev. Anal. Chem. 52, 1644–1661

21. McDowell, J.C. et al. (1994) Determination of intrinsic transcription termination efficiency by RNA polymerase elongation rate. Science. 266, 822–825

22. Bains, J.K. et al. (2023) Cell-free transcription-translation system: a dual read-out assay to characterize riboswitch function. Nucleic Acids Res. 51, e82

23. Zadeh, J.N. et al. (2011) NUPACK: Analysis and design of nucleic acid systems. J. Comput. Chem. 32, 170–173

24. Dunn, J.J. et al. (1983) Complete nucleotide sequence of bacteriophage T7 DNA and the locations of T7 genetic elements. J. Mol. Biol. 166, 477–535

25. Calvopina-Chavez, D.G. et al. (2022) Engineering efficient termination of bacteriophage T7 RNA polymerase transcription. G3 12, jkac070

26. Kang, D.H. et al. (2022) RoboMoClo: a robotics-assisted modular cloning framework for multiple gene assembly in biofoundry. ACS Synth. Biol. 11, 1336–1348

27. Stukenberg, D. et al. (2021) The Marburg Collection: A Golden Gate DNA assembly framework for synthetic biology applications in Vibrio natriegens. ACS Synth. Biol. 10, 1904– 1919

28. Xu, M. et al. (2022) Development and application of transcription terminators for polyhydroxylkanoates production in halophilic Halomonas bluephagenesis TD01. Front. Microbiol. 13, 941306

29. Li, J. et al. (2025) Intelligent Design of Escherichia coli Terminators by Coupling Prediction and Generation Models. ACS Synth. Biol. 14, 3744–3752

30. Chen, Z. et al. (2020) iLearn: an integrated platform and meta-learner for feature engineering, machine-learning analysis and modeling of DNA, RNA and protein sequence data. Brief. Bioinform. 21, 1047–1057

31. Chen, W. et al. (2014) PseKNC: a flexible web server for generating pseudo K-tuple nucleotide composition. Anal. Biochem. 456, 53–60

32. Gardner, P.P. et al. (2011) RNIE: genome-wide prediction of bacterial intrinsic terminators. Nucleic Acids Res. 39, 5845–5852

33. Taheri Ghahfarokhi, S.M.A. and Peña-Castillo, L. (2025) BacTermFinder: a comprehensive and general bacterial terminator finder using a CNN ensemble. NAR Genomics Bioinforma. 7, lqaf016

34. Tjaden, B. (2026) TerminatorNet: comprehensive identification of intrinsic transcription terminators in bacteria. Bioinformatics 42, btag116

35. Lin, M.-T. et al. (2016) Novel utilization of terminators in the design of biologically adjustable synthetic filters. ACS Synth. Biol. 5, 365–374

36. Sobczak, K. et al. (2003) RNA structure of trinucleotide repeats associated with human neurological diseases. Nucleic Acids Res. 31, 5469–5482

37. Kiliszek, A. et al. (2012) Crystallographic characterization of CCG repeats. Nucleic Acids Res. 40, 8155–8162

38. Zadeh, J.N. et al. (2011) Nucleic acid sequence design via efficient ensemble defect optimization. J. Comput. Chem. 32, 439–452

39. Zuber, J. et al. (2022) Nearest neighbor rules for RNA helix folding thermodynamics: improved end effects. Nucleic Acids Res. 50, 5251–5262

40. Popenda, M. et al. (1997) Solution structure of RNA duplexes containing alternating CG base pairs: NMR study of r (CGCGCG) 2 and 2′-O-Me (CGCGCG) 2 under low salt conditions. Nucleic Acids Res. 25, 4589–4598

41. Kailasam, S., et al. (2014) Sequence dependent variations in RNA duplex are related to non-canonical hydrogen bond interactions in dinucleotide steps. BMC Res. Notes 7, 83

42. Lubkowska, L. et al. (2011) RNA folding in transcription elongation complex: implication for transcription termination. J. Biol. Chem. 286, 31576–31585

43. Chen, Y. et al. (2026) Pipeline and insulated dual-reporter system to predict and quantify strong terminators from RNA-seq datasets for metabolic engineering. Trends Biotechnol. 44, 1452–1477

44. Shannon, C.E. (1948) A mathematical theory of communication. Bell Syst. Tech. J. 27, 379– 423

45. Tian, Y. and Zhang, Y. (2022) A comprehensive survey on regularization strategies in machine learning. Inf. Fusion 80, 146–166

46. Lundberg, S.M. and Lee, S.-I. (2017) A unified approach to interpreting model predictions. Adv. Neural Inf. Process. Syst. 30

47. Kendall, M.G. (1938) A new measure of rank correlation. Biometrika 30, 81–93

48. Kendall, M.G. (1945) The treatment of ties in ranking problems. Biometrika 33, 239–251

49. Kartje, Z.J. et al. (2021) Revisiting T7 RNA polymerase transcription in vitro with the Broccoli RNA aptamer as a simplified real-time fluorescent reporter. J. Biol. Chem. 296, 100175

50. Chen, T. and Guestrin, C. (2016) XGBoost: A scalable tree boosting system. in Proceedings of the 22nd ACM SIGKDD international conference on knowledge discovery and data mining, pp. 785–794

51. Pedregosa, F. et al. (2011) Scikit-learn: Machine learning in Python. J. Mach. Learn. Res. 12, 2825–2830

52. McKinney, W. (2010) Data structures for statistical computing in Python. in Proceedings of the 9th Python in Science Conference, 445, 51–56

53. Harris, C.R. et al. (2020) Array programming with NumPy. Nature 585, 357–362

54. Virtanen, P. et al. (2020) SciPy 1.0: fundamental algorithms for scientific computing in Python. Nat. Methods 17, 261–272

55. Hunter, J.D. (2007) Matplotlib: A 2D graphics environment. Comput. Sci. Eng. 9, 90–95

56. Varoquaux, G. et al. (2024) joblib/joblib: 1.4.2. Zenodo.

