## Supplementary material for "Rational design of intrinsic transcription terminators with programmable strength"

**Supplementary Table 1. Comparison of computational approaches for intrinsic terminator identification, strength prediction, and design.**

Computational approaches to intrinsic terminator prediction began to emerge in the 1980s, with Brendel and colleagues developing some of the earliest rule-based methods [1,2]. Building on this line of work, d'Aubenton Carafa *et al.* [3] combined hairpin stability with the number and position of thymine residues in the downstream T-rich tract (U-tract in RNA), providing an early quantitative scoring scheme for terminator detection and an early attempt to estimate *in vitro* termination. Moving to genome-scale prediction, Ermolaeva *et al.* [4] developed TransTerm, which combined a hairpin energy score and U-tail score with the genomic location of candidate terminators relative to annotated genes. By comparing terminator-like structures in intergenic and intragenic regions, the method assigned each candidate an empirical confidence estimate and enabled genome-scale prediction across 12 bacteria. Lesnik *et al.* [5] extended genome-scale detection using RNAMotif, combining explicit sequence and structural constraints with a thermodynamic score incorporating hairpin stability and RNA–DNA hybrid energetics across distinct portions of the U-tract. Their framework also included an upstream A-region to aid identification of bidirectional terminators.

Adapting this scoring framework to *B. subtilis*, de Hoon et al. [6] derived a decision rule from 463 experimentally supported terminating regions and 567 non-terminating regions, combining normalized stem-loop free energy with a weighted U-stretch score. The same rule achieved high sensitivity across 57 Firmicutes, supporting the conclusion that key terminator properties are broadly conserved. Improving the speed of genome-scale identification, Kingsford *et al.* [7] developed a fast method to identify low-energy hairpins followed by a U-tract and assign a score reflecting how unlikely each candidate was to have arisen by chance. The algorithm was substantially faster than previous methods, scanning a 4-megabase genome in under 50 s. Those approaches, however, were developed for terminator detection rather than quantitative prediction of termination efficiency.

Later approaches moved from terminator detection toward quantitative sequence-to-function modeling. Cambray *et al.* [8] developed a multiple linear regression model to predict termination efficiency from terminator sequence features, while using cotranscriptional folding simulations to identify context-dependent structures that could confound these relationships. Their best-performing four-feature model incorporated normalized hairpin stability, the identity of the closing stack at the stem base, U-tract quality, and downstream DNA composition. However, this model was trained on only 31 of the 54 terminators used for modeling, after excluding low-efficiency, low-folding-frequency, and extended terminators, indicating that its strongest predictive performance depended on restricting the terminators considered. Expanding substantially, Chen *et al.* [9] characterized 582 natural and synthetic *E. coli* terminators and identified sequence features associated with terminator strength, incorporating these into a biophysical model based on the hybrid-shearing mechanism of termination, with hairpin nucleation followed by U-tract ratcheting. The model explained only a fraction of the observed variation in terminator strength. Importantly, systematic exchange of apparently favorable sequence features among strong terminator scaffolds revealed pronounced context dependence: features that increased strength in one scaffold could reduce it in another. Thus, identifying individual features associated with strength did not provide a general rule for combining them to construct stronger terminators.

Subsequent studies began to address terminator construction more directly. Cui *et al.* [10] developed a forward-design framework for broad-host-range minimal intrinsic terminators, deriving sequence-activity rules from hairpin thermodynamics and U-tract length to construct *de novo* minimal terminators. Termination efficiency was subsequently tuned over a broad range by systematically varying U-tract length within selected designed scaffolds, enabling rule-based strength modulation without a quantitative sequence-to-strength predictor. Xu *et al.* [11] developed a multiple linear regression model combining sequence and thermodynamic features, trained on the 582-terminator Chen dataset, and combined this modeling with empirically identified structural main contributors to guide rational optimization and *de* *novo* terminator construction in *Halomonas bluephagenesis*. The three *de novo* designs achieved termination efficiencies of 88.1-94.7%, exceeding the T7 control, although none reached the nominal >97.5% TE criterion corresponding to the Chen *et al*. strong-terminator definition. Extending these data-driven approaches, Zhai *et al.* [12] combined sequence and thermodynamic features to train an XGBoost classifier that distinguished strong from weak terminators. Finally, during the preparation of this manuscript, Li *et al.* [13] published a model trained on the same dataset using a substantially different and much larger feature space based on k-mer, CKSNAP, PseKNC, thermodynamic, and secondary-structure descriptors; although they reported a higher test $R^2$ than ours, the experimental validation did not establish that any of the 18 generated terminators outperformed B0010, which itself was not the strongest terminator in the Chen dataset. The differences between the two approaches are discussed in greater detail in the Discussion. In parallel, computational methods for intrinsic terminator identification have continued to advance through probabilistic and machine-learning approaches, including RNIE [14], iTerm-PseKNC [15], termNN [16], BacTermFinder [17], and TerminatorNet [18]; however, these methods primarily address terminator detection rather than quantitative strength prediction or design. SVR: Support Vector Regression; WGAN-GP: Wasserstein generative adversarial network with gradient penalty.

| **Study** | **Approach** | **Output** | **Quantitative sequence-to-strength prediction** | **Model-enabled design capability** | **Limitation** |
|---|---|---|---|---|---|
| d'Aubenton Carafa *et al.* [3] | Empirical scoring model based on hairpin stability and downstream U-tract features | Terminator score | Limited | No | Detection-focused heuristic; efficiency was only estimated approximately from the terminator score |
| Ermolaeva *et al.* [4] | Hairpin energy and U-tract scores combined with genomic context | Empirical confidence estimate | No | No | Predicts terminator confidence rather than strength |
| Lesnik *et al.* [5] | RNAMotif sequence and structural constraints with thermodynamic scoring | Candidate terminators and thermodynamic scores | No | No | Mechanistically informed detection score without calibration to measured termination efficiency |
| de Hoon *et al.* [6] | Decision rule combining normalized stem-loop free energy with a weighted U-tract score | Terminator/ non-terminator classification | No | No | Detects terminators rather than predicting quantitative strength |
| Kingsford *et al.* [7] | Fast hairpin/U-tract search with background-based scoring | Chance-based terminator score | No | No | Ranks candidate terminators but does not predict |

| | | | | | |
|---|---|---|---|---|---|
| | | | | | quantitative strength |
| Cambray *et al.* [8] | Multiple linear regression using sequence and biophysical features, informed by cotranscriptional folding simulations | Predicted termination efficiency | Yes | No | Best-performing model was restricted to 31 of 54 terminators after excluding low-efficiency, low-folding-frequency, and extended terminators |
| Chen *et al.* [9] | Biophysical model based on thermodynamic contributions of the U-tract, hairpin loop/stem base, and A-tract | Predicted termination strength | Yes | Feature-guided construction | Favorable sequence features did not combine predictably across scaffolds |
| Cui *et al.* [10] | Data-driven sequence–activity rules combining hairpin thermodynamics and U-tract length with in silico hairpin design | Designed minimal terminators with tunable termination efficiency | No | Rule-based forward design | Strength was tuned primarily by varying U-tract length within designed scaffolds rather than predicted or designed toward a specified strength |
| Xu *et al.* [11] | Multiple linear regression using sequence and thermodynamic features coupled to feature-guided engineering | Predicted termination efficiency and rational/de novo terminator designs | Yes | Feature-guided rational design | *De novo* designs achieved 88.1–94.7% TE, remaining below the strong-terminator threshold defined by Chen *et al.* |
| Zhai *et al.* [12] | XGBoost classifier using sequence and thermodynamic features | Strong / weak classification | No | Variant design for classifier validation | Binary output does not predict quantitative strength, with overlap between predicted classes for moderate-strength terminators |
| Li *et al.* [13] | SVR using 3-mer, thermodynamic, and RNA secondary-structure features, | Predicted termination strength and generated candidate sequences | Yes | Generate-and-screen design | Of 18 generated candidates tested experimentally, five showed termination efficiencies |

| | | | | | |
|---|---|---|---|---|---|
| | coupled to WGAN-GP for sequence generation | | | | comparable to B0010; the study did not demonstrate that any candidate clearly outperformed B0010, which was not the strongest terminator in the Chen source dataset |
| This work | Interpretable sequence-to-function model coupled to inverse design | Continuous predicted strength and candidate sequences for specified strengths | Yes | Target-driven inverse design | Designed terminators validated *in vivo* and *in vitro*, but utility in a downstream application remains to be demonstrated |

**Supplementary Table 2.** Features evaluated in this study. The sixteen features with the best performance, which were subsequently used, are highlighted in green.

| Name | Region | Description |
|---|---|---|
| Entropy_A_tract | A-tract | Shannon entropy |
| A_Tract_state-change | A-tract | Number of nucleotide transitions |
| A%_6_A_tract | A-tract | % of adenines in the last six nucleotides |
| C%_6_A_tract | A-tract | % of cytosines in the last six nucleotides |
| Hairpin Length without Loop | Hairpin | Number of base-paired nucleotides in the hairpin |
| Length Loop | Hairpin | Number of unpaired nucleotides in the hairpin |
| GC%_Loop | Hairpin | % of G or C among unpaired nucleotides in the hairpin |
| Entropy_HP_w/o_Loop | Hairpin | Shannon entropy of paired nucleotides in the hairpin |
| Hairpin_state-change | Hairpin | Number of nucleotide transitions |
| GC_Inicial_Hairpin | Hairpin | Number of G or C starting from the first hairpin nucleotide before encountering an A or U |
| Entropy_U_tract | U-tract | Shannon entropy |
| U_Tract_state-change | U-tract | Number of nucleotide transitions |
| A%_6_U_tract | U-tract | % of adenines in the first six nucleotides |
| C%_U_tract | U-tract | Total % of cytosines |
| U%_6_U_tract | U-tract | % of uracils in the first six nucleotides |
| U%_10_U_tract | U-tract | % of uracils in the first ten nucleotides |
| Ti_Tv_Ratio_A_Tract | A-tract | Ratio of transitions to transversions |
| a-factor | A-tract | A score was given to each adenine in the sequence, with positions closer to the hairpin receiving higher values |
| A%_1_A_tract | A-tract | % of adenines in the last nucleotide |
| A%_2_A_tract | A-tract | % of adenines in the last two nucleotides |
| A%_3_A_tract | A-tract | % of adenines in the last three nucleotides |
| A%_4_A_tract | A-tract | % of adenines in the last four nucleotides |
| A%_5_A_tract | A-tract | % of adenines in the last five nucleotides |
| A%_7_A_tract | A-tract | % of adenines in the last seven nucleotides |

| | | |
|---|---|---|
| A%_A_tract | A-tract | Total % of adenines |
| C%_1_A_tract | A-tract | % of cytosines in the last nucleotide |
| C%_2_A_tract | A-tract | % of cytosines in the last two nucleotides |
| C%_3_A_tract | A-tract | % of cytosines in the last three nucleotides |
| C%_4_A_tract | A-tract | % of cytosines in the last four nucleotides |
| C%_5_A_tract | A-tract | % of cytosines in the last five nucleotides |
| C%_7_A_tract | A-tract | % of cytosines in the last seven nucleotides |
| C%_A_tract | A-tract | Total % of cytosines |
| G%_1_A_tract | A-tract | % of guanines in the last nucleotide |
| G%_2_A_tract | A-tract | % of guanines in the last two nucleotides |
| G%_3_A_tract | A-tract | % of guanines in the last three nucleotides |
| G%_4_A_tract | A-tract | % of guanines in the last four nucleotides |
| G%_5_A_tract | A-tract | % of guanines in the last five nucleotides |
| G%_6_A_tract | A-tract | % of guanines in the last six nucleotides |
| G%_7_A_tract | A-tract | % of guanines in the last seven nucleotides |
| G%_A_tract | A-tract | Total % of guanines |
| U%_1_A_tract | A-tract | % of uracils in the last nucleotide |
| U%_2_A_tract | A-tract | % of uracils in the last two nucleotides |
| U%_3_A_tract | A-tract | % of uracils in the last three nucleotides |
| U%_4_A_tract | A-tract | % of uracils in the last four nucleotides |
| U%_5_A_tract | A-tract | % of uracils in the last five nucleotides |
| U%_6_A_tract | A-tract | % of uracils in the last six nucleotides |
| U%_7_A_tract | A-tract | % of uracils in the last seven nucleotides |
| U%_A_tract | A-tract | Total % of uracils |
| GC%_1_A_tract | A-tract | % of G or C in the last nucleotide |
| GC%_2_A_tract | A-tract | % of G or C in the last two nucleotides |
| GC%_3_A_tract | A-tract | % of G or C in the last three nucleotides |
| GC%_4_A_tract | A-tract | % of G or C in the last four nucleotides |

| | | |
|---|---|---|
| GC%_5_A_tract | A-tract | % of G or C in the last five nucleotides |
| GC%_6_A_tract | A-tract | % of G or C in the last six nucleotides |
| GC%_7_A_tract | A-tract | % of G or C in the last seven nucleotides |
| GC%_A_tract | A-tract | Total % of G or C |
| GC%_Base_1 | Hairpin | % of G or C in the first nucleotide of the hairpin |
| 'GC%_Base_2 | Hairpin | % of G or C in the first two nucleotides of the hairpin |
| GC%_Base_3 | Hairpin | % of G or C in the first three nucleotides of the hairpin |
| GC%_HP | Hairpin | % of G or C among paired nucleotides in the hairpin |
| %A_HP | Hairpin | % of adenines among paired nucleotides in the hairpin |
| %C_HP | Hairpin | % of cytosines among paired nucleotides in the hairpin |
| %G_HP | Hairpin | % of guanines among paired nucleotides in the hairpin |
| %U_HP | Hairpin | % of uracils among paired nucleotides in the hairpin |
| %A_Loop | Hairpin | % of adenines among unpaired nucleotides in the hairpin |
| %C_Loop | Hairpin | % of cytosines among unpaired nucleotides in the hairpin |
| %G_Loop | Hairpin | % of guanines among unpaired nucleotides in the hairpin |
| %U_Loop | Hairpin | % of uracils among unpaired nucleotides in the hairpin |
| GC%_Loop | Hairpin | % of G or C among unpaired nucleotides in the hairpin |
| Entropy_Loop | Hairpin | Shannon entropy of unpaired nucleotides in the hairpin |
| coef_est | Hairpin | Product of attributes 'Hairpin Length without Loop' and '%GC_HP' |
| Ti_Tv_Ratio_Hairpin | Hairpin | Ratio of transitions to transversions |
| Ti_Tv_Ratio_U_Tract | U-tract | Ratio of transitions to transversions |
| u-factor | U-tract | A score was given to each uracil in the sequence, with positions closer to the hairpin receiving higher values |
| A%_1_U_tract | U-tract | % of adenines in the first nucleotide |
| A%_2_U_tract | U-tract | % of adenines in the first two nucleotides |
| A%_3_U_tract | U-tract | % of adenines in the first three nucleotides |
| A%_4_U_tract | U-tract | % of adenines in the first four nucleotides |
| A%_5_U_tract | U-tract | % of adenines in the first five nucleotides |

| | | |
|---|---|---|
| A%_7_U_tract | U-tract | % of adenines in the first seven nucleotides |
| A%_8_U_tract | U-tract | % of adenines in the first eight nucleotides |
| A%_9_U_tract | U-tract | % of adenines in the first nine nucleotides |
| A%_10_U_tract | U-tract | % of adenines in the first ten nucleotides |
| A%_11_U_tract | U-tract | % of adenines in the first eleven nucleotides |
| A%_U_tract | U-tract | Total % of adenines |
| C%_1_U_tract | U-tract | % of cytosines in the first nucleotide |
| C%_2_U_tract | U-tract | % of cytosines in the first two nucleotides |
| C%_3_U_tract | U-tract | % of cytosines in the first three nucleotides |
| C%_4_U_tract | U-tract | % of cytosines in the first four nucleotides |
| C%_5_U_tract | U-tract | % of cytosines in the first five nucleotides |
| C%_6_U_tract | U-tract | % of cytosines in the first six nucleotides |
| C%_7_U_tract | U-tract | % of cytosines in the first seven nucleotides |
| C%_8_U_tract | U-tract | % of cytosines in the first eight nucleotides |
| C%_9_U_tract | U-tract | % of cytosines in the first nine nucleotides |
| C%_10_U_tract | U-tract | % of cytosines in the first ten nucleotides |
| C%_11_U_tract | U-tract | % of cytosines in the first eleven nucleotides |
| G%_1_U_tract | U-tract | % of guanines in the first nucleotide |
| G%_2_U_tract | U-tract | % of guanines in the first two nucleotides |
| G%_3_U_tract | U-tract | % of guanines in the first three nucleotides |
| G%_4_U_tract | U-tract | % of guanines in the first four nucleotides |
| G%_5_U_tract | U-tract | % of guanines in the first five nucleotides |
| G%_6_U_tract | U-tract | % of guanines in the first six nucleotides |
| G%_7_U_tract | U-tract | % of guanines in the first seven nucleotides |
| G%_8_U_tract | U-tract | % of guanines in the first eight nucleotides |
| G%_9_U_tract | U-tract | % of guanines in the first nine nucleotides |
| G%_10_U_tract | U-tract | % of guanines in the first ten nucleotides |
| G%_11_U_tract | U-tract | % of guanines in the first eleven nucleotides |

| | | |
|---|---|---|
| G%_U_tract | U-tract | Total % of guanines |
| U%_1_U_tract | U-tract | % of uracils in the first nucleotide |
| U%_2_U_tract | U-tract | % of uracils in the first two nucleotides |
| U%_3_U_tract | U-tract | % of uracils in the first three nucleotides |
| U%_4_U_tract | U-tract | % of uracils in the first four nucleotides |
| U%_5_U_tract | U-tract | % of uracils in the first five nucleotides |
| U%_7_U_tract | U-tract | % of uracils in the first seven nucleotides |
| U%_8_U_tract | U-tract | % of uracils in the first eight nucleotides |
| U%_9_U_tract | U-tract | % of uracils in the first nine nucleotides |
| U%_11_U_tract | U-tract | % of uracils in the first eleven nucleotides |
| U%_U_tract | U-tract | Total % of uracils |
| GC%_1_U_tract | U-tract | % of G or C in the first nucleotide |
| GC%_2_U_tract | U-tract | % of G or C in the first two nucleotides |
| GC%_3_U_tract | U-tract | % of G or C in the last three nucleotides |
| GC%_4_U_tract | U-tract | % of G or C in the first four nucleotides |
| GC%_5_U_tract | U-tract | % of G or C in the first five nucleotides |
| GC%_6_U_tract | U-tract | % of G or C in the first six nucleotides |
| GC%_7_U_tract | U-tract | % of G or C in the first seven nucleotides |
| GC%_8_U_tract | U-tract | % of G or C in the first eight nucleotides |
| GC%_9_U_tract | U-tract | % of G or C in the first nine nucleotides |
| GC%_10_U_tract | U-tract | % of G or C in the first ten nucleotides |
| GC%_11_U_tract | U-tract | % of G or C in the first eleven nucleotides |
| GC%_U_tract | U-tract | Total % of G or C |

**Supplementary Table 3. Dataset imbalances and their implications for model interpretation.** Here, we summarize characteristics of the dataset that may have influenced model development and interpretation.

During feature selection, among U-tract composition metrics, C content was more influential than either G or GC content. However, because C- and G-containing U-tracts were unevenly represented in the relatively limited training dataset, this apparent preference may reflect dataset composition rather than a specific functional role for cytosine over guanine. A larger dataset with more balanced representation of C- and G-containing U-tracts would be required to distinguish between these possibilities. Interestingly, cytosine content in the final six nucleotides of the A-tract was also selected as an informative feature. In this region, the imbalance between C and G representation was less pronounced but still present, and the apparent preference for cytosine may therefore also partly reflect dataset composition.

Terminator strengths were also strongly imbalanced, with less than one fifth of the terminators characterized as strong. This limits the information available for learning quantitative differences among strong terminators, especially given the small size of the dataset. Another important consideration when evaluating model performance is that the strength scale adopted by Chen *et al.* magnifies small differences among highly efficient terminators: strengths of 100, 200, and 300 correspond to termination efficiencies of approximately 99.0%, 99.5%, and 99.67%, respectively. Consequently, comparatively large prediction errors on the strength scale may correspond to much smaller differences in termination efficiency and can disproportionately penalize model performance as measured by $R^2$, even when the terminator is assigned to the correct functional class.

To further evaluate model reliability, we first assessed weak/strong classification across all held-out sequences. The model achieved 90.21% accuracy, indicating strong classification performance. However, because 83.16% of the dataset consists of weak terminators, a trivial classifier assigning every sequence to the weak class would already achieve 83.16% accuracy. We therefore performed a second analysis restricted to experimentally strong terminators, asking how often these sequences were also predicted as strong. In this more stringent test, 83.67% of strong terminators were correctly classified, demonstrating that the model's performance was not simply driven by the dominant weak class.

Finally, an additional limitation of the dataset became apparent during *in vivo* validation. B0010 showed a strength comparable to Tmax in our assay, whereas in the Chen dataset its reported strength was approximately 4.6-fold lower. This result is consistent with the widespread use of B0010 as an efficient terminator in synthetic-biology applications and raises the possibility that its strength was underestimated during construction of the original dataset. Such discrepancies are particularly relevant given the small number of very strong terminators available for model training. Nevertheless, the model-designed TK outperformed both B0010 and Tmax when evaluated under the same assay framework used by Chen *et al.*, suggesting that the model captured broader sequence determinants of efficient termination despite the potentially atypical training value reported to B0010.

| **C/G composition of the U-tract** | **Number of terminators** |
|---|---|
| Cytosine and guanine | 385 (66.15%) |
| Only cytosine | 141 (24.23%) |
| Only guanine | 47 (8.08%) |
| Neither | 9 (1.55%) |
| **C/G composition of the final six nucleotides of the A-tract** | **Number of terminators** |

| | |
|---|---|
| Cytosine and guanine | 256 (43.99%) |
| Only cytosine | 135 (23.20%) |
| Only guanine | 99 (17.01%) |
| Neither | 92 (15.81%) |
| **Strength distribution** | **Number of terminators** |
| <40 | 484 (83.16%) |
| 40 - <100 | 46 (7.90%) |
| 100 - <200 | 27 (4.64%) |
| 200 - 300 | 19 (3.26%) |
| >300 | 6 (1.03%) |

**Supplementary Table 4. Comparison of model-selected features with previous descriptions of intrinsic terminator sequence determinants.** Here, we compare the descriptors identified in this work both with previous observations in the literature and with those reported by Chen *et al.*, whose terminator library was used to train our model.

The main differences emerged within the hairpin. First, Shannon entropy and state-change are, to our knowledge, previously undescribed descriptors of terminator strength and, therefore, have not been previously discussed in the literature. Stronger predictions were associated with greater nucleotide diversity, captured by entropy, and more frequent changes in nucleotide identity along the hairpin sequence, captured by state-change. These associations describe sequence composition and organization, their possible structural implications are considered as hypotheses in the Discussion.

A second distinction concerned initial GC count. Previous studies emphasized the contribution of basal stem pairing, and Chen *et al.* quantified the energetic contribution of the first three base pairs. We propose, however, that the relevance of GC content expands beyond this traditionally analyzed window. In our model, the length of the contiguous basal GC region was more informative than GC content in the first 3, 2 or 1 positions. This distinction became particularly apparent during terminator optimization: as the permitted hairpin length increased, maximum-strength designs also acquired progressively longer basal GC regions, reaching nine initial GC pairs in the longest optimized sequence and eight in TK. These observations suggest that hairpin length and basal GC pairing should be considered jointly, with their possible interaction remaining to be tested experimentally.

Hairpin length also differed remarkably from previous descriptions. Chen *et al.* found no correlation between stem length and terminator strength across their analyzed library. This is consistent with previous literature. For example, Wilson and von Hippel [19] proposed an optimal stem of approximately 8-9 bp for λtR2 and observed that further stem extension reduced termination despite increasing predicted hairpin stability. These findings illustrate that the effect of stem length cannot be inferred from equilibrium stability alone. Mechanistic work has emphasized the need to coordinate hairpin formation with the limited opportunity for termination during transcription. Lubkowska *et al.* [20] showed that sequestration of the downstream hairpin arm by RNA polymerase makes productive folding sensitive to competing interactions involving upstream RNA. They proposed that longer stems could increase susceptibility to such interactions, but could also favor earlier hairpin nucleation and thereby facilitate timely completion of the stem. Thus, previous mechanistic discussions accommodate both beneficial and detrimental effects of stem length, depending on sequence context and folding dynamics. In our model, longer hairpins were associated with stronger predictions. TK encodes the longest hairpin permitted during optimization, a predicted 24-bp stem, substantially exceeding the optimum proposed for λtR2. Its experimentally validated performance, in opposition to the previous literature, shows that an extended stem can support efficient termination, although the contributions of its associated sequence features remain unresolved. Targeted variation of the basal GC region, entropy, and state-change would help distinguish their individual and combined contributions.

The importance of considering these descriptors jointly is further illustrated by miniTK. Restricting hairpin length generated a shorter design that retained a high initial GC count and high nucleotide diversity, with state-change remaining maximal for its available length. Nevertheless, miniTK showed lower termination efficiency than TK. This comparison supports the functional relevance of the combination of features favored by the model.

For the loop, the model identified length and GC content as informative descriptors, associating shorter sequences and lower GC content with stronger predictions. Although Chen *et al.* found no direct correlation between loop length and terminator strength, they associated stronger termination with favorable loop-closure energy and reported that GAAA performed best on average among the loop substitutions tested. These findings are consistent with the longstanding observation that tetraloops are prevalent among intrinsic terminators [3]. The model therefore recapitulates characteristics of established loop motifs, including the compact size and low GC content of GAAA.

For the A-tract, the model associated stronger predictions with lower entropy and state-change, together with higher A content and lower C content in the six nucleotides immediately upstream of the hairpin. The higher A content is consistent with the previously recognized A-rich sequence bias immediately upstream of intrinsic terminator hairpins[3], and the most informative compositional window matches the region in which Chen *et al.* reported A enrichment among strong terminators. Here, Shannon entropy and state-change serve as new descriptors of a previously recognized sequence pattern.

For the U-tract, lower entropy and state-change were associated with stronger predictions, consistent with a more uniform, U-rich sequence and longstanding experimental evidence for the importance of U-tract integrity and length [21,22]. Higher U content and lower A content in the six nucleotides immediately downstream of the hairpin agree with the proximal U enrichment reported by Chen *et al.*, whose thermodynamic analysis considered an eight-nucleotide window. The selection of U content across ten nucleotides and C content across the full tract suggests that broader downstream composition also provides predictive information. This is consistent with earlier evidence that sequences beyond the termination site can influence efficiency in a context-dependent manner [23,24]. Together, these descriptors capture established U-tract sequence preferences while suggesting that informative variation extends beyond the proximal region.

| Feature | Previous literature | Chen *et al.* [9] | This work |
|---|---|---|---|
| Hairpin entropy | No direct precedent was identified as a hairpin descriptor. | Not evaluated. | Higher entropy was associated with stronger termination, introducing nucleotide diversity as a new hairpin descriptor. |
| Hairpin state-change | No direct precedent was identified as a hairpin descriptor. | Not evaluated. | Higher state-change was associated with stronger termination, identifying nucleotide alternation as a new hairpin descriptor. |
| Hairpin length | Comparative analyses identified relatively short stems as characteristic of intrinsic terminators[1,3], while direct manipulation of the tR2 terminator suggested an optimum of approximately 8-9 base pairs [19]. | No correlation between stem length and terminator strength was observed, and length was not included in the biophysical model. | Longer hairpins were associated with stronger predictions, and very long stems remained compatible with high termination strength. |
| Initial GC count (hairpin) | Strong GC pairing at the hairpin base, particularly within the first 3 bp, has been associated with efficient termination[3,8]. | GC enrichment was concentrated at the stem base, and the model considered the free energy of the first three basal base pairs. | The length of the contiguous basal GC-rich region was more informative than GC content within a fixed three-position window, particularly for longer hairpins. |
| Loop length | Short loops, particularly stable tetraloops, have been described as characteristic of intrinsic terminators [1,3]. | No direct correlation with strength was observed, although tetraloops were the most frequent loop size. | Shorter loops were favored, consistent with the prevalence of compact terminator loops. |

| | | | |
|---|---|---|---|
| GC% Loop | Specific stable loop motifs such as GAAA and UUCG have been associated with intrinsic terminators[3,19]; GC content itself has not been established as a determinant. | Loop sequence and closure energy were associated with strength; GAAA was the best loop tested on average. GC% was not evaluated independently. | Lower loop GC content favored stronger predictions, indirectly recapitulating the preference for the GAAA tetraloop described previously. |
| A-tract entropy | A-rich sequences immediately upstream of the hairpin have long been associated with intrinsic terminators [3,25,26]. | An enrichment near the hairpin was associated with stronger termination. | Low entropy can represent A-enrichment. New descriptor capturing established biology. |
| A-Tract state-change | A-rich sequences immediately upstream of the hairpin have long been associated with intrinsic terminators [3,25,26]. | An enrichment near the hairpin was associated with stronger termination. | Low state-change can represent A-enrichment. New descriptor capturing established biology. |
| A% 6 A-tract | A-rich sequences immediately upstream of the hairpin have long been associated with intrinsic terminators [3,25,26]. | An enrichment near the hairpin was associated with stronger termination. | Higher A content is consistent with established biology. |
| C% 6 A-tract | A-rich sequences immediately upstream of the hairpin have long been associated with intrinsic terminators [3,25,26]. | An enrichment near the hairpin was associated with stronger termination. | Lower C content is consistent with established biology. |
| Entropy U-tract | U-rich sequences immediately downstream of the hairpin have long been associated with intrinsic terminators[1,21,22]. | High U content was strongly associated with terminator strength. | Low entropy can represent U-enrichment. New descriptor capturing established biology. |
| U-tract state-change | U-rich sequences immediately downstream of the hairpin have long been associated with intrinsic terminators [1,21,22]. | High U content was strongly associated with terminator strength. | Low state-change can represent U-enrichment. New descriptor capturing established biology. |
| A% 6 U-tract | U-rich sequences immediately downstream of the hairpin have long been associated with intrinsic terminators [1,21,22]. | High U content was strongly associated with terminator strength. | Lower A content is consistent with established biology. |
| U% 6 U-tract | U-rich sequences immediately downstream of the hairpin have long been associated with intrinsic terminators [1,21,22]. | High U content was strongly associated with terminator strength. | Higher U content is consistent with established biology. |
| U% 10 U-tract | Sequence effects extending beyond the canonical U-tract have been reported, with distal positions influencing termination efficiency in a | U enrichment declined progressively along the tract; the quantitative model considered the first 8 nt. | U content across 10 nt was more informative than the proximal 6 nt descriptor, indicating that distal positions remain informative |

| | | | |
|---|---|---|---|
| | context-dependent manner [23,24]. | | |
| C% U-tract | Sequence effects extending beyond the canonical 7-9 nt U-tract have been reported, with distal positions influencing termination efficiency in a context-dependent manner [23,24]. | U enrichment declined progressively along the tract; the quantitative model considered the first 8 nt. | Lower C content across the extended U-tract favored stronger predictions, indicating that distal positions remain informative |

**Supplementary Table 5.** Glossary of terminators evaluated in this work.

| Terminator | Description | Sequence |
|---|---|---|
| T1 | ECK120010810, from the *malZ* transcription unit, was renamed T1 here for simplicity. In the original Chen et al. dataset, this terminator had a measured strength (Ts) of 0.99 ± 0.08, indicating near-equal fluorescence output from the upstream and downstream protein reporters. Thus, it was selected as a low-strength reference terminator expected to permit extensive readthrough. Chen et al.'s model predicted a strength of 3.32 for this sequence. ECK120011168 had a similar measured strength of 0.99 but was not selected because its reported standard deviation was higher (0.22). | CCACTGCTCTGACCACAAGTAATTGTTCAGATTGATAAAACA |
| T41 | The *IlvBN* terminator, associated with the *ilvBN* and *ilvGEDA* transcription units, was renamed T41 here for simplicity. In the original Chen et al. dataset, this terminator had a measured strength of 40.39 ± 12.19, making it the weakest terminator above the threshold of 40 used by Chen et al. to distinguish strong from weak terminators. We therefore adopted this same threshold in the present work, with terminators above this value considered strong. The model reported by Chen et al. predicted a strength of 93.66 for this sequence. | GTAAAAAAGGCAGCCATCTGGCTGCCTTAGTCTCCCCA |
| Tmax | L3S2P21 was the strongest synthetic terminator reported by Chen et al., with a measured strength of 382.13 ± 18.42. It was renamed Tmax here for simplicity. Because it had the highest experimentally quantified strength in the Chen et al. library, Tmax was used as a reference for evaluating the generative capabilities of our system. Chen et al.'s model predicted a strength of 122.71 for Tmax. | AGAAAAGAGGCCTCCCGAAAGGGGGGCCTTTTTTCGTTTT |
| B0010 | One of the most widely used terminators for recombinant expression in bacteria, B0010 has been described as an efficient terminator for this purpose since the 1980s[27]. Originally named T1 from the rrnB operon, we use here the B0010 nomenclature standardized by the iGEM Foundation Registry of Standard Biological Parts to avoid confusion between the | AAATAAAACGAAAGGCTCAGTCGAAAGACTGGGCCTTTCGTTTTATCTGTTG |

| | | |
|---|---|---|
| | name T1 and a terminator strength of 1. In the iGEM Registry alone, B0010 is catalogued as being used in 12,828 parts, illustrating its broad adoption as a genetic part. In the original Chen et al. dataset, B0010 is described as a strong terminator, with a measured strength of 83.57 ± 11.54. However, in our in vivo assay we quantified a higher termination efficiency, more consistent with our expectations given its frequent use. The model reported by Chen et al. predicted a strength of 22.15 for this sequence. | |
| TK | First synthetic terminator developed in this work. | CGAAAAAACCGCGCCGAGGCCGGCGCCGATATACAAATATCGGCGCCGGCCTCGGCGCGGTTTTTTTTCATT |
| Mini TK | A shortened version of TK designed by imposing a maximum hairpin-length constraint during the generation process. | CGAAAAAACGCGCGCGATACAAATCGCGCGCGTTTTTTTTCATT |
| T7wt | Wild-type T7 terminator | AACCCCTTGGGGCCTCTAAACGGGTCTTGAGGGGTTTTTTTT |
| T7opt | A-tract and U-tract from TK combined to the wild-type T7 terminator hairpin. | GTAAAAAACCCCTTGGGGCCTCTAAACGGGTCTTGAGGGGTTTTTTTTCATT |

**Supplementary Table 6. Terminator strength values and corresponding calculated termination efficiencies.** Terminator strength (TS) values were calculated as described by Chen et al. for the in vivo reporter assay:

$$T_s = \left(\frac{\langle GFP \rangle_{Term}}{\langle RFP \rangle_{Term}}\right)\left(\frac{\langle GFP \rangle_0}{\langle RFP \rangle_0}\right)^{-1}$$

The calculation was adjusted for the in vitro assay, using equation:

$$T_s = \left(\frac{\langle mango \rangle_{Term}}{\langle broccoli \rangle_{Term}}\right)\left(\frac{\langle mango \rangle_0}{\langle broccoli \rangle_0}\right)^{-1}$$

The Terminator strength (Ts) relates to the Termination efficiency (TE) through the equation:

$$TE = \left[1 - \left(\frac{1}{T_s}\right)\right] 100$$

TS values were converted to TE to provide a complementary representation of terminator activity and facilitate comparison with previous studies.

| Terminator | $T_S$ | TE |
|---|---|---|
| ***In vivo*** | | |
| TK | 475.1 ± 6.8 | 99.79 ± 0.02 |
| B0010 | 357.4 ± 17.3 | 99.72 ± 0.01 |
| Tmax | 342.2 ± 11.2 | 99.71 ± 0.01 |
| miniTK | 201.2 ± 7.2 | 99.50 ± 0.00 |
| ***In vitro*** | | |
| TK | 2.72 ± 0.24 | 63.09 ± 3.38 |
| miniTK | 2.12 ± 0.28 | 52.14 ± 6.33 |
| T7opt | 2.08 ± 0.14 | 51.77 ± 3.19 |
| Tmax | 2.00 ± 0.03 | 50.01 ± 0.86 |
| T7wt | 1.44 ± 0.14 | 30.12 ± 6.63 |
| T41 | 1.44 ± 0.09 | 30.43 ± 4.18 |

**Supplementary Table 7**. Features of the seven terminators analyzed in the *in vitro* and *in vivo* assays. The predicted strength of the four terminators not present in the original dataset is also shown.

| | Terminators | | | | | | | |
|---|---|---|---|---|---|---|---|---|
| **Features** | **TK** | **MiniTK** | **T7wild** | **T7opt** | **T1** | **T41** | **T Max** | **B0010** |
| **Loop length** | 4 | 4 | 6 | 6 | 4 | 4 | 4 | 4 |
| **A% (first 6 nt of A-tract)** | 6 | 6 | 3 | 6 | 1 | 6 | 5 | 5 |
| **C% (first 6 nt of A-tract)** | 0 | 0 | 2 | 0 | 2 | 0 | 0 | 0 |
| **U% (first 10 nt of U-tract)** | 8 | 8 | 8 | 8 | 0 | 4 | 8 | 7 |
| **U% (first 6 nt of U-tract)** | 6 | 6 | 6 | 6 | 0 | 3 | 6 | 5 |
| **A% (first 6 nt of U-tract)** | 0 | 0 | 0 | 0 | 2 | 1 | 0 | 1 |
| **C% (entire U-tract)** | 1 | 1 | 1 | 1 | 1 | 5 | 1 | 1 |
| **Hairpin length (no loop)** | 48 | 20 | 26 | 26 | 18 | 14 | 16 | 28 |
| **Loop G/C content** | 1 | 1 | 2 | 2 | 1 | 1 | 1 | 1 |
| **Shannon Entropy (A-tract)** | 1.0613 | 1.0613 | 1.5613 | 1.0613 | 1.6645 | 1.0613 | 0.8113 | 0.5435 |
| **Shannon Entropy (U-tract)** | 0.8167 | 0.8167 | 1.4183 | 0.8167 | 0.9864 | 1.7842 | 0.8167 | 1.4183 |
| **Shannon Entropy (hairpin without loop)** | 1.7383 | 1.7383 | 1.6935 | 1.6935 | 1.9708 | 1.5917 | 1.2717 | 1.9627 |
| **State changes (A-tract)** | 2 | 2 | 4 | 2 | 6 | 2 | 4 | 2 |
| **State changes (U-tract)** | 3 | 3 | 3 | 3 | 7 | 7 | 3 | 7 |
| **State changes (hairpin without loop)** | 34 | 19 | 11 | 11 | 13 | 9 | 5 | 19 |
| **Initial GC pairs in hairpin** | 8 | 8 | 4 | 4 | 1 | 3 | 4 | 2 |
| **Predicted Strength** | 275.18 | 255.64 | 26.09 | 190.74 | | | | |

**Supplementary Table 8. Numerical values corresponding to the cross-species terminator predictions shown in Figure 3**.

Three datasets were evaluated. Kang et al.[28] characterized six terminators in *Corynebacterium glutamicum*. The authors reported terminator strengths on a 0–10 scale and ranked the terminators according to their activity, with T1 being the strongest and T6 the weakest, but did not report exact numerical values. Stukenberg et al.[29] characterized ten terminators in *Vibrio natriegens*. B0010 was excluded because it was present in the TerSP training dataset, and B0015 was excluded because it corresponds to a double terminator. Since detailed numerical values were not reported again, the experimental rank order was inferred from Figure 6D of the original study. Finally, Xu et al.[11] quantified the activity of 19 terminators in *Halomonas bluephagenesis*. Four terminators with negative reported termination efficiencies were excluded from the analysis. In addition, the synthetic terminator G04 was excluded because its U-tract was shortened to eight nucleotides, below the 12-nucleotide U-tract length used for TerSP analysis. For the remaining 14 terminators, the experimental TE values reported in the original study are shown together with the corresponding Ts values calculated using the equation described above. For all three datasets, TerSP-predicted TS and TE values are provided. Considering that complete quantitative data were not available for all datasets, the comparison was limited to a qualitative rank-order analysis. Terminators are listed in decreasing order of experimentally reported activity. Predicted values are shown in black when their relative order is consistent with the experimental ranking. Predicted values are shown in red when TerSP assigns a terminator a higher activity than one or more experimentally stronger terminators listed above it, indicating a rank-order inversion. NR means that numerical values were not reported by the original paper; only graphical results were presented.

| Terminator | Experimental TE | Experimental Ts | Predicted TE | Predicted Ts |
|---|---|---|---|---|
| ***Corynebacterium glutamicum*** | | | | |
| T1 | NR | NR | 95.1 | 20.39 |
| T2 | NR | NR | 94.2 | 17.21 |
| T3 | NR | NR | 92.4 | 13.24 |
| T4 | NR | NR | 90.4 | 10.47 |
| T5 | NR | NR | 88.2 | 8.44 |
| T6 | NR | NR | 90.1 | 10.07 |
| ***Vibrio natriegens*** | | | | |
| B1006 | NR | NR | 155.08 | 99.4 |
| B1002 | NR | NR | 23.37 | 95.7 |
| B1003 | NR | NR | 9.49 | 89.5 |
| B1007 | NR | NR | 33.54 | 97.0 |
| B1010 | NR | NR | 8.03 | 87.5 |
| B1005 | NR | NR | 4.57 | 78.1 |
| B1004 | NR | NR | 6.65 | 85.0 |
| B1009 | NR | NR | 10.52 | 90.5 |
| ***Halomonas bluephagenesis*** | | | | |
| A02 | 97.6 | 41.7 | 90.00 | 10.00 |
| A01 | 96.9 | 32.3 | 82.52 | 5.72 |
| D01 | 94.7 | 18.9 | 99.58 | 237.44 |
| D03 | 89.8 | 9.8 | 98.83 | 85.29 |
| D02 | 88.1 | 8.4 | 92.07 | 12.61 |
| A07 | 83.6 | 6.1 | 96.89 | 32.13 |

| | | | | |
|---|---|---|---|---|
| A09 | 51.4 | 2.1 | 81.41 | 5.38 |
| G01 | 50.1 | 2.0 | 96.26 | 26.72 |
| G02 | 39.1 | 1.6 | 96.26 | 26.72 |
| A10 | 36.5 | 1.6 | 95.05 | 20.19 |
| G03 | 32.8 | 1.5 | 93.50 | 15.39 |
| A06 | 27.9 | 1.4 | 81.06 | 5.28 |
| A03 | 24.4 | 1.3 | 77.58 | 4.46 |
| A05 | 16.6 | 1.2 | 81.06 | 5.28 |

**Supplementary Table 9.** Maximum-strength terminators for each hairpin length.

| Hairpin Length | Predicted Strength | Terminator RNA sequence: A-tract- 1st half of the hairpin-Loop- 2nd half of the hairpin- U-tract |
|---|---|---|
| 6 | 262.26 | AAAAAAAACACGAAAGUGUUUUUUCGUUUU |
| 8 | 264.10 | AAAAAAAACACAGAAAUGUGUUUUUUCGUUUU |
| 10 | 264.96 | AAAAAAAACAACAGAAAUGUUGUUUUUUCGUUUU |
| 12 | 298.70 | AAAAAAAACACACGGAAACGUGUGUUUUUUCGUUUU |
| 14 | 284.22 | AAAAAAAACAACACAGAAAUGUGUUGUUUUUUCGUUUU |
| 16 | 288.07 | AAAAAAAACAAACACA GAAAUGUGUUUGUUUUUUCGUUUU |
| 18 | 286.52 | AAAAAAAACAAAACACAGAAAUGUGUUUUGUUUUUUCGUUUU |
| 20 | 285.01 | AAAAAAAACAAAAACACGGAAACGUGUUUUUGUUUUUUCGUUUU |
| 22 | 285.16 | AAAAAAAACAAAAAACACGGAAACGUGUUUUUUGUUUUUUCGUUUU |
| 24 | 286.71 | AAAAAAAACAAAAAAACACGGAAACGUGUUUUUUUGUUUUUUCGUUUU |
| 26 | 286.73 | AAAAAAAACACACACGCGCGAAAGCGCGUGUGUGUGUUUUUUCGUUUU |
| 28 | 285.66 | AAAAAAAACACACACACACACAGAAAUGUGUGUGUGUGUGUUUUUUCGUUUU |
| 30 | 288.54 | AAAAAAAACACACACACACGCGCGAAAGCGCGUGUGUGUGUGUUUUUUCGUUUU |
| 32 | 289.14 | AAAAAAAACACACACACACGCGCGGAAACGCGCGUGUGUGUGUGUUUUUUCGUUUU |
| 34 | 288.14 | AAAAAAAACACACACACACACACACGAAAGUGUGUGUGUGUGUGUGUUUUUUCGUUUU |
| 36 | 290.96 | AAAAAAAACAACACACACACACACAUAUGAAAAUAUGUGUGUGUGUGUUGUUUUUUCGUUUU |
| 38 | 291.38 | AAAAAAAACAAACACACACACACACAUAUGAAAAUAUGUGUGUGUGUGUUUGUUUUUUCGUUUU |
| 40 | 290.00 | AAAAAAAACAAAACACACACACACACACAUGAAAAUGUGUGUGUGUGUGUUUUGUUUUUUCGUUUU |
| 42 | 293.70 | AAAAAAAACCCCACCGCGCGCGCGCGCGAAAGCGCGCGCGCGCGGUGGGGUUUUUUAGUUUU |
| 44 | 295.60 | AAAAAAAACCCCACCCGCGCGCGCGCGCGAAAGCGCGCGCGCGCGGGUGGGGUUUUUUAGUUUU |
| 46 | 297.10 | AAAAAAAACCCCACCCCGCGCGCGCGCGCGAAAGCGCGCGCGCGCGGGGUGGGGUUUUUUAGUUUU |
| 48 | 299.40 | AAAAAAAACCCCCCCCGACACACACACACACAGAAAUGUGUGUGUGUGUCGGGGGGGGUUUUUUCGUUUU |

**Supplementary Table 10.** List of oligonucleotides used in this work. The T7 promoter is highlighted in red.

| Name | Sequence |
|---|---|
| fw_T7 | GAAATTAATACGACTCACTATAGGGAGA |
| fw_T7_BabySpinach | GAAATTAATACGACTCACTATAGGGAGAGGTGAAGGACGGGTCCAGTAGTTCGCTACT GTTGAGTAGAGTGTGAGCTCC |
| rv_T7_BabySpinach | GGAGCTCACACTCTACTCAACAGTAGCGAACTACTGGACCCGTCCTTCACCTCT CCCTATAGTGAGTCGTATTAATTTC |
| fw_T7_Corn | GAAATTAATACGACTCACTATAGGGAGAGGCGCGAGGAAGGAGGTCTGAGG AGGTCACTGCGCC |
| rv_T7_Corn | GGCGCAGTGACCTCCTCAGACCTCCTTCCTCGCGCCTCTCCCTATAGTGAGTCG TATTAATTTC |
| fw_T7_Broccoli | GAAATTAATACGACTCACTATAGGGAGAGAGACGGTCGGGTCCAGATATTCG TATCTGTCGAGTAGAGTGTGGGCTC |
| rv_T7_Broccoli | GAGCCCACACTCTACTCGACAGATACGAATATCTGGACCCGACCGTCTCTCTCC CTATAGTGAGTCGTATTAATTTC |
| fw_T7_Mango_II | GAAATTAATACGACTCACTATAGGGAGAGGCACGTACGAAGGAGAGGAGAG GAAGAGGAGAGTACGTGC |
| rv_T7_Mango_II | GCACGTACTCTCCTCTTCCTCTCCTCTCCTTCGTACGTGCCTCTCCCTATAGTGA GTCGTATTAATTTC |
| fw_T7_Mango_III | GAAATTAATACGACTCACTATAGGGAGAGGCACGTACGAAGGAAGGTTTGGT ATGTGGTATATTCGTACGTGC |
| rv_T7_Mango_III | GCACGTACGAATATACCACATACCAAACCTTCCTTCGTACGTGCCTCTCCCTAT AGTGAGTCGTATTAATTTC |

**Supplementary Table 11.** List of gene fragments used in this work. The T7 promoter is highlighted in red, the terminators in blue, Broccoli in green, and Mango III in yellow.

| Name | Sequence |
|---|---|
| T7_Broccoli_T1_Mango_III | GAAATTAATACGACTCACTATAGGGAGAGCCCGGATAGCTCAGTCGGTAGAGCAGCGGCCGGAGACGGTCGGGTCCAGATATTCGTATCTGTCGAGTAGAGTGTGGGCTCCGGCCGCGGGTCCAGGGTTCAAGTCCCTGTTCGGGCGCCAGTTGCCATGTGTATGTGGGTTCGCCCACATACTCTGATGATCCCCACTGCTCTGACCACAAGTAATTGTTCAGATTGATAAAACAAAGCTTCCCGGGAAAGTATATATGAGTAAAGATATCGGCACGTACGAAGGAAGGTTTGGTATGTGGTATATTCGTACGTGCGATATCCCCGGGCTAGCA |
| T7_ Mango_III _T1_ Broccoli | ACCACTGCTGAAATTAATACGACTCACTATAGGGAGAGCCCGGATAGCTCAGTCGGTAGAGCAGCGGCCGGGCACGTACGAAGGAAGGTTTGGTATGTGGTATATTCGTACGTGCCGGCCGCGGGTCCAGGGTTCAAGTCCCTGTTCGGGCGCCACCACTGCTCTGACCACAAGTAATTGTTCAGATTGATAAAACAAAGCTTCCCGGGAAAGTATATATGAGTAAAGATATCGAGACGGTCGGGTCCAGATATTCGTATCTGTCGAGTAGAGTGTGGGCTCGATATCCCCGGGCTAGCA |
| T7_Broccoli_T41_Mango_III | GTAAACCACTGCTGAAATTAATACGACTCACTATAGGGAGAGCCCGGATAGCTCAGTCGGTAGAGCAGCGGCCGGAGACGGTCGGGTCCAGATATTCGTATCTGTCGAGTAGAGTGTGGGCTCCGGCCGCGGGTCCAGGGTTCAAGTCCCTGTTCGGGCGCCAGTAAAAAAGGCAGCCATCTGGCTGCCTTAGTCTCCCCAAAGCTTCCCGGGAAAGTATATATGAGTAAAGATATCGGCACGTACGAAGGAAGGTTTGGTATGTGGTATATTCGTACGTGCGATATCCCCGGGCTAGCA |
| T7_ Mango_III _T41_ Broccoli | GTAAACCACTGCTGAAATTAATACGACTCACTATAGGGAGAGCCCGGATAGCTCAGTCGGTAGAGCAGCGGCCGGGCACGTACGAAGGAAGGTTTGGTATGTGGTATATTCGTACGTGCCGGCCGCGGGTCCAGGGTTCAAGTCCCTGTTCGGGCGCCAGTAAAAAAGGCAGCCATCTGGCTGCCTTAGTCTCCCCAAAGCTTCCCGGGAAAGTATATATGAGTAAAGATATCGAGACGGTCGGGTCCAGATATTCGTATCTGTCGAGTAGAGTGTGGGCTCGATATCCCCGGGCTAGCA |
| T7_Broccoli_Tmax_Mango_III | ATACCACTGCTGAAATTAATACGACTCACTATAGGGAGAGCCCGGATAGCTCAGTCGGTAGAGCAGCGGCCGGAGACGGTCGGGTCCAGATATTCGTATCTGTCGAGTAGAGTGTGGGCTCCGGCCGCGGGTCCAGGGTTCAAGTCCCTGTTCGGGCGCCAAGAAAAGAGGCCTCCCGAAAGGGGGGCCTTTTTTCGTTTTAAGCTTCCCGGGAAAGTATATATGAGTAAAGATATCGGCACGTACGAAGGAAGGTTTGGTATGTGGTATATTCGTACGTGCGATATCCCCGGGCTAGCA |
| T7_ Mango_III _Tmax_ Broccoli | TACCACTGCTGAAATTAATACGACTCACTATAGGGAGAGCCCGGATAGCTCAGTCGGTAGAGCAGCGGCCGGGCACGTACGAAGGAAGGTTTGGTATGTGGTATATTCGTACGTGCCGGCCGCGGGTCCAGGGTTCAAGTCCCTGTTCGGGCGCCAAGAAAAGAGGCCTCCCGAAAGGGGGGCCTTTTTTCGTTTTAAGCTTCCCGGGAAAGTATATATGAGTAAAGATATCGAGACGGTCGGGTCCAGATATTCGTATCTGTCGAGTAGAGTGTGGGCTCGATATCCCCGGGCTAGCA |
| T7_Broccoli_T7wild_Mango_III | ACCACTGCTGAAATTAATACGACTCACTATAGGGAGAGCCCGGATAGCTCAGTCGGTAGAGCAGCGGCCGGAGACGGTCGGGTCCAGATATTCGTATCTGTCGAGTAGAGTGTGGGCTCCGGCCGCGGGTCCAGGGTTCAAGTCCCTGTTCGGGCGCCAAACCCCTTGGGGCCTCTAAACGGGTCTTGAGGGGTTTTTTTTAAGCTTCCCGGGAAAGTATATATGAGTAAAGATATCGGCACGTACGAAGGAAGGTTTGGTATGTGGTATATTCGTACGTGCGATATCCCCGGGCTAGCA |

| | |
|---|---|
| T7_ Mango_III_T7wild_ Broccoli | accactgcTGAAATTAATACGACTCACTATAGGGAGAGCCCGGATAGCTCAGTCGGTAGAGCAGCGGCCGGGCACGTACGAAGGAAGGTTTGGTATGTGGTATATTCGTACGTGCCGGCCGCGGGTCCAGGGTTCAAGTCCCTGTTCGGGCGCCAAACCCCTTGGGGCCTCTAAACGGGTCTTGAGGGGTTTTTTTTAAGCTTCCCGGGAAAGTATATATGAGTAAAGATATCGAGACGGTCGGGTCCAGATATTCGTATCTGTCGAGTAGAGTGTGGGCTCGATATCCCCGGGCTAGCA |
| T7_Broccoli_T7opt_Mango_III | GAAATTAATACGACTCACTATAGGGAGAGCCCGGATAGCTCAGTCGGTAGAGCAGCGGCCGGAGACGGTCGGGTCCAGATATTCGTATCTGTCGAGTAGAGTGTGGGCTCCGGCCGCGGGTCCAGGGTTCAAGTCCCTGTTCGGGCGCCAGTAAAAAACCCCTTGGGGCCTCTAAACGGGTCTTGAGGGGTTTTTTTTCATTAAGCTTCCCGGGAAAGTATATATGAGTAAAGATATCGGCACGTACGAAGGAAGGTTTGGTATGTGGTATATTCGTACGTGCGATATCCCCGGGCTAGCA |
| T7_ Mango_III_T7opt_ Broccoli | GAAATTAATACGACTCACTATAGGGAGAGCCCGGATAGCTCAGTCGGTAGAGCAGCGGCCGGGCACGTACGAAGGAAGGTTTGGTATGTGGTATATTCGTACGTGCCGGCCGCGGGTCCAGGGTTCAAGTCCCTGTTCGGGCGCCAGTAAAAAACCCCTTGGGGCCTCTAAACGGGTCTTGAGGGGTTTTTTTTCATTAAGCTTCCCGGGAAAGTATATATGAGTAAAGATATCGAGACGGTCGGGTCCAGATATTCGTATCTGTCGAGTAGAGTGTGGGCTCGATATCCCCGGGCTAGCA |
| T7_Broccoli_TK_Mango_III | GAAATTAATACGACTCACTATAGGGAGAGCCCGGATAGCTCAGTCGGTAGAGCAGCGGCCGGAGACGGTCGGGTCCAGATATTCGTATCTGTCGAGTAGAGTGTGGGCTCCGGCCGCGGGTCCAGGGTTCAAGTCCCTGTTCGGGCGCCACGAAAAAACCGCGCCGAGGCCGGCGCCGATATACAAATATCGGCGCCGGCCTCGGCGCGGTTTTTTTTCATTAAGCTTCCCGGGAAAGTATATATGAGTAAAGATATCGGCACGTACGAAGGAAGGTTTGGTATGTGGTATATTCGTACGTGCGATATCCCCGGGCTAGCA |
| T7_ Mango_III_TK_ Broccoli | GAAATTAATACGACTCACTATAGGGAGAGCCCGGATAGCTCAGTCGGTAGAGCAGCGGCCGGGCACGTACGAAGGAAGGTTTGGTATGTGGTATATTCGTACGTGCCGGCCGCGGGTCCAGGGTTCAAGTCCCTGTTCGGGCGCCACGAAAAAACCGCGCCGAGGCCGGCGCCGATATACAAATATCGGCGCCGGCCTCGGCGCGGTTTTTTTTCATTAAGCTTCCCGGGAAAGTATATATGAGTAAAGATATCGAGACGGTCGGGTCCAGATATTCGTATCTGTCGAGTAGAGTGTGGGCTCGATATCCCCGGGCTAGCA |
| T7_Broccoli_MiniTK_Mango_III | cactgcTGAAATTAATACGACTCACTATAGGGAGAGCCCGGATAGCTCAGTCGGTAGAGCAGCGGCCGGAGACGGTCGGGTCCAGATATTCGTATCTGTCGAGTAGAGTGTGGGCTCCGGCCGCGGGTCCAGGGTTCAAGTCCCTGTTCGGGCGCCACGAAAAAACGCGCGCGATACAAATCGCGCGCGTTTTTTTTCATTAAGCTTCCCGGGAAAGTATATATGAGTAAAGATATCGGCACGTACGAAGGAAGGTTTGGTATGTGGTATATTCGTACGTGCGATATCCCCGGGCTAGCA |
| T7_ Mango_III_MiniTK_ Broccoli | cactgcTGAAATTAATACGACTCACTATAGGGAGAGCCCGGATAGCTCAGTCGGTAGAGCAGCGGCCGGGCACGTACGAAGGAAGGTTTGGTATGTGGTATATTCGTACGTGCCGGCCGCGGGTCCAGGGTTCAAGTCCCTGTTCGGGCGCCACGAAAAAACGCGCGCGATACAAATCGCGCGCGTTTTTTTTCATTAAGCTTCCCGGGAAAGTATATATGAGTAAAGATATCGAGACGGTCGGGTCCAGATATTCGTATCTGTCGAGTAGAGTGTGGGCTCGATATCCCCGGGCTAGCA |

**Supplementary Figure 1. Sequential selection of sequence-derived features for terminator strength prediction.** All candidate features listed in Supplementary Table 5 were evaluated during feature selection. The figure shows only the retained feature set, corresponding to the 16 features that improved test-set $R^2$ and were therefore used for final model training. Candidate features that did not improve test-set performance were omitted from the model and are not shown here for simplicity. Each panel represents one step in the feature-selection process, in which one or more features were added to the cumulative feature set and the model was retrained. The left plot in each panel shows the experimental average strength values from the Chen et al. dataset and is repeated across panels as a fixed reference. The right plot shows the corresponding model predictions obtained using the cumulative feature set indicated below that panel, with the train and test $R^2$ values reported inside the plot. The x-axis labelled “Number” corresponds to the index assigned to each terminator after sorting the dataset from weakest to strongest according to experimental average strength. The red dashed horizontal line marks the strength threshold of 40 used to distinguish weak and strong terminators. The blue dashed vertical line marks the position in the sorted experimental dataset at which this threshold is crossed. Thus, in the prediction plots, points in the lower-left and upper-right quadrants are classified consistent with the experimental weak/strong threshold, whereas points in the upper-left and lower-right quadrants correspond to misclassified terminators. The feature labels below the plots indicate the sequence features added at each selection step and their corresponding terminator segment. The segment labels A-tract, Hairpin, Loop, and U-tract refer to the four terminator regions originally defined by Chen et al. and illustrated in Figure 1a.

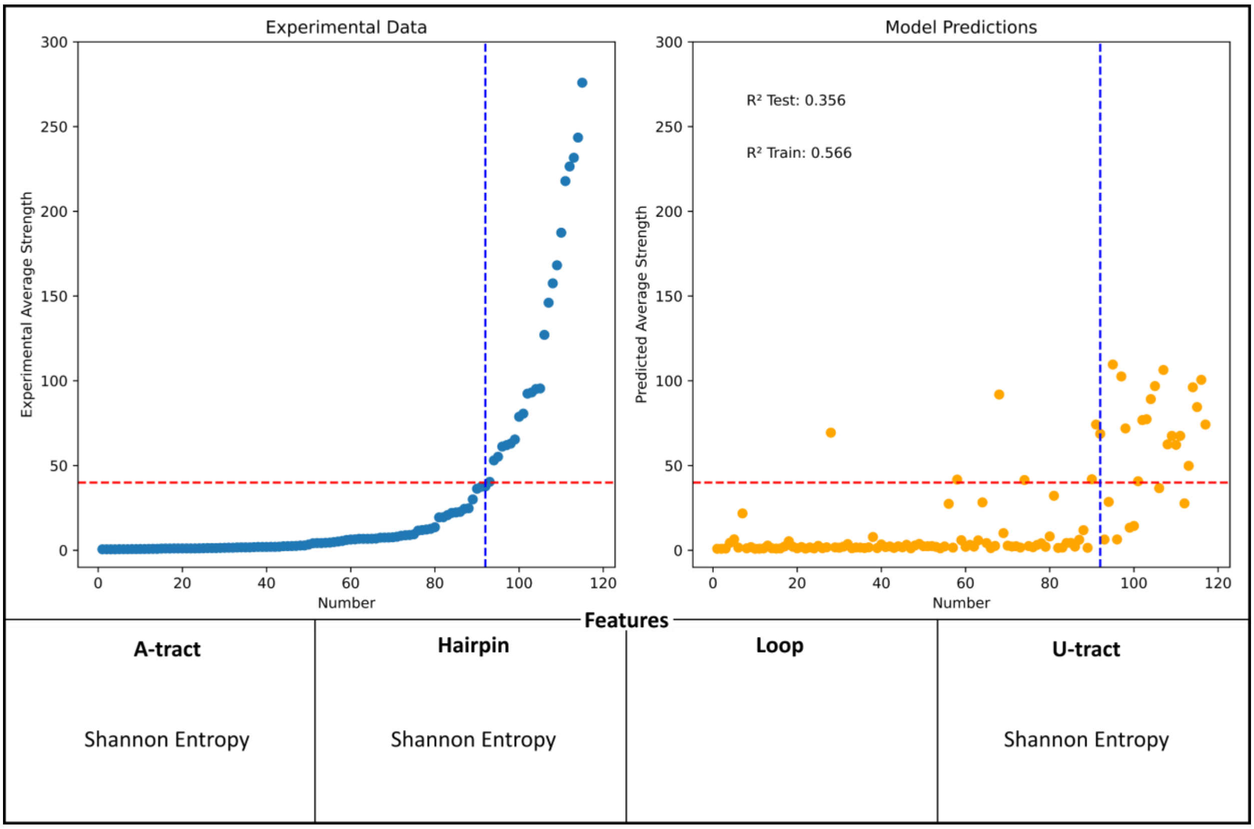

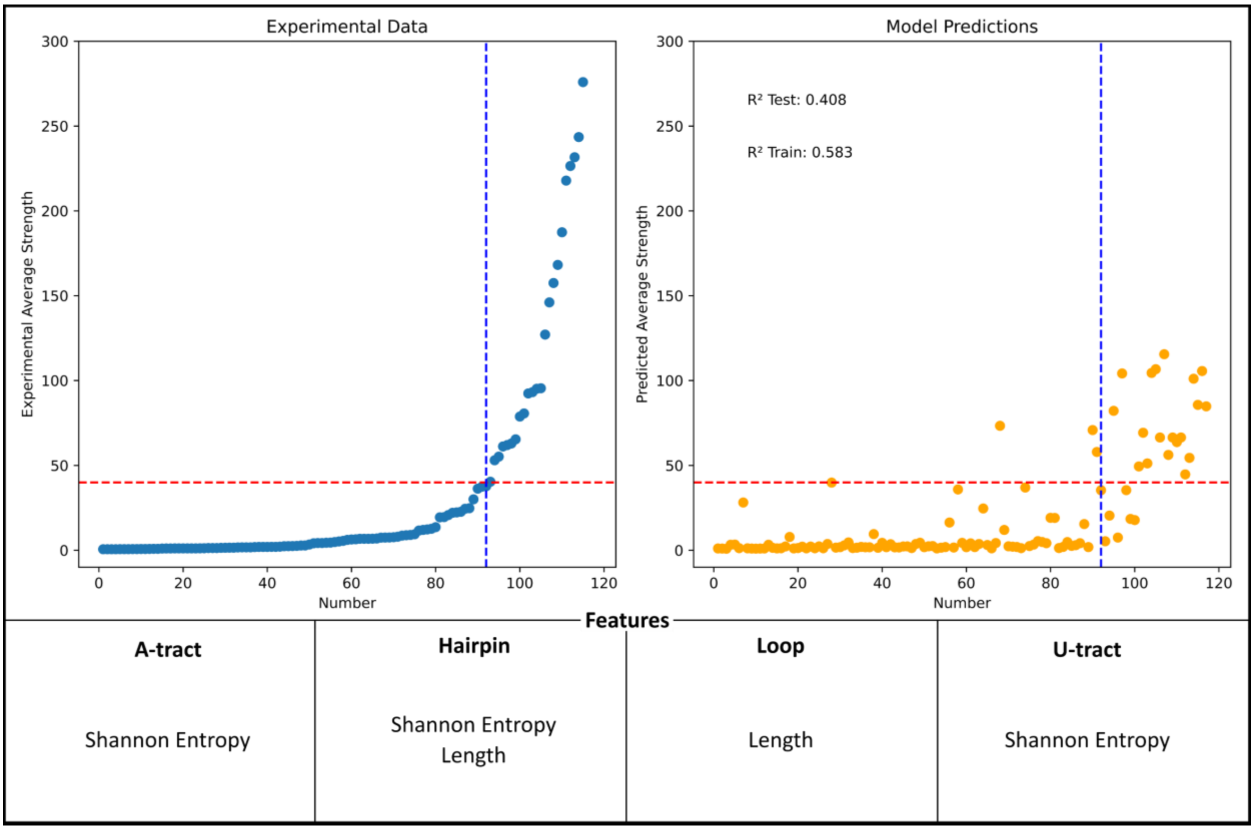
Experimental Data
Experimental Average Strength
Number
Model Predictions
R² Test: 0.408
R² Train: 0.583
Predicted Average Strength
Number
Features
A-tract
Shannon Entropy
Hairpin
Shannon Entropy
Length
Loop
Length
U-tract
Shannon Entropy

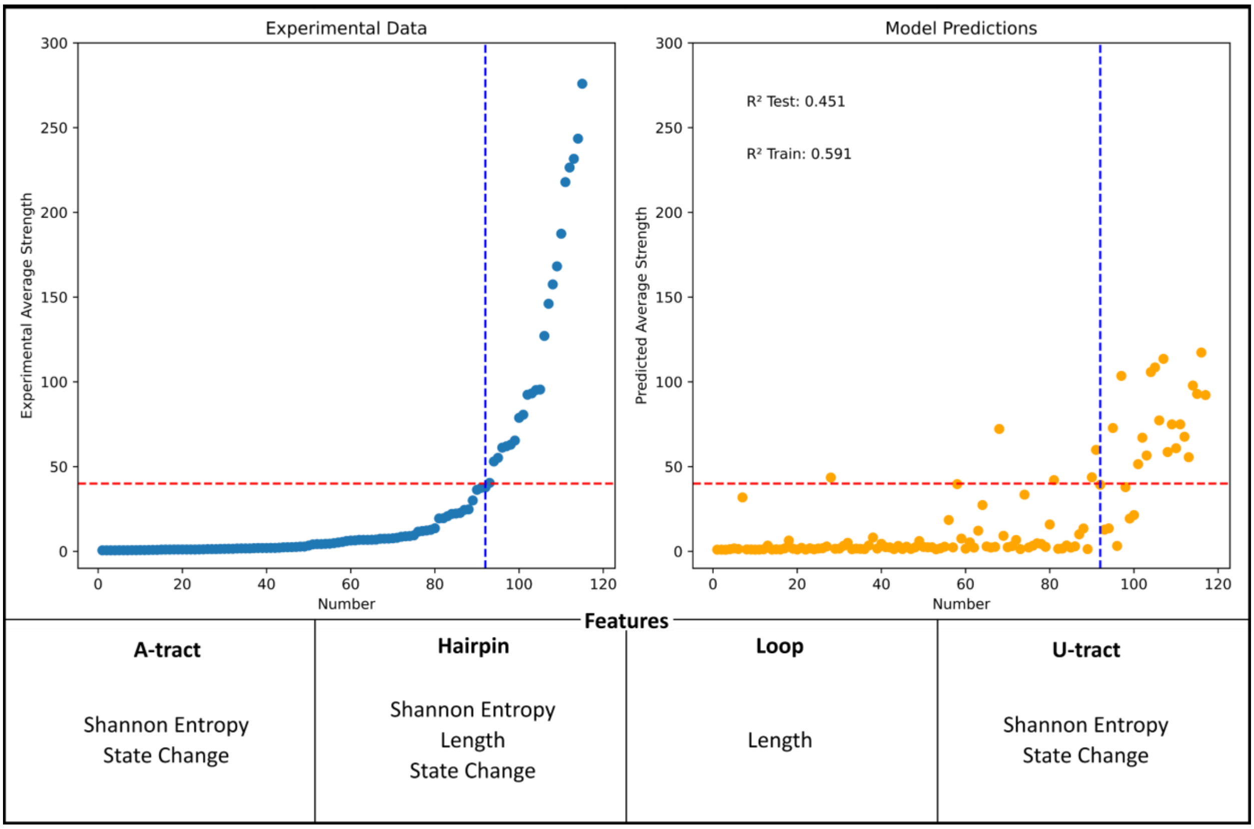
Experimental Data
Experimental Average Strength
Number
Model Predictions
R² Test: 0.451
R² Train: 0.591
Predicted Average Strength
Number
Features
A-tract
Shannon Entropy
State Change
Hairpin
Shannon Entropy
Length
State Change
Loop
Length
U-tract
Shannon Entropy
State Change

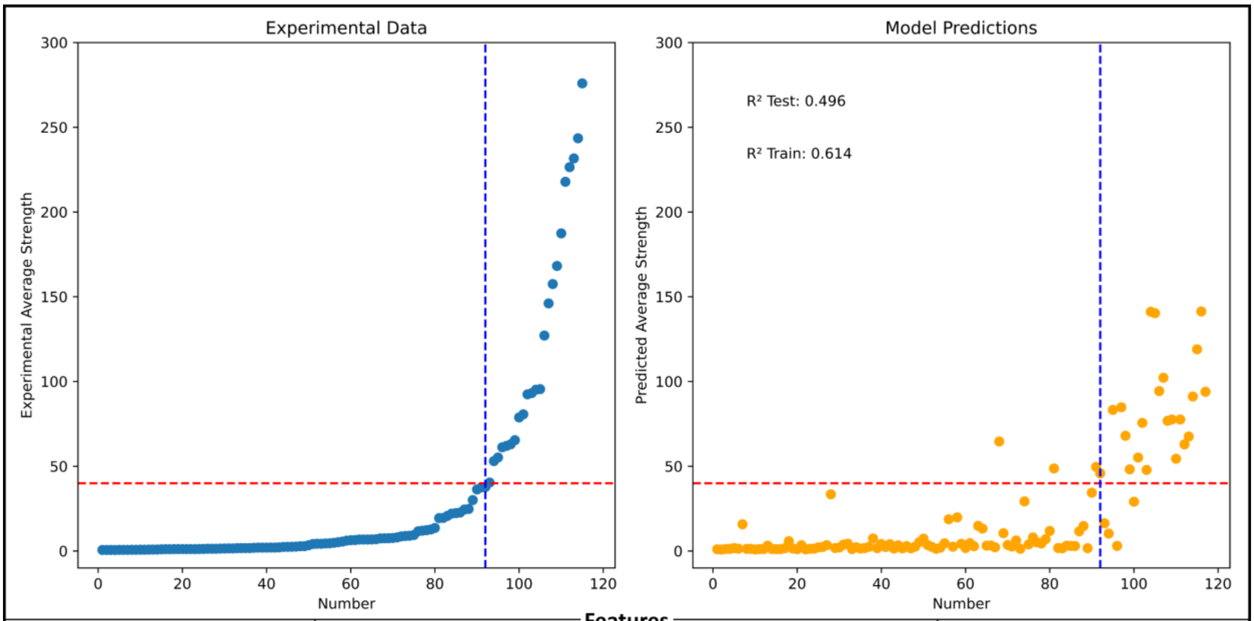


**Features**

| A-tract | Hairpin | Loop | U-tract |
| --- | --- | --- | --- |
| Shannon Entropy<br>State Change | Shannon Entropy<br>Length<br>State Change<br>Initial G/C | Length<br>%GC | Shannon Entropy<br>State Change |

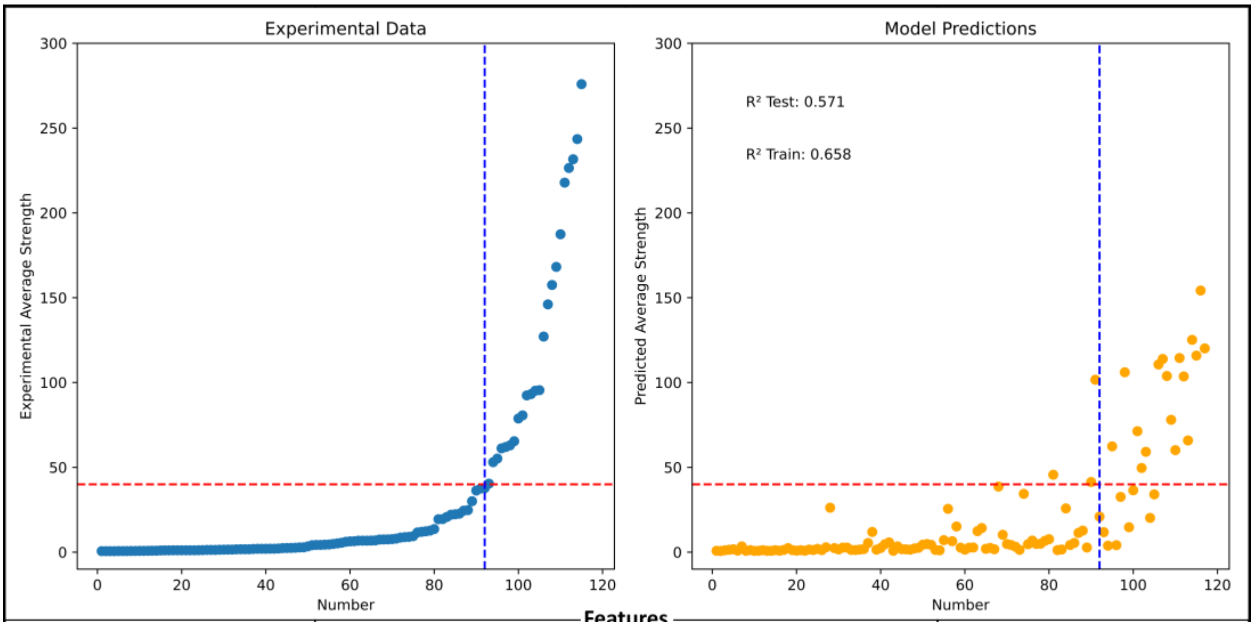


**Features**

| A-tract | Hairpin | Loop | U-tract |
| --- | --- | --- | --- |
| Shannon Entropy<br>State Change<br>A%-6<br>C%-6 | Shannon Entropy<br>Length<br>State Change<br>Initial G/C | Length<br>%GC | Shannon Entropy<br>State Change<br>A%-6<br>U%-6<br>U%-10<br>C%-12 |

**Supplementary Figure 2. Complete hyperparameter optimization landscape of the XGBoost model.** Expanded version of the summarized version presented in Figure 1b. The heatmap shows the cross-validation (CV) mean test $R^2$ obtained for each evaluated XGBoost hyperparameter combination. Columns represent combinations of the number of estimators (n) and learning rate (lr), while rows represent combinations of maximum tree depth (d), minimum child weight (cw), and subsample fraction (sub). Color intensity indicates the mean test $R^2$ across folds. Hatched cells indicate configurations discarded from consideration because the train–test $R^2$ gap exceeded 0.3 or the training $R^2$ exceeded 0.9, consistent with the overfitting filters applied during model selection. The selected configuration is highlighted by the rectangle and corresponds to n_estimators = 4000, learning_rate = 0.001, max_depth = 5, min_child_weight = 3, and subsample = 0.65.

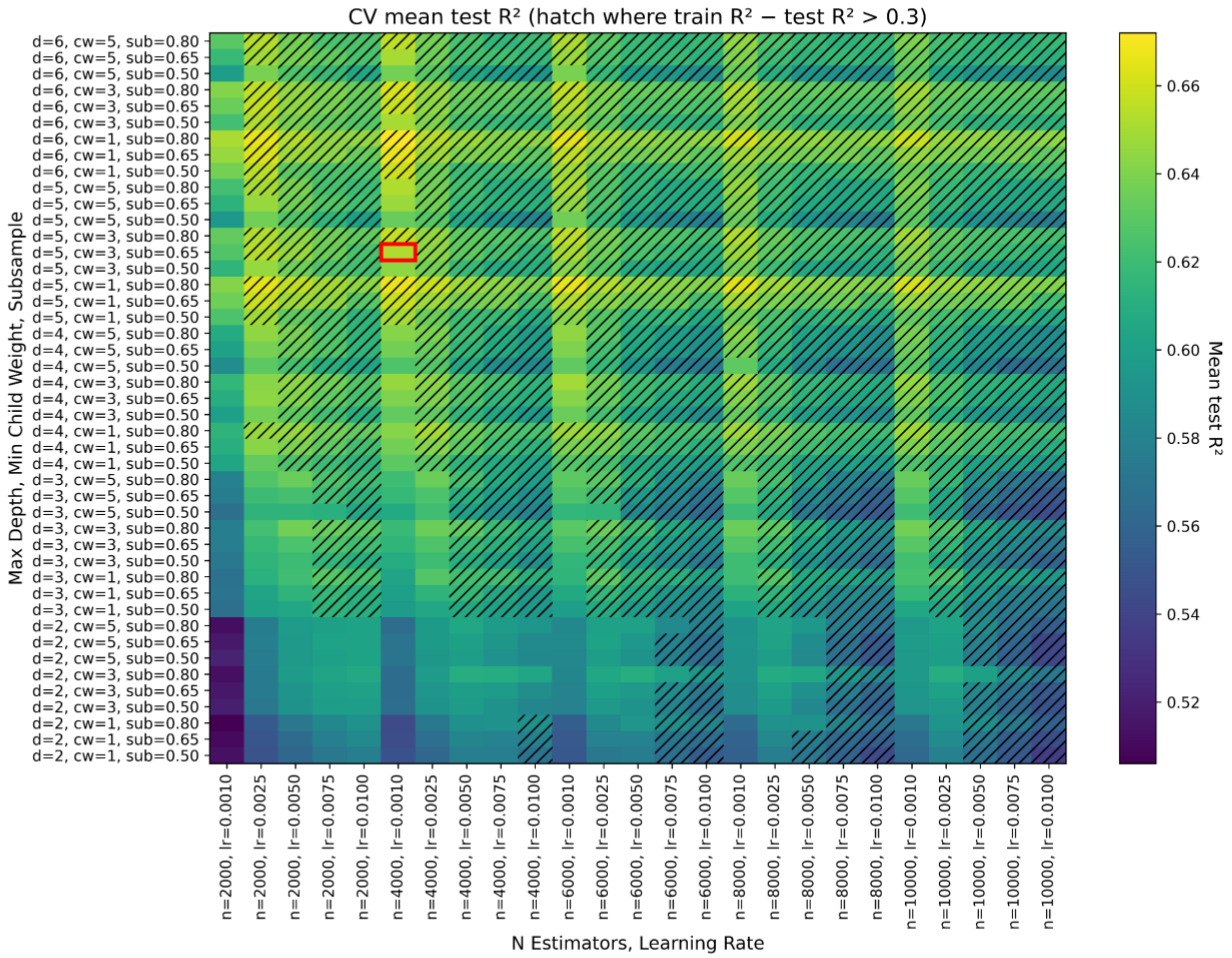

**Supplementary Figure 3.** Predictions obtained with the biophysical model reported by Chen et al. Experimental average strengths from the Chen et al. dataset were compared with the strengths predicted by their biophysical model, defined as: $T_S = 1 + \frac{1}{B_1 e^{\beta_1 \Delta G_L} + B_4 e^{\beta_4 (\Delta G_B + \Delta G_A - \Delta G_U)} (1 + B_1 e^{\beta_1 \Delta G_A})}$

where B1 = 0.005, B4 = 6.0, β1 = 0.6, and β4 = 0.45 are fit parameters. Terminators are ordered from weakest to strongest according to experimental average strength. The left panel shows the experimental values, and the right panel shows the corresponding predictions from the Chen et al. model in the same order. The red dashed line indicates the strength threshold of 40 used to distinguish weak and strong terminators. In the original publication, Chen et al. reported an $R^2$ of 0.40 for their biophysical model using the log-scale measured-versus-predicted comparison shown in their Figure 4a. For consistency with the evaluation of our machine-learning models, $R^2$ was recalculated on the untransformed average strength scale.

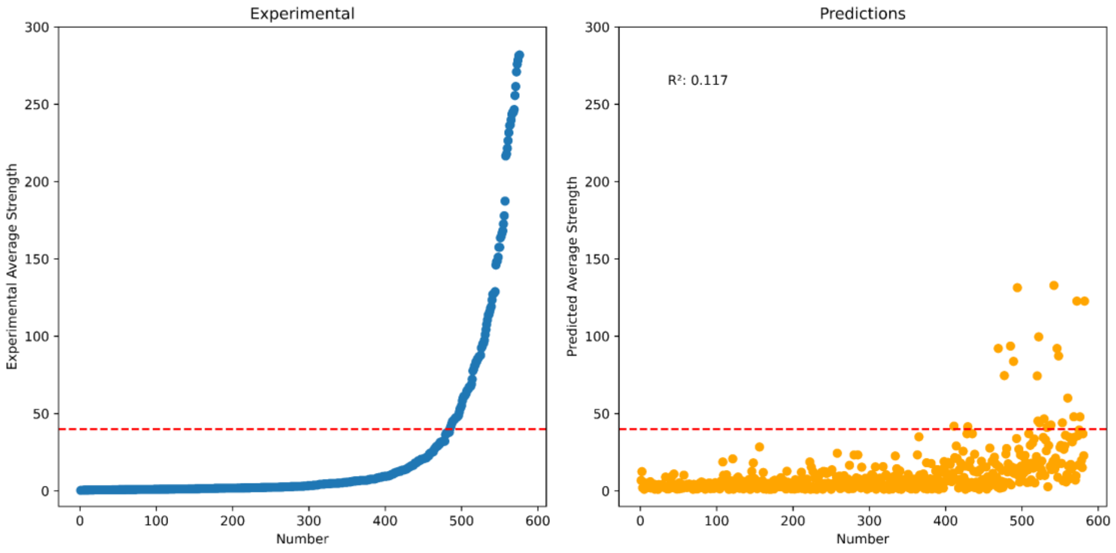

**Supplementary Figure 4. Performance of linear regression under five-fold cross-validation.** Linear regression was evaluated using the selected sequence-derived features under five-fold cross-validation. In each fold, experimental terminator strengths in the test set are shown on the left, ordered from weakest to strongest, and the corresponding linear regression predictions are shown on the right using the same order. The red dashed horizontal line marks the strength threshold of 40 used to distinguish weak and strong terminators. Across the five folds, linear regression achieved an overall mean train $R^2$ of 0.404 and an overall mean test $R^2$ of 0.363. This limited predictive performance indicates that a linear model is insufficient to fully capture the relationship between terminator sequence features and measured strength, supporting the use of nonlinear machine-learning approaches.

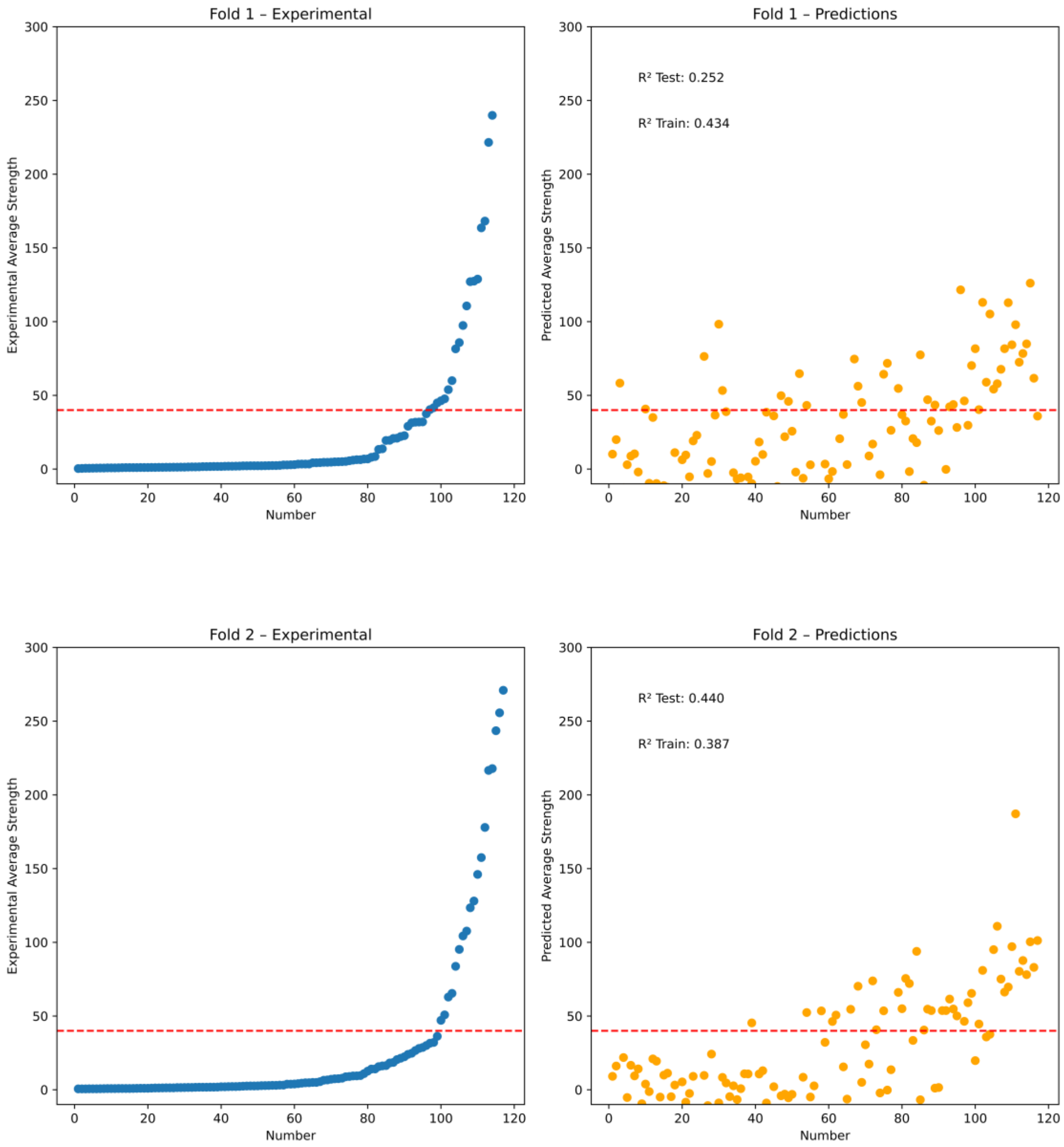

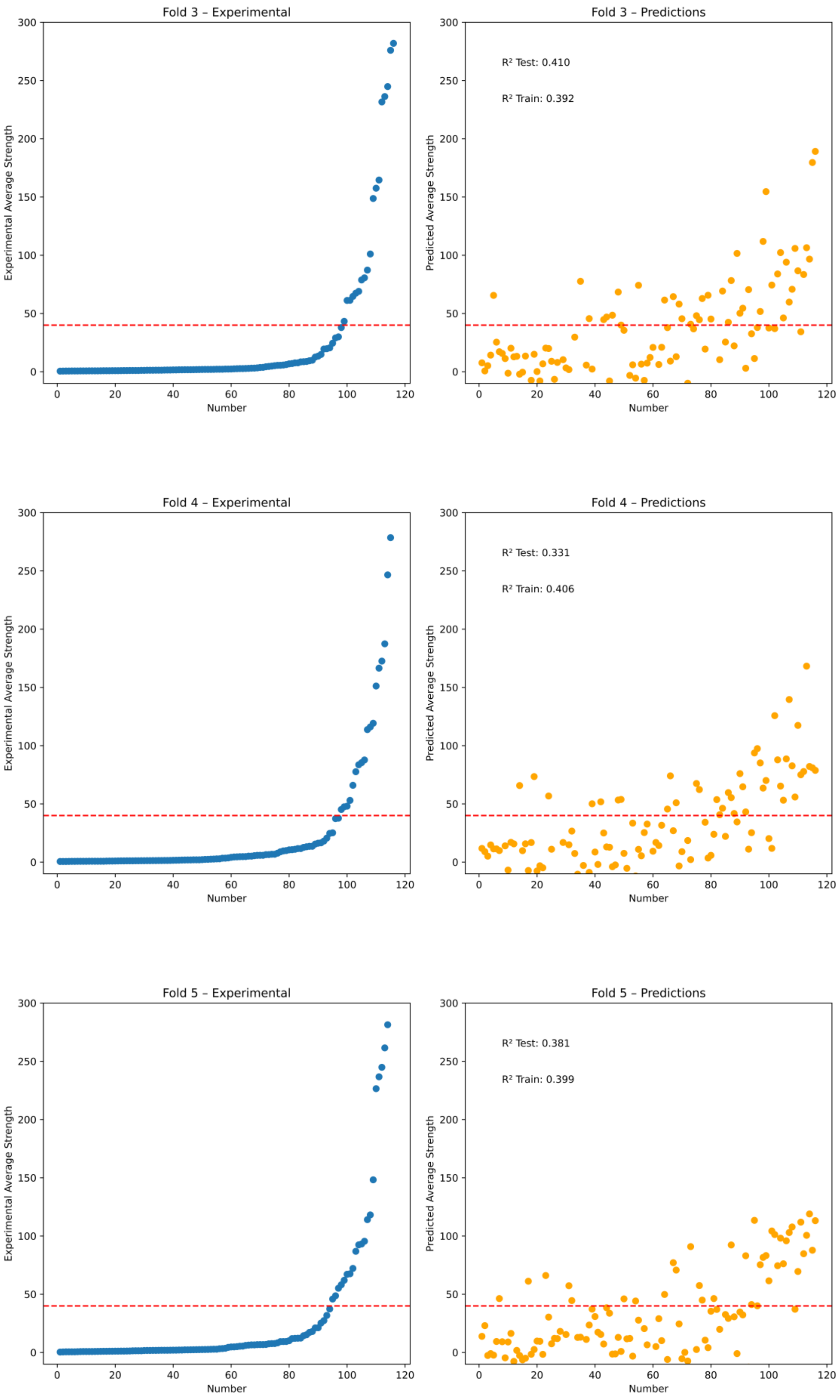
Fold 3 – Experimental
Experimental Average Strength
Number
Fold 3 – Predictions
R² Test: 0.410
R² Train: 0.392
Predicted Average Strength
Number
Fold 4 – Experimental
Experimental Average Strength
Number
Fold 4 – Predictions
R² Test: 0.331
R² Train: 0.406
Predicted Average Strength
Number
Fold 5 – Experimental
Experimental Average Strength
Number
Fold 5 – Predictions
R² Test: 0.381
R² Train: 0.399
Predicted Average Strength
Number

**Supplementary Figure 5. Evaluation of different MLPRegressor architectures.** Mean train and test $R^2$ values obtained under five-fold cross-validation for different MLPRegressor hidden-layer architectures. Architecture labels indicate the number of neurons in each hidden layer; for example, 10:20:10 corresponds to three hidden layers containing 10, 20 and 10 neurons. Bars represent mean $R^2$ across folds, and error bars indicate variability among folds. Although the 50:100:100:50 architecture showed the highest mean test $R^2$, its improvement over 10:20:10 was small (0.501 versus 0.490) and was accompanied by a larger train–test gap (0.173 versus 0.086), indicating that the additional complexity did not provide a clear gain in generalization. Conversely, the single-layer 10-neuron architecture showed a smaller train–test gap but slightly lower test performance, suggesting limited model flexibility. The 10:20:10 architecture was therefore selected for subsequent analyses as a balanced configuration, providing near-maximal cross-validated test performance with moderate complexity and a smaller train–test discrepancy than larger architectures.

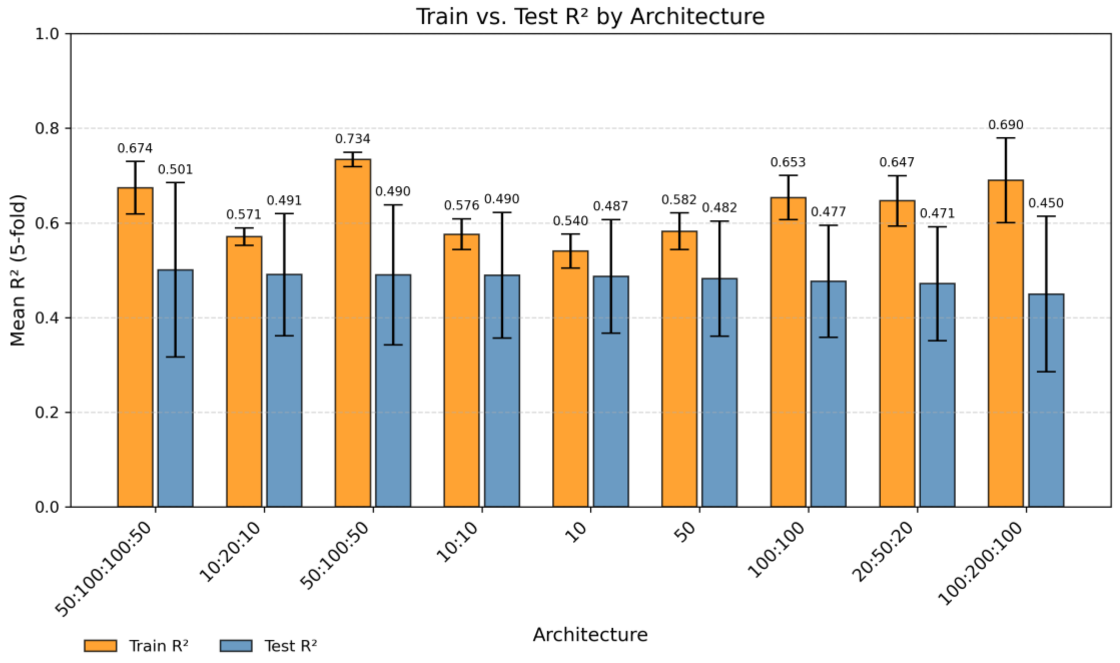

**Supplementary Figure 6. Evaluation of weighted XGBoost–MLPRegressor model combinations.** Mean train and test $R^2$ values obtained under five-fold cross-validation for XGBoost, MLPRegressor and weighted combinations of both models. Model-combination labels indicate the relative contribution of each model to the final prediction; for example, XGBoost 10:1 MLPRegressor corresponds to a weighted ensemble in which XGBoost contributes ten times more than MLPRegressor, whereas XGBoost 1:0 MLPRegressor and XGBoost 0:1 MLPRegressor correspond to XGBoost alone and MLPRegressor alone, respectively. Bars represent mean $R^2$ across folds, and error bars indicate variability among folds.

Weighted combinations were tested because XGBoost and MLPRegressor represent different nonlinear modeling strategies and could, in principle, capture complementary aspects of the relationship between sequence-derived features and terminator strength. However, XGBoost alone achieved the highest mean test $R^2$ (0.652), whereas MLPRegressor alone showed lower performance (test $R^2$ = 0.491). Adding MLPRegressor predictions to XGBoost did not improve test-set performance: combinations strongly weighted toward XGBoost produced similar but slightly lower test $R^2$ values, while increasing the MLPRegressor contribution progressively reduced performance. This suggests that, under the tested conditions, the MLPRegressor did not provide complementary predictive information beyond that already captured by XGBoost. Instead, averaging with the weaker model likely diluted the XGBoost predictions, supporting the selection of XGBoost alone as the final model.

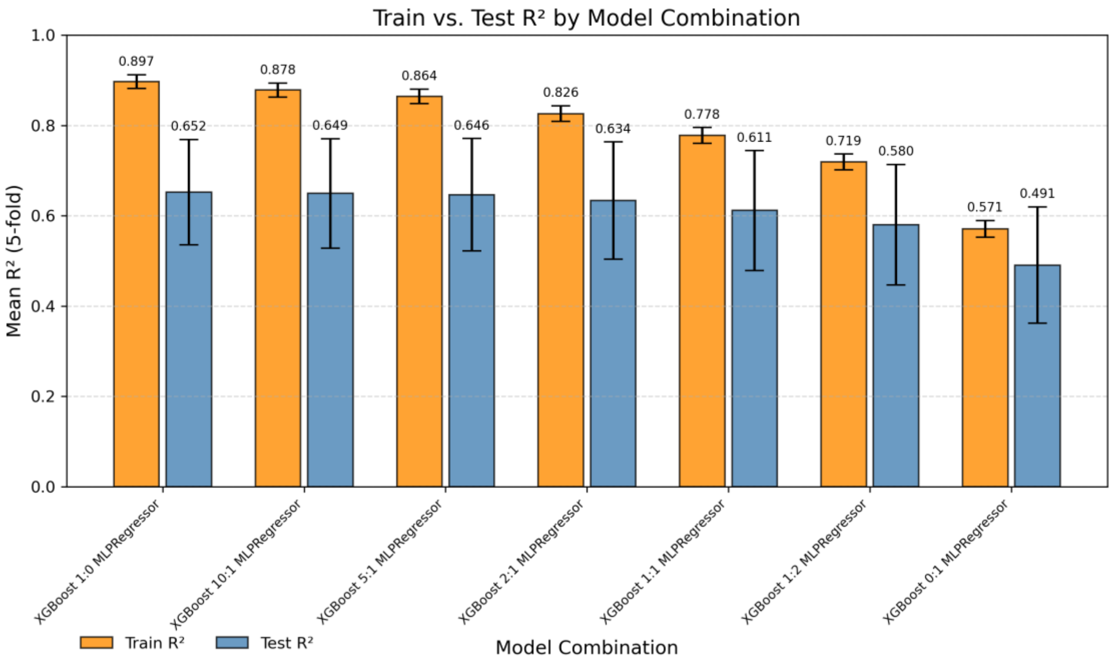

**Supplementary Figure 7. Reporter-only crosstalk controls in the Broccoli detection channel.** Reporter-only controls were used to determine whether reporter–fluorophore combinations produced a signal under the same acquisition settings used within the two reporter detection windows. In the Broccoli detection channel (excitation at 470 nm, emission at 505 nm), Broccoli–BI generated a strong fluorescence signal, whereas Mango III–BI and Broccoli–TO1 remained near background. Mango III–TO1 produced a smaller but measurable signal, indicating partial spectral spillover. To correct for this interference, the spillover was quantified using a calibration curve and subtracted from the Broccoli readout in the final dual-FLAP analysis. Reciprocal measurements using the Mango III detection settings showed no detectable signal from Mango III–BI, Broccoli–BI, or Broccoli–TO1, indicating negligible crosstalk into the Mango III channel under the selected assay conditions.

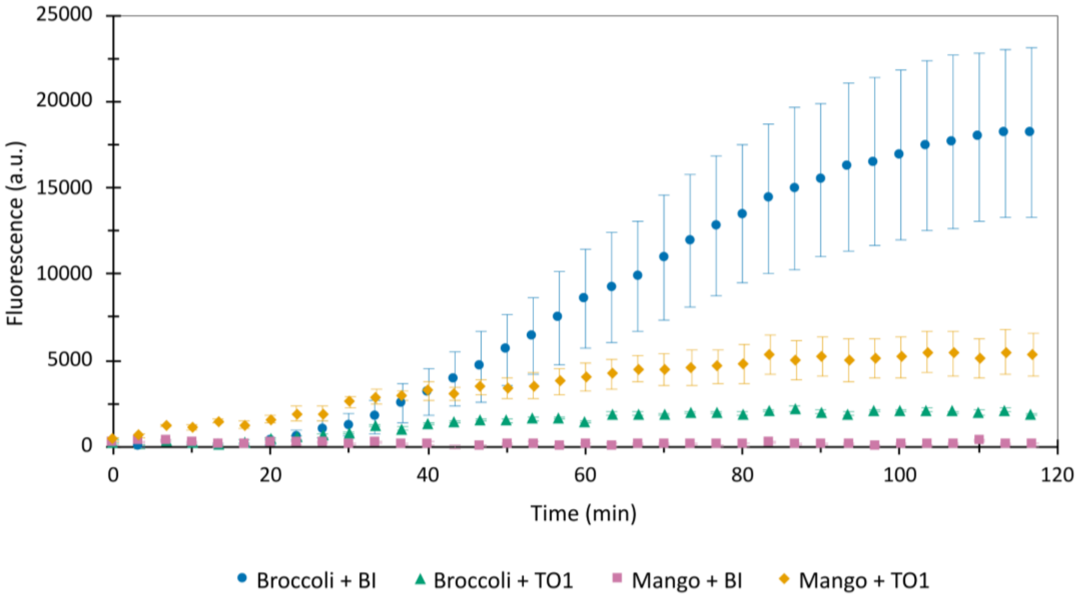

**Supplementary Figure 8. Effect of DNA template configuration on Broccoli fluorescence during *in vitro* transcription.** Broccoli fluorescence (excitation at 470 nm, emission at 505 nm) was monitored in real time from either a fully double-stranded DNA template or a hybrid single-stranded DNA template. In the hybrid template, the Broccoli-coding region remained single-stranded, while a short complementary oligonucleotide rendered the T7 promoter region double-stranded. Although this template format has previously been reported to support real-time Broccoli-based monitoring of T7 transcription[30], it performed less efficiently under our assay conditions, producing delayed signal accumulation and substantially lower fluorescence than the fully double-stranded template. These results supported the use of fully double-stranded DNA templates in the final dual-FLAP termination assay.

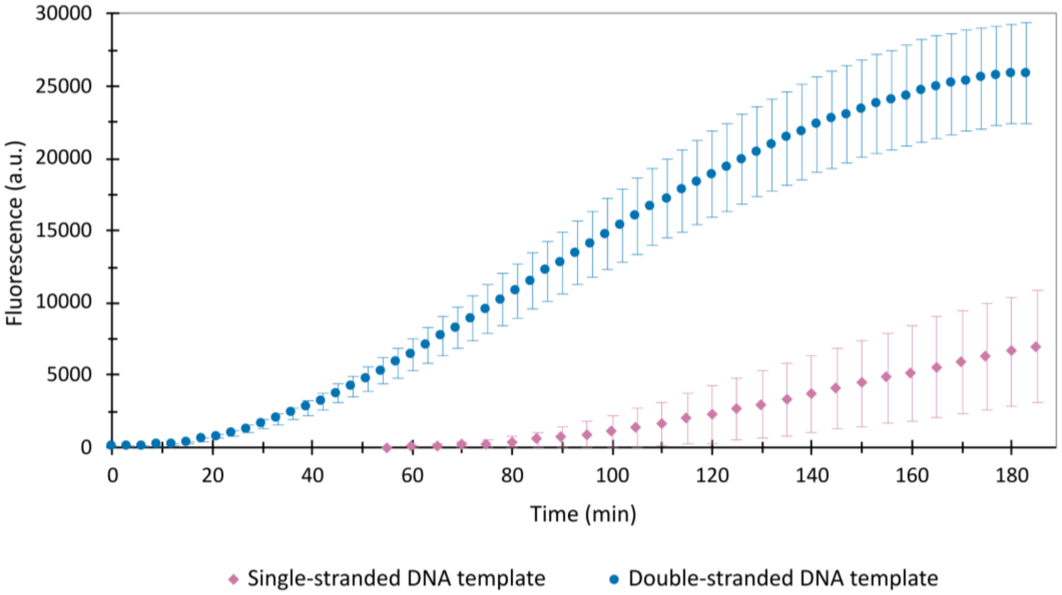

**Supplementary Figure 9. Validation of the dual-FLAP termination assay.** A Mango III–T1–Broccoli template was used to validate simultaneous detection of the upstream and downstream fluorescent RNA reporters. In the original Chen et al.[9] protein-reporter assay, T1 produced the same fluorescence output from the upstream and downstream protein reporters, resulting in a termination strength of 1. Thus, T1 represents a terminator sequence expected to permit extensive transcriptional readthrough. Consistent with this expectation, fluorescence accumulation was observed in both the upstream Mango III channel (a), which provides a transcriptional reference, and the downstream Broccoli channel (b), which reports downstream readthrough. More efficient terminators are expected to reduce the downstream Broccoli signal while leaving the Mango III signal comparatively unchanged.

a)

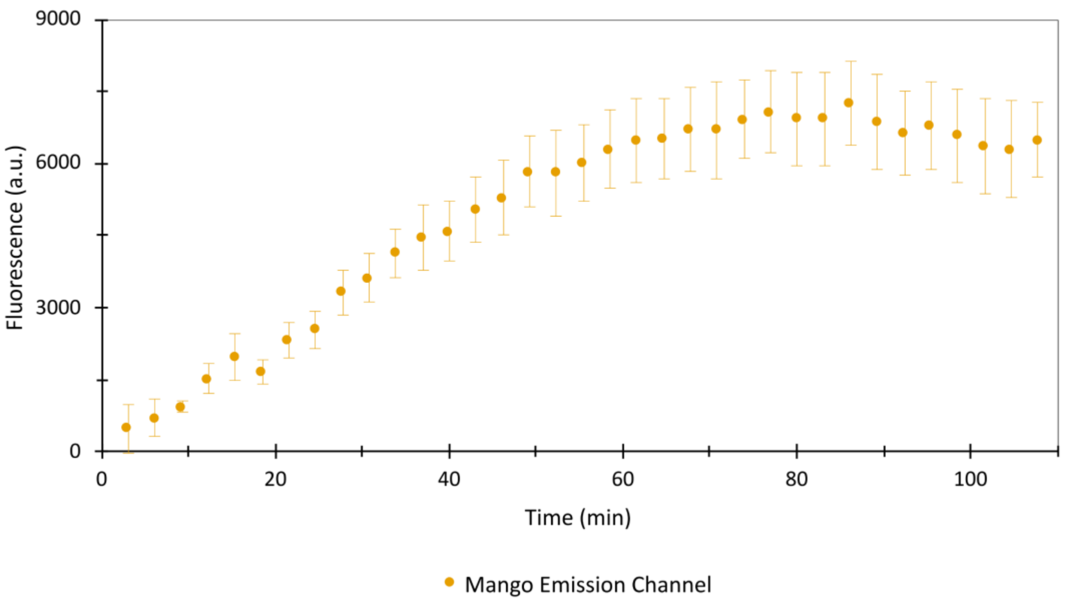


b)

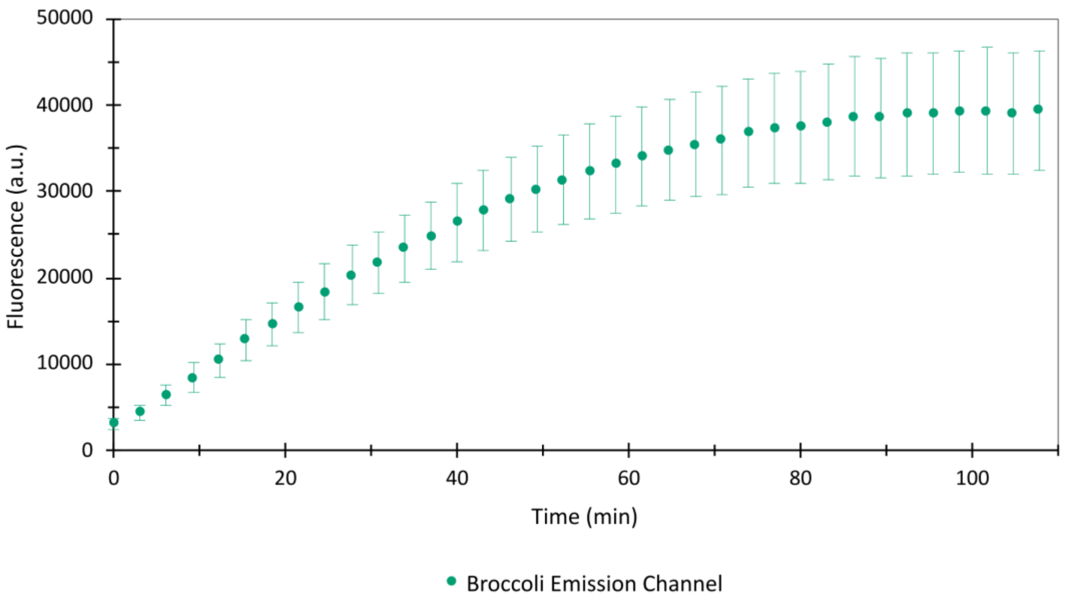

**Supplementary Figure 10. Reporter orientation in the dual-FLAP in vitro transcription assay.** For the assay to resolve differences among strong terminators, the downstream reporter must produce a detectable readthrough-dependent signal even when termination efficiency is high. We therefore evaluated whether the inverse orientation, Broccoli–terminator–Mango III, was suitable for termination measurements by placing T41 upstream of Mango III. T41 was the weakest terminator among those classified as strong in the Chen dataset. In this configuration, Broccoli produced a robust upstream fluorescence signal (a), indicating efficient transcription from the template. By contrast, the downstream Mango III signal remained close to background (b). This indicated that the Broccoli–terminator–Mango orientation would not provide sufficient dynamic range to discriminate among strong terminators, as stronger terminators would be expected to further reduce Mango III output. Therefore, the opposite orientation, Mango–terminator–Broccoli, was used in the final assay, enabling the differentiation of terminator strengths shown in Figure 2e.

a)

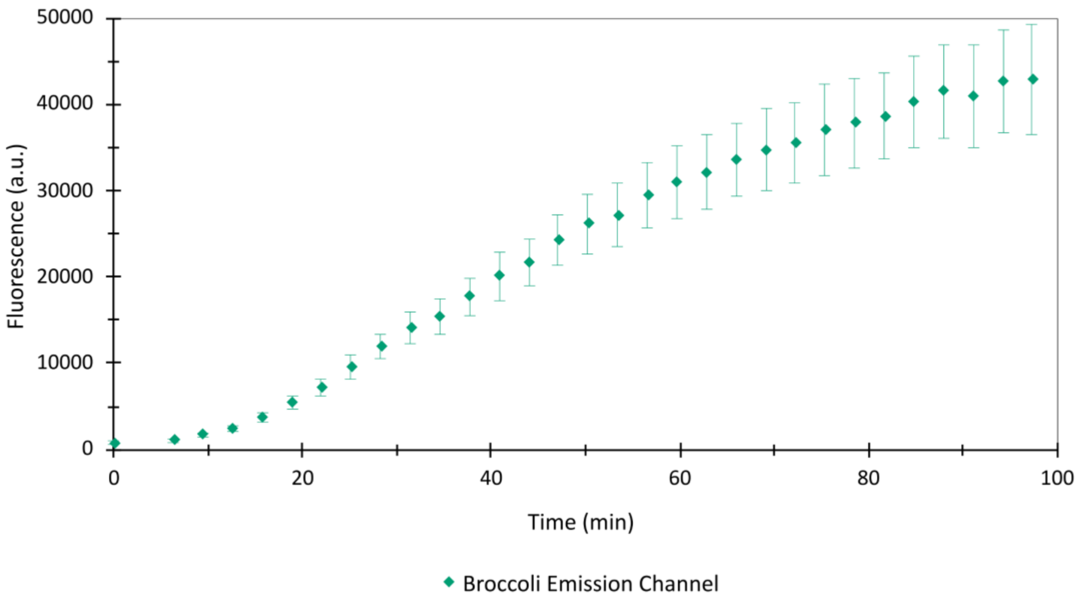


b)

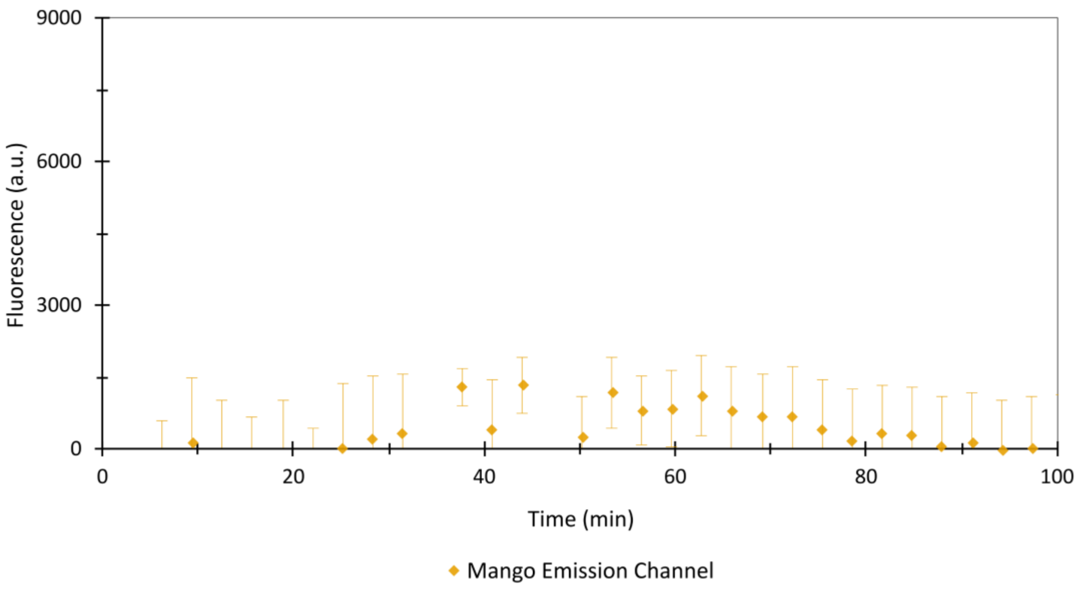

**Supplementary Figure 11. Effect of rNTP concentration on readthrough detection in the dual-FLAP termination assay.** The downstream Broccoli signal was monitored for terminators with different strengths under two rNTP concentrations. Commercial *in vitro* transcription kits are optimized for high RNA yield and typically use elevated rNTP concentrations. At 5 mM rNTPs, substantial downstream fluorescence was observed even for Tmax, indicating high readthrough-associated signal under these conditions. Reducing the rNTP concentration to 0.3 mM decreased this background readthrough and improved the dynamic range of the assay, allowing clearer discrimination between the weak terminator T1, the threshold-strength terminator T41, and the very strong terminator Tmax. The reduction in readthrough at lower rNTP concentration has been described previously[31] and is consistent with the long-standing model that intrinsic termination reflects kinetic competition between elongation through the terminator and transcript release. Note that the y scales are different for each graph

a)

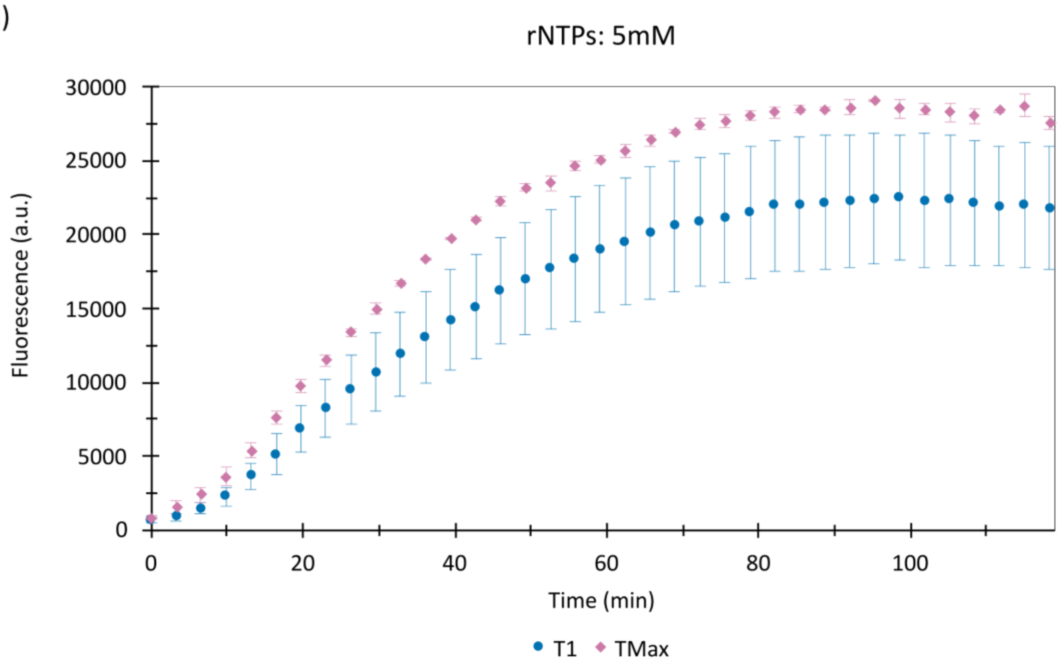


b)

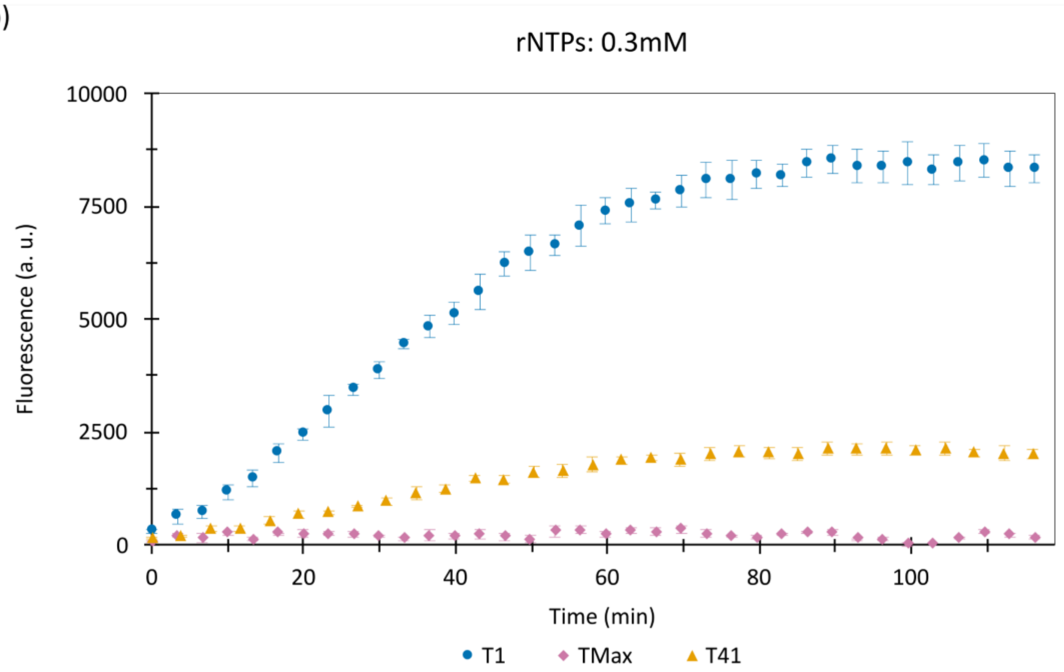

**Supplementary Figure 12.** Development of TerSP and TerFac. Parallelograms indicate inputs and outputs, rectangles represent computational steps, and rounded rectangles denote the start and end points. Arrows indicate the direction of data flow across modules. The uncolored region outlines the model development process, including feature engineering, selection, and training. Colored areas correspond to the major components of the deployed system: TerSP Web (blue), TerSP offline (green), feature mapping (purple), TerFac Web (yellow), and TerFac offline (red).

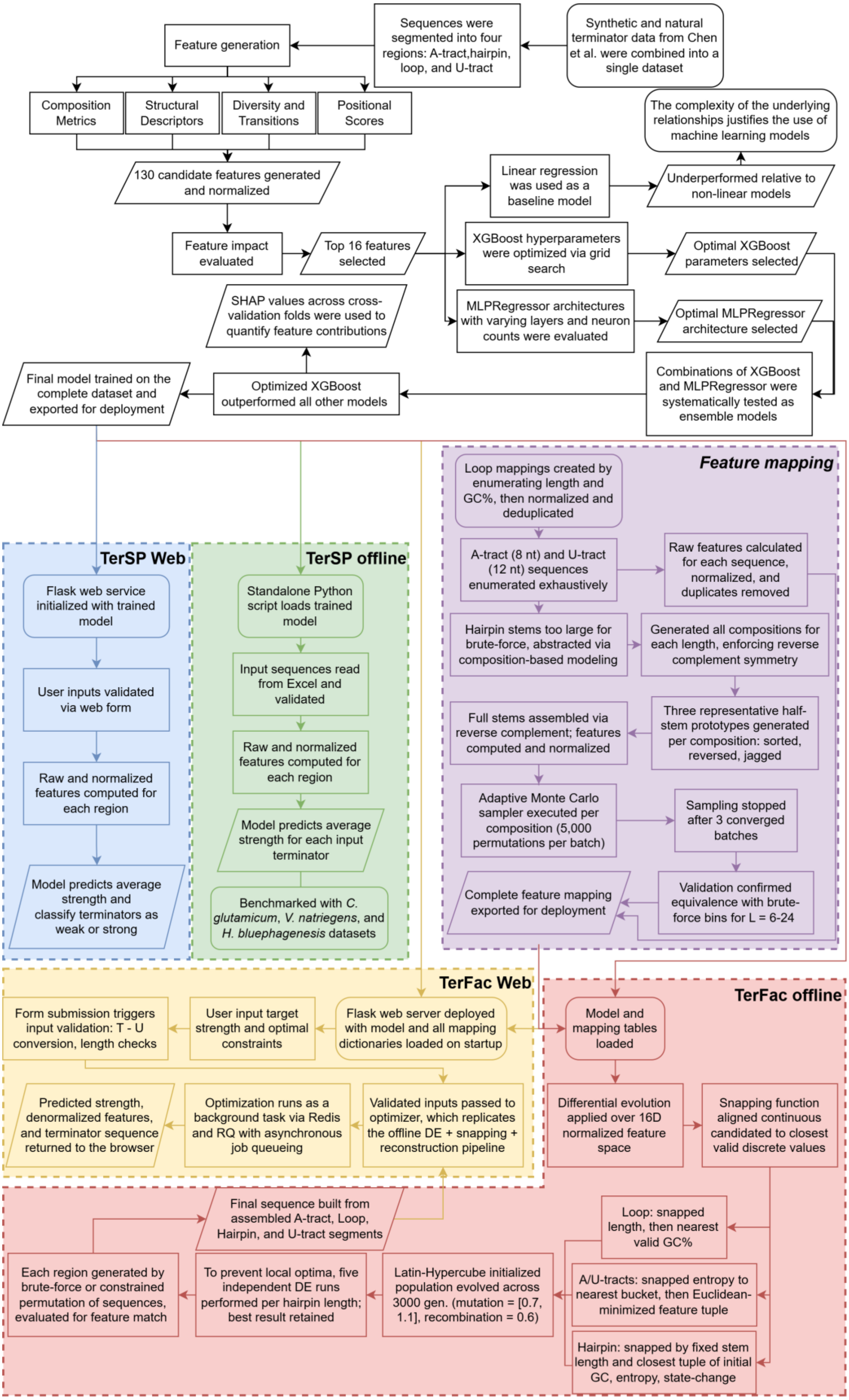
Synthetic and natural terminator data from Chen et al. were combined into a single dataset
Sequences were segmented into four regions: A-tract,hairpin, loop, and U-tract
Feature generation
Composition Metrics
Structural Descriptors
Diversity and Transitions
Positional Scores
130 candidate features generated and normalized
Feature impact evaluated
Top 16 features selected
SHAP values across cross-validation folds were used to quantify feature contributions
Linear regression was used as a baseline model
Underperformed relative to non-linear models
The complexity of the underlying relationships justifies the use of machine learning models
XGBoost hyperparameters were optimized via grid search
Optimal XGBoost parameters selected
MLPRegressor architectures with varying layers and neuron counts were evaluated
Optimal MLPRegressor architecture selected
Combinations of XGBoost and MLPRegressor were systematically tested as ensemble models
Optimized XGBoost outperformed all other models
Final model trained on the complete dataset and exported for deployment
TerSP Web
Flask web service initialized with trained model
User inputs validated via web form
Raw and normalized features computed for each region
Model predicts average strength and classify terminators as weak or strong
TerSP offline
Standalone Python script loads trained model
Input sequences read from Excel and validated
Raw and normalized features computed for each region
Model predicts average strength for each input terminator
Benchmarked with C. glutamicum, V. natriegens, and H. bluephagenesis datasets
Feature mapping
Loop mappings created by enumerating length and GC%, then normalized and deduplicated
A-tract (8 nt) and U-tract (12 nt) sequences enumerated exhaustively
Raw features calculated for each sequence, normalized, and duplicates removed
Hairpin stems too large for brute-force, abstracted via composition-based modeling
Generated all compositions for each length, enforcing reverse complement symmetry
Three representative half-stem prototypes generated per composition: sorted, reversed, jagged
Full stems assembled via reverse complement; features computed and normalized
Adaptive Monte Carlo sampler executed per composition (5,000 permutations per batch)
Sampling stopped after 3 converged batches
Validation confirmed equivalence with brute-force bins for L = 6-24
Complete feature mapping exported for deployment
TerFac Web
Form submission triggers input validation: T - U conversion, length checks
User input target strength and optimal constraints
Flask web server deployed with model and all mapping dictionaries loaded on startup
Predicted strength, denormalized features, and terminator sequence returned to the browser
Optimization runs as a background task via Redis and RQ with asynchronous job queueing
Validated inputs passed to optimizer, which replicates the offline DE + snapping + reconstruction pipeline
TerFac offline
Model and mapping tables loaded
Differential evolution applied over 16D normalized feature space
Snapping function aligned continuous candidates to closest valid discrete values
Loop: snapped length, then nearest valid GC%
A/U-tracts: snapped entropy to nearest bucket, then Euclidean-minimized feature tuple
Hairpin: snapped by fixed stem length and closest tuple of initial GC, entropy, state-change
Latin-Hypercube initialized population evolved across 3000 gen. (mutation = [0.7, 1.1], recombination = 0.6)
To prevent local optima, five independent DE runs performed per hairpin length; best result retained
Each region generated by brute-force or constrained permutation of sequences, evaluated for feature match
Final sequence built from assembled A-tract, Loop, Hairpin, and U-tract segments

### Protocol for *in vitro* terminator strength quantification reaction

#### Reagents

- Optimized **5× Transcription Buffer (Promega)**: 40 mM Tris-HCl (pH 7.9), 6 mM $MgCl_2$, 2 mM spermidine, and 10 mM NaCl (after dilution).
- RNase-free Potassium chloride (**KCl**, DEPC-treated), **4 M**
- RNase-free Magnesium chloride (**$MgCl_2$**, DEPC-treated), **1 M**
- Dithiothreitol (**DTT**), **100 mM**
- Ribonucleoside triphosphates (**rNTPs**), **1 mM**
- RNase-free **water**
- (Z)-3-((1H-benzo[d]imidazol-4-yl)methyl)-5-(3,5-difluoro-4-hydroxybenzylidene)-2-methyl-3,5-dihydro-4H-imidazol-4-one (**BI**), **2.5 mM**
- 1-methyl-2-[(3-methyl-3H-benzothiazol-2-ylidene)methyl]quinolinium iodide (**TO1**), **400 µM**
- **T7 RNA polymerase** (Promega)
- Inorganic pyrophosphatase (**PPase**) (Thermo Fisher)
- **DNA template**

#### Reaction setup

- At room temperature, prepare a master mix containing, per reaction:
  2 µL 5x Transcription Buffer, 0.38 µL KCl, 0.3 µL $MgCl_2$, and 1 µL DTT.
- Dispense 3.68 µL of this master mix into a PCR tube for each reaction.
- Separately, dilute 50 ng of DNA template in RNase-free water to a final volume of 3.62 µL for a 10 µL reaction.
- Add the diluted DNA to the reagents from the first step.
- Transfer the PCR tubes to ice.
- Add 1.5 µL of rNTPs to each tube.
- Prepare a fluorophore mix containing, per reaction, 0.25 µL BI and 0.25 µL TO1.
- Protect the tubes from light.
- Add 0.5 µL of the fluorophore mix to each tube.
- Add 0.2 µL of PPase to each tube.
- Add 0.5 µL of T7 RNA polymerase to each tube.
- Transfer the contents of the tubes to the plate used for fluorescence measurement and start the quantification on the plate reader.

*The DNA should not be added to the buffer while on ice, because spermidine can cause DNA precipitation at low temperatures. PPase is not essential, but it facilitates signal acquisition by reducing the risk of inorganic phosphate accumulation.*

### References


1. Brendel, V. and Trifonov, E.N. (1984) A computer algorithm for testing potential prokaryotic terminators. *Nucleic Acids Res.* 12, 4411–4427

2. Brendel, V. *et al.* (1986) Terminators of transcription with RNA polymerase from Escherichia coli: what they look like and how to find them. *J. Biomol. Struct. Dyn.* 3, 705–723

3. Carafa, Y. d'Aubenton *et al.* (1990) Prediction of rho-independent Escherichia coli transcription terminators: a statistical analysis of their RNA stem-loop structures. *J. Mol. Biol.* 216, 835–858

4. Ermolaeva, M.D. *et al.* (2000) Prediction of transcription terminators in bacterial genomes. *J. Mol. Biol.* 301, 27–33

5. Lesnik, E.A. *et al.* (2001) Prediction of rho-independent transcriptional terminators in Escherichia coli. *Nucleic Acids Res.* 29, 3583–3594

6. de Hoon, M.J.L. *et al.* (2005) Prediction of transcriptional terminators in Bacillus subtilis and related species. *PLoS Comput. Biol.* 1, e25

7. Kingsford, C.L. *et al.* (2007) Rapid, accurate, computational discovery of Rho-independent transcription terminators illuminates their relationship to DNA uptake. *Genome Biol.* 8, R22

8. Cambray, G. *et al.* (2013) Measurement and modeling of intrinsic transcription terminators. *Nucleic Acids Res.* 41, 5139–5148

9. Chen, Y.-J. *et al.* (2013) Characterization of 582 natural and synthetic terminators and quantification of their design constraints. *Nat. Methods* 10, 659–664

10. Cui, W. *et al.* (2021) Data-driven and in silico-assisted design of broad host-range minimal intrinsic terminators adapted for bacteria. *ACS Synth. Biol.* 10, 1438–1450

11. Xu, M. *et al.* (2022) Development and application of transcription terminators for polyhydroxylkanoates production in halophilic Halomonas bluephagenesis TD01. *Front. Microbiol.* 13, 941306

12. Zhai, W. *et al.* (2022) Sequence and thermodynamic characteristics of terminators revealed by FlowSeq and the discrimination of terminators strength. *Synth. Syst. Biotechnol.* 7, 1046–1055

13. Li, J. *et al.* (2025) Intelligent Design of Escherichia coli Terminators by Coupling Prediction and Generation Models. *ACS Synth. Biol.* 14, 3744–3752

14. Gardner, P.P. *et al.* (2011) RNIE: genome-wide prediction of bacterial intrinsic terminators. *Nucleic Acids Res.* 39, 5845–5852

15. Feng, C.-Q. *et al.* (2019) iTerm-PseKNC: a sequence-based tool for predicting bacterial transcriptional terminators. *Bioinformatics* 35, 1469–1477

16. Brandenburg, V.B. *et al.* (2022) Inverse folding based pre-training for the reliable identification of intrinsic transcription terminators. *PLoS Comput. Biol.* 18, e1010240

17. Taheri Ghahfarokhi, S.M.A. and Peña-Castillo, L. (2025) BacTermFinder: a comprehensive and general bacterial terminator finder using a CNN ensemble. *NAR Genomics*

*Bioinforma.* 7, lqaf016

18. Tjaden, B. (2026) TerminatorNet: comprehensive identification of intrinsic transcription terminators in bacteria. *Bioinformatics* 42, btag116

19. Wilson, K.S. and von Hippel, P.H. (1995) Transcription termination at intrinsic terminators: the role of the RNA hairpin. *Proc. Natl. Acad. Sci.* 92, 8793–8797

20. Lubkowska, L. *et al.* (2011) RNA folding in transcription elongation complex: implication for transcription termination. *J. Biol. Chem.* 286, 31576–31585

21. Lynn, S.P. *et al.* (1988) Contributions of RNA secondary structure and length of the thymidine tract to transcription termination at the thr operon attenuator. *J. Biol. Chem.* 263, 472–479

22. Sipos, K. *et al.* (2007) Systematic mutagenesis of the thymidine tract of the pyrBI attenuator and its effects on intrinsic transcription termination in Escherichia coli. *Mol. Microbiol.* 66, 127–138

23. Telesnitsky, A. and Chamberlin, M.J. (1989) Terminator-distal sequences determine the in vitro efficiency of the early terminators of bacteriophages T3 and T7. *Biochemistry* 28, 5210–5218

24. Reynolds, R. and Chamberlin, M.J. (1992) Parameters affecting transcription termination by Escherichia coli RNA: II. Construction and analysis of hybrid terminators. *J. Mol. Biol.* 224, 53–63

25. Holmes, W.M. *et al.* (1983) Termination of transcription in E. coli. *Cell* 32, 1029–1032

26. Postle, K. and Good, R.F. (1985) A bidirectional rho-independent transcription terminator between the E. coli tonB gene and an opposing gene. *Cell* 41, 577–585

27. Brosius, J. (1984) Toxicity of an overproduced foreign gene product in Escherichia coli and its use in plasmid vectors for the selection of transcription terminators. *Gene* 27, 161–172

28. Kang, D.H. *et al.* (2022) RoboMoClo: a robotics-assisted modular cloning framework for multiple gene assembly in biofoundry. *ACS Synth. Biol.* 11, 1336–1348

29. Stukenberg, D. *et al.* (2021) The Marburg Collection: A Golden Gate DNA assembly framework for synthetic biology applications in Vibrio natriegens. *ACS Synth. Biol.* 10, 1904–1919

30. Kartje, Z.J. *et al.* (2021) Revisiting T7 RNA polymerase transcription in vitro with the Broccoli RNA aptamer as a simplified real-time fluorescent reporter. *J. Biol. Chem.* 296, 100175

31. McDowell, J.C. *et al.* (1994) Determination of intrinsic transcription termination efficiency by RNA polymerase elongation rate. *Science* 266, 822–825
